# C-terminal amides mark proteins for degradation via SCF/FBXO31

**DOI:** 10.1101/2023.06.29.547030

**Authors:** Matthias Muhar, Jakob Farnung, Raphael Hofmann, Martina Cernakova, Nikolaos D. Sidiropoulos, Jeffrey W. Bode, Jacob E. Corn

## Abstract

During normal cellular homeostasis unfolded and mis-localized proteins are recognized and removed, preventing the build-up of toxic byproducts^1^. When protein homeostasis is perturbed during aging, neurodegeneration or cellular stress, proteins can accumulate several forms of chemical damage through reactive metabolites^2, 3^. Such modifications have been proposed to trigger the selective removal of chemically marked proteins^3–6;^ however, discovering modifications sufficient to induce protein degradation has remained challenging. Using a semi-synthetic chemical biology approach coupled to cellular assays, we found that C-terminal amide-bearing proteins (CTAPs) are rapidly cleared from human cells. A CRISPR screen identified the SCF/FBXO31 ubiquitin ligase as a reader of C-terminal amides, which ubiquitylates CTAPs for subsequent proteasomal degradation. A conserved binding pocket enables FBXO31 to bind almost any C-terminal peptide bearing an amide while retaining exquisite selectivity over non-modified clients. This mechanism facilitates binding and turnover of endogenous CTAPs that are formed following oxidative stress. A dominant human mutation found in neurodevelopmental disorders switches CTAP recognition, such that non-amidated neosubstrates are now degraded and FBXO31 becomes markedly toxic. We propose that CTAPs may represent the vanguard of a largely unexplored class of modified amino acid degrons that could provide a general strategy for selective yet broad surveillance of chemically damaged proteins.

## Main

Cellular protein homeostasis is the essential process of regulating the biogenesis, localization and turnover of proteins. Selective degradation of proteins by the ubiquitin-proteasome system is a central effector mechanism in protein homeostasis^7^. At the molecular level, degradation is initiated by post-translational modifications (PTMs), such as Lys48-linked poly-ubiquitylation, that mark proteins for further processing by downstream effectors. These modifications induce recruitment and activation of the proteasome for processive proteolysis. The specificity of this system is established by over 600 human ubiquitin ligases that can bind specific interaction motifs – termed degrons – on their respective client proteins^8^.

While many well-studied PTMs are deposited or erased by dedicated enzymes, amino acid side chains and the protein backbone itself can also experience a plethora of non-enzymatic modifications, including oxidative damage or alkylation^2^. In total, over 300 protein modifications have been described to-date^9^. For the majority of these, their role in protein homeostasis remains unexplored, in part due to a lack of scalable experimental methods. However, non-canonical PTMs accumulate following proteasome inhibition^5^ and treatment with alkylating or oxidating agents can stimulate protein turnover^10–12^. Particular chemical modifications have therefore long been postulated as marks for protein damage to trigger protein clearance^3^. However, specific modifications sufficient to induce degradation and the mechanisms by which they are recognized remain largely unknown.

### C-terminal amides mark proteins for degradation by the ubiquitin proteasome system

We sought a reductionist approach to discover the effect of individual chemical modifications of proteins on their turnover. To this end, we created a set of semi-synthetic fluorescent reporters and introduced them into human cells to measure their stability (**Fig. 1a and Extended Data Fig. 1a**).

**Fig 1.**
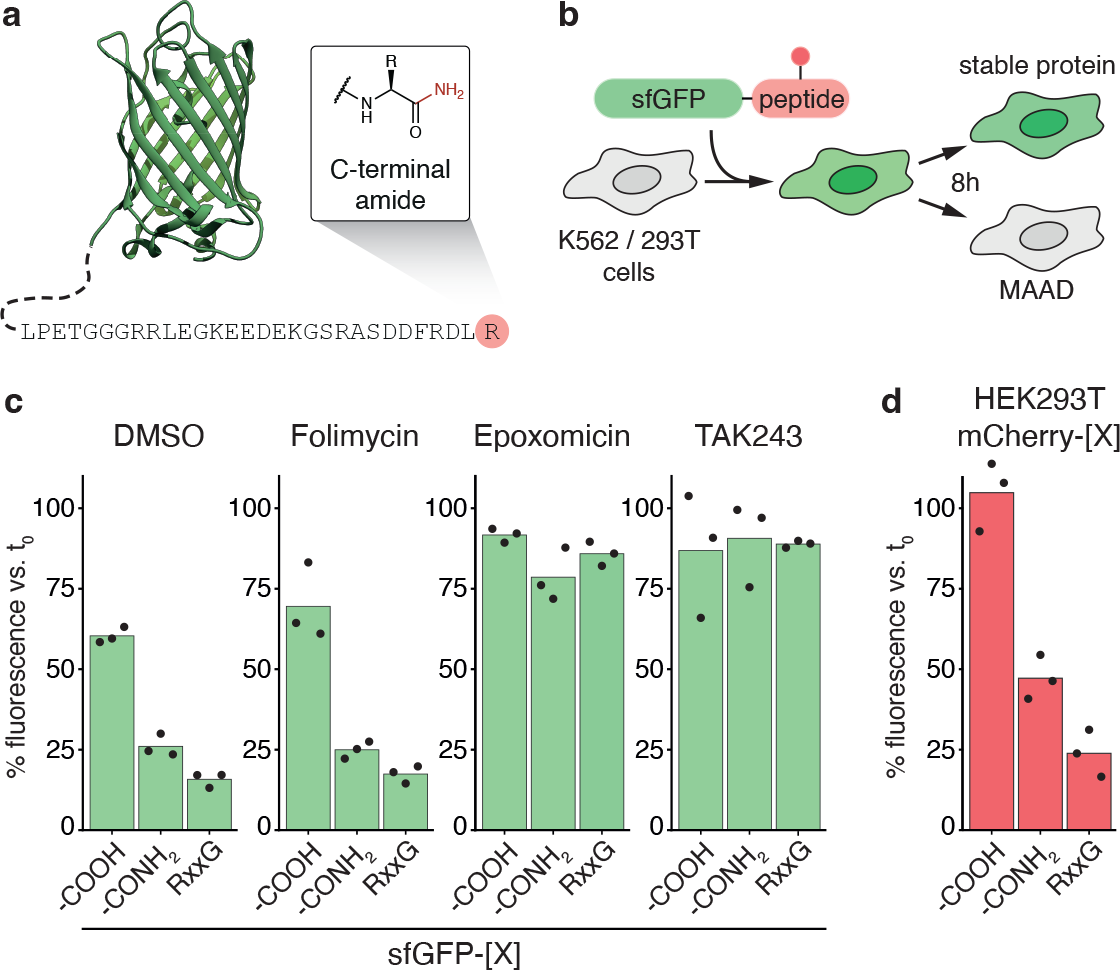
Proteins with C-terminal amides are selectively degraded by the ubiquitin proteasome system. (**a**) Schematic of an sfGFP-based semi-synthetic reporter protein carrying a C-terminal amide. (**b**) Schematic of a fluorescent in-cell reporter assay to distinguish modified amino acid degrons (MAADs) from neutral modifications by electroporation of reporter proteins into human cell lines. (**c**) Results of an in-cell reporter assay for the sfGFP conjugate shown in (a) with (-CONH2) or without (-COOH) terminal amide. RxxG denotes an sfGFP variant containing a positive control degron motif (LPETGGGRRLEGKEEDEKGSRASDRFRGLR). K562 cells received mock-treatment (DMSO), lysosomal inhibitor folimycin (100 nM), proteasome inhibitor epoxomicin (500 nM) or E1 ubiquitin ligase inhibitor TAK243 (1 µM) upon sfGFP delivery. (**d**) Reporter assay as in (c) in HEK293T cells using the same peptides conjugated to mCherry. Full protein sequences are listed in Supplementary Table 1. Bars represent means of three experiments. Black dots indicate individual measurements.

Using solid phase peptide synthesis, we generated a set of peptides carrying defined modifications that represent different types of protein damage: tyrosine modification through oxidation or misincorporation (L-3,4-dihydroxyphenylalanine, L-DOPA)^13^, carbonylation (N(6)-hexanoyllysine)^14^, advanced glycation end products (N(6)-carboxymethyllysine)^15^, carbamylation (homocitrulline)^4^ and primary amide-forming backbone cleavage (C-terminal amide)^16, 17^. Sortase A-mediated conjugation of these peptides onto the C-terminus of recombinant fluorescent proteins yielded a set of chemically defined modified reporters for further study (**Extended Data Fig. 1b, c, Extended Data Table 1).**

We measured the effect of each modification on protein degradation in the human erythroleukemia cell line K562 with a fluorescent in-cell reporter assay. We introduced individual semi-synthetic proteins by electroporation into human cells and followed their clearance over time by flow cytometry (**Fig. 1b**). In this setting, unconjugated superfolder GFP (sfGFP-SRT-H_6_) alone was highly stable (*t*_1/2_

> 16h), while introduction of a strong degron sequence derived from the human ASCC3 C-terminus (RxxG)^18^ induced rapid degradation (**Extended Data Fig. 1d**). None of the internal modifications tested affected protein stability in this cell line and amino acid context, suggesting that they are not universally sufficient to induce protein degradation (**Extended Data Fig. 1e**).

In contrast, sfGFP carrying a primary amide on its C-terminus was rapidly degraded in two independent sequence contexts (**Fig. 1c, Extended Data Fig. 1e**). Amidated sfGFP degradation was prevented by inhibitors of total cellular ubiquitylation (TAK243) or the proteasome (epoxomicin), but not by inhibition of lysosomal acidification (folimycin). This implies that C-terminal amide-bearing proteins (henceforth CTAPs) are actively cleared from cells by the ubiquitin proteasome system. CTAP forms of both mCherry and mTagBFP2, but not their unmodified counterparts, were also efficiently degraded when delivered to human embryonic kidney-derived HEK293T cells (**Fig. 1d and Extended Data Fig. 1f**). The presence of a C-terminal amide is therefore sufficient to induce protein degradation in different peptide-, protein- and cell contexts.

### SCF/FBXO31 mediates CTAP clearance

To identify the cellular machinery underlying the recognition and removal of CTAPs, we devised a genome-wide CRISPR screen for genes responsible for specific degradation of C-terminally amidated sfGFP (sfGFP-CONH_2_, **Fig. 2a**). Since knockout of central protein quality control and -turnover genes may impede cell survival, we generated a clonal K562 cell line with a tightly controllable Cas9 allele (iCas9) (**Extended Data Fig. 2a, b**). Following transduction with the TKOv3 genome-wide sgRNA library^19^, Cas9 expression was induced for 5 days to allow for efficient knockout while minimizing loss of cells with defects in essential pathways^20^. We then delivered sfGFP-CONH_2_ by electroporation together with mTagBFP2-RxxG, an internal control for general protein turnover carrying a sequence-based degron. Following short-term culture allowing for protein degradation (14 h), we isolated cells that were proficient in general protein turnover but deficient in CTAP clearance (sfGFP+ mTagBFP2-), as well as a control population with no defect in protein clearance (sfGFP-mTagBFP2-). Next generation sequencing and quantification of sgRNAs from both populations identified gene knockouts enriched among CTAP-positive cells, which are candidate CTAP clearance factors.

**Fig 2.**
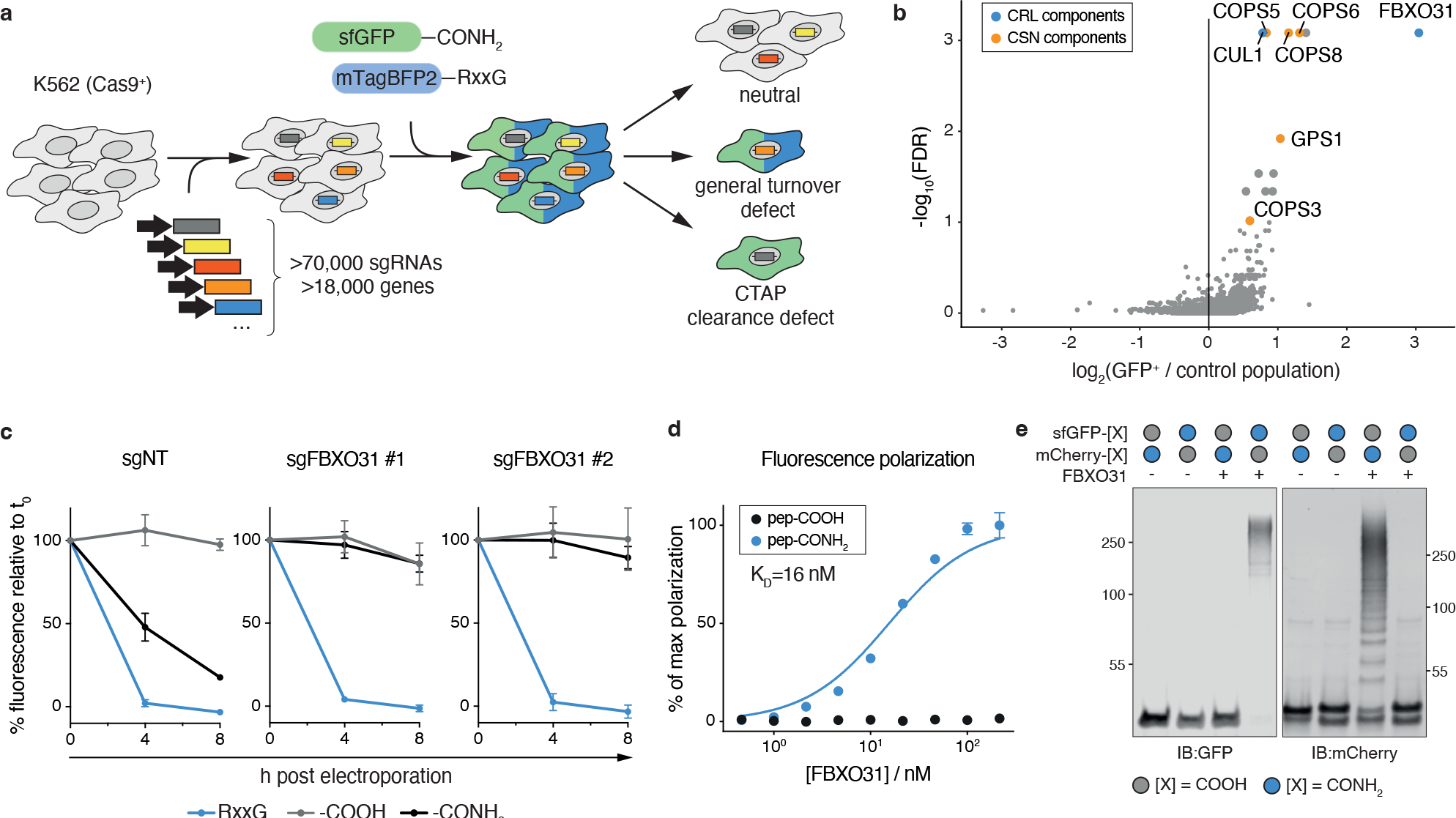
CRISPR screen identifies SCF/FBXO31 as a CTAP clearance factor. (**a**) Schematic of a genome-wide CRISPR screen to identify genes required for CTAP clearance. Model substrates were sfGFP-GGGKDLEGKGGSAGSGSAGGSKYPYDVPDYAKS-CONH2 (sfGFP-pep1-CONH2) and mTagBFP2-GGGRRLEGKEEDEKGSRASDRFRGLR-COOH (pep2-RxxG). (**b**) Screening results showing mean enrichment of sgRNAs targeting each gene in CTAP-clearance deficient cells (sfGFP+ mTagBFP2-) compared to the control population (sfGFP-mTagBFP2-) on the abscissa. Ordinate values depict FDR-adjusted significance of enrichment across sgRNAs and duplicate screens. (**c**) Time-course of sfGFP degradation in CRISPRi competent K562 cells expressing sgRNAs targeting the transcription start site of FBXO31 and a non-targeting control (NT). sfGFP was conjugated to pep1 or a positive control degron RxxG as in (b). Error bars indicate the standard deviation of three independent experiments. (**d**) Fluorescence polarization (FP) assay of fluorescently labeled peptide (pep2-short, KEEDEKGSRASDDFRDLR) with indicated C-termini. Error bars indicate the standard deviation of three separate FP reactions. (**e**) *In vitro* ubiquitylation assay using sfGFP-pep-1 and mCherry-pep2 with indicated C-termini, recombinant FBXO31, remaining E3 complex members (SKP1, CUL1-NEDD8, RBX1), E1 (UBA1) and E2 (UBE2R1, UBE2D3) enzymes.

Bioinformatic analysis (Methods) revealed a stark enrichment of few targeted genes in the CTAP clearance-deficient population. The most prominent hit was FBXO31 (**Fig. 2b, Supplementary Table 1**), which is a substrate adaptor for the SCF (SKP1-CUL1-F-box protein) E3 ubiquitin ligase assembly. The screen also identified SCF’s central scaffolding protein CUL1 and several subunits of the COP9 signalosome complex, which is required for SCF function^21^ (**Extended Data Fig. 2c**). Members of the SCF and the COP9 signalosome are essential genes^22^, but were identified due to the use of the iCas9 system. FBXO31 is non-essential under basal conditions, as might be expected for a factor acting in response to cellular stress.

Individually silencing FBXO31 by CRISPR inhibition (CRISPRi)^23^ using two independent guide RNAs in K562 cells completely stabilized the CTAP-form of the GFP reporter, while the RxxG degron form remained unaffected (**Fig. 2c**). FBXO31 knockout in the human embryonic kidney derived cell line HEK293 also abolished CTAP clearance (**Extended Data Fig. 2d**). Overexpression of wildtype FBXO31 rescued CTAP degradation in the knockout background, but a mutant of FBXO31 lacking the F-box domain required for SCF assembly did not (**Extended Data Fig. 2d**). These findings establish SCF/FBXO31 as a central effector of CTAP clearance in the context of both multiple model substrates and multiple cellular backgrounds.

We mechanistically investigated the role of SCF/FBXO31 in CTAP clearance using multiple orthogonal assays. We first tested whether SCF/FBXO31 directly binds and ubiquitylates amidated clients, or whether it plays an indirect role in CTAP clearance. We purified recombinant FBXO31 in complex with its binding partner SKP1 and measured its affinity for various peptides by fluorescence polarization (FP). *In vitro*, FBXO31 bound the peptide-amide used for screening with high affinity (K_D_ = 16 ± 2 nM) while no binding could be detected for the carboxylic acid form (**Fig. 2d**). To test whether FBXO31’s interaction with peptide amides also occurs in cells, we performed co-immunoprecipitation (co-IP) of affinity-tagged FBXO31, which exhibited binding exclusively to the model substrate mCherry-CONH_2_, but not the unmodified counterpart (**Extended Data Fig. 2e**).

To test whether FBXO31 binding leads to productive ubiquitylation of substrates, we reconstituted the full SCF/FBXO31 E3 ligase assembly from recombinant components. The SCF/FBXO31 complex ubiquitylated sfGFP-CONH_2_ *in vitro* with high efficiency but showed no detectable activity towards sfGFP-COOH (**Extended Data Fig. 2f)**. In reactions simultaneously containing both unmodified and amide-bearing substrates, SCF/FBXO31 exclusively ubiquitylated the CTAP modified protein. SCF/FBXO31 is therefore capable of precisely distinguishing amidated from non-amidated substrates without cross-talk (**Fig. 2e**). Together these results establish that SCF/FBXO31 is a reader of C-terminal protein amides, and that this amide is required for SCF/FBXO31 to ubiquitylate targets.

### FBXO31 broadly and selectively binds amidated C-termini

If FBXO31 is a general surveillance factor for C-terminal amides, it would need to sample a broad range of C-terminal sequences while maintaining high selectivity for amides over carboxylic acids. Moreover, it could not simply bind any primary amide group, as these are found in asparagine and glutamine side chains of virtually all proteins. We therefore set out to determine the substrate scope of FBXO31 using *in vitro* binding studies.

We first tested whether FBXO31 specifically binds C-terminal amides rather than side chain amides in asparagine (N) or glutamine (Q). FP assays using recombinant FBXO31/SKP1 and fluorescently labeled peptides showed no affinity for peptides with unmodified N or Q at the C-terminus (**Fig. 3a**). However, the same peptide sequences were bound with high affinity when carrying a C-terminal primary amide (X-N-CONH_2_: K_D_ = 24 ± 3 nM, X-Q-CONH_2_: K_D_ = 55 ± 4 nM). Extending this assay to peptides bearing the primary amide derivatives of each of the 20 natural amino acids revealed that FBXO31 binds virtually any C-terminal amide with nanomolar affinity (**Fig. 3b, Extended Data Fig. 3**). The weakest binders were peptides bearing glycine and acidic residues, with X-D-CONH_2_ exhibiting a K_D_ of 304 ± 22 nM. Hydrophobic residues were most strongly bound, with X-F-CONH_2_ binding even more strongly (K_D_ ≈ 6 nM) than the most potent peptide identified during our initial studies, followed by uncharged and charged hydrophilic side chains. These findings demonstrate that FBXO31 binds diverse C-terminal amides with high affinity and selectivity over unmodified C-termini and side chain amides.

**Fig 3.**
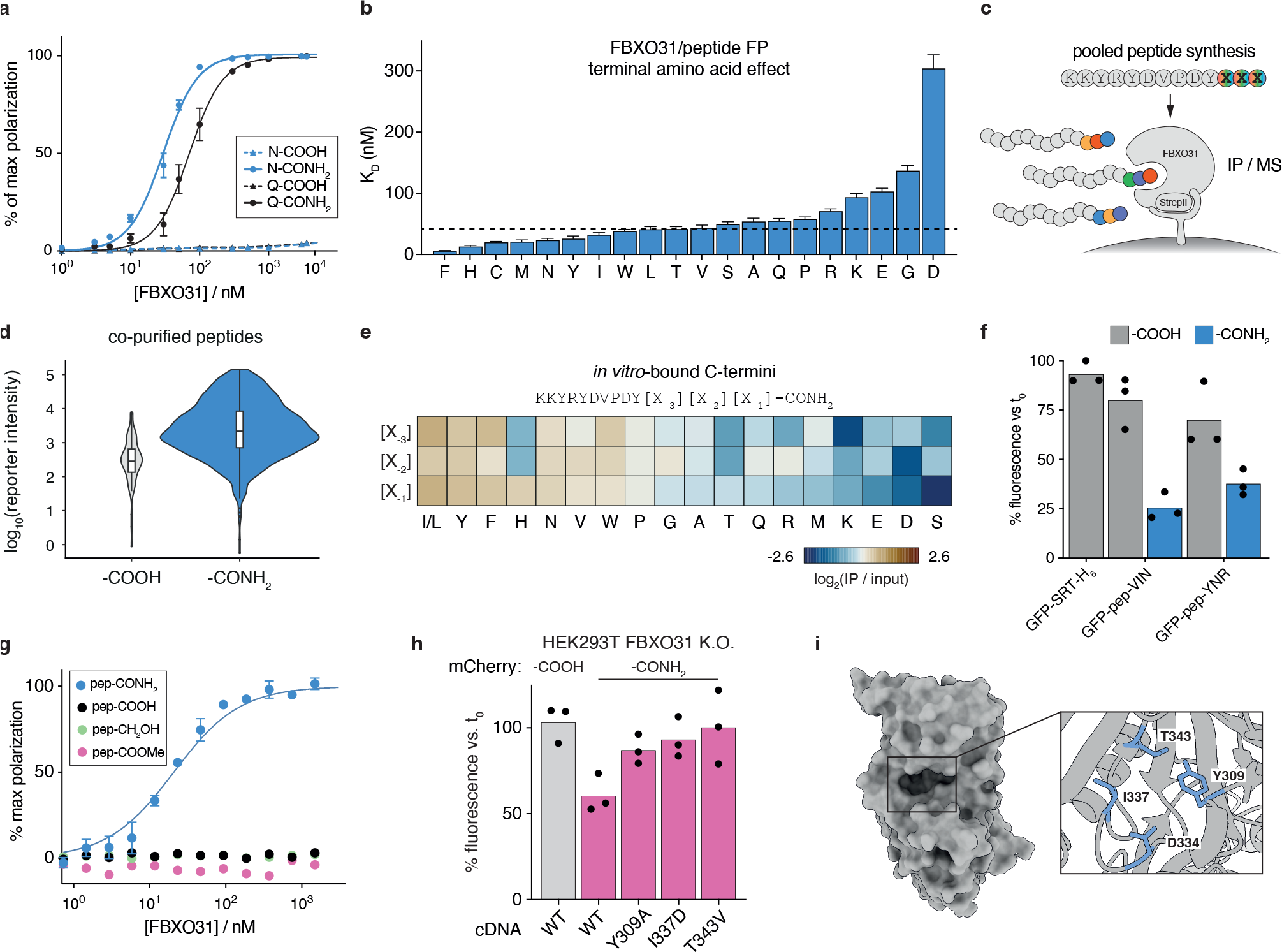
FBXO31 broadly and selectively binds CTAPs. (**a**) FP assay of FBXO31 binding pep3 (KKYRYDVPDYSA[X]) with indicated terminal amino acids and modifications. (**b**) KD for FBXO31 binding pep3-CONH2 as in (a) for different amino acids in the terminal position. The dashed line indicates the median across terminal amino acids. (**c**) Schematic of a pooled FBXO31 interaction screen for a library of unmodified and C-terminally amidated peptides. (**d**) Violin plot showing number and relative recovery of peptides bound by FBXO31 *in vitro* as in (d) for unmodified (-COOH) and amide-bearing (-CONH2) C-termini. Y-axis values depict total reporter ion intensities for identified peptides scaled to an isobarically labeled input library (see Methods). The median scaled intensity differs by 7.6-fold (p < 10^-17^, Wilcoxon rank-sum test). (**e**) Heatmap of amino acid frequencies among FBXO31-bound peptides identified in (d) relative to input. (**f**) In-cell protein stability assay for sfGFP conjugated to top-scoring amide-bearing C-termini bound by FBXO31 in (d). Reporter proteins were delivered by electroporation and quantified by flow cytometry after 0.5 (t0) and 8 hours. (**g**) FP assay of FBXO31 interaction with [fluorescein]-KEEDEKGSRASDDFRDLR for indicated C-terminal modifications. (**h**) In-cell reporter assay of a model CTAP (mCherry-pep2) with indicated C-terminal groups for FBXO31 knockout HEK293 cells stably expressing wt or mutant FBXO31 cDNAs. (**i**) Crystal structure of FBXO31 from PDB:5VZT. Inset shows zoom-in on its substrate binding pocket. All error bars indicate SD of three reactions. All bar graphs represent means of three experiments. Black dots indicate individual replicates in (f) and (h).

We extended the screen of amidated C-termini by devising a massively parallel protein-peptide interaction screen to identify sequence preferences for FBXO31 binding (**Fig. 3c**). Using isokinetic mixtures of 19 natural amino acids (all except cysteine) in the first three coupling steps, we synthesized peptide libraries containing >2’000 distinct C-termini detectable by mass spectrometry (MS) (**Extended Data Fig. 4a, Supplementary Table 2**). To quantify FBXO31/SKP1 binding to these sequences, we performed *in vitro* co-immunoprecipitation of the library and quantified the abundance of each peptide alongside the input pool using isobaric labeling and MS.

Overall, FBXO31 exhibited broad binding to 841 distinct C-terminal amides co-purified with FBXO31, compared to just 73 unmodified C-termini. The C-terminal amides showed 7.6-fold greater enrichment compared to the few unmodified peptides detected in this assay (**Fig. 3d**). Consistent with individual FP measurements, FBXO31 preferred hydrophobic side chains while acidic residues were less efficiently bound, especially in the terminal position (**Fig. 3e**). Despite these preferences, each tested amino acid was detected in all three ultimate positions among bound peptides (**Extended Data Fig. 4b, c**). To validate that these *in vitro* measurements also translate to CTAP-clearance in cells, we measured degradation for GFP conjugated to the two top-scoring motifs (VIN, YNR) in HEK293T cells. Both top FBXO31-binding motifs were degraded in the amide-bearing context in a cellular assay (**Fig. 3f**).

We overall conclude that FBXO31 is specific for peptide amidation with a slight preference for hydrophobic termini. However, unlike conventional sequence-based C-degrons, FBXO31 is mostly agnostic to specific sequence motifs, potentially enabling it to broadly surveil C-terminal amides across the diverse proteome.

### CTAPs are bound by a conserved pocket in FBXO31

CTAPs are characterized by the isosteric substitution of an oxygen atom with nitrogen and the associated loss of a negative charge under physiological conditions. We asked how FBXO31 exhibits such strong and selective binding to amide-bearing C-termini, as C-terminal sequence-based degrons are typically recognized through positive charges on the ubiquitin ligase that bind to the unmodified negatively charged C-terminus^24–27^. In contrast, a previous crystal structure of FBXO31 in complex with an unmodified C-terminal peptide from Cyclin D1 revealed a substrate binding pocket of FBXO31 that exhibits a negative charge at D334^28^ and might therefore disfavor unmodified C-termini (**Extended Data Fig. 5a, b**).

To test whether exclusion of a negatively charged C-terminus alone underlies CTAP selectivity, we measured FBXO31 binding of various modified peptides by FP. Replacing the negatively charged C-terminus (-COOH) of a peptide with an alcohol (-CH_2_OH) or blocking it with a methyl ester (-COOMe) was not sufficient to induce FBXO31 binding (**Fig. 3g**). These data suggest that a specific interaction with the terminal primary amide is required for binding.

To assess the contribution of residues in the FBXO31 binding pocket to C-terminal amide recognition, we overexpressed FBXO31 cDNAs in an FBXO31 knockout background and measured the impact of individual point mutations on CTAP clearance (**Fig. 3h, i**). Wildtype cDNA rescued FBXO31 knockout. In contrast, mutations in the binding pocket completely abolished CTAP clearance, either when mutating the hydrophobic residues in the binding pocket (Y309A and I337D) or conservatively replacing a putative hydrogen-bonding residue (T343V). Based on these and the above findings, we propose that the selectivity of FBXO31 for C-terminal amides arises from the combined charge exclusion of carboxylic acids and direct interaction with the primary amide of its clients.

### FBXO31 recognizes endogenous CTAPs formed under oxidative stress

To the best of our knowledge, C-terminal amidation has so far not been reported for intracellular proteins. However, enzymatic and non-enzymatic mechanisms for terminal amide formation have been described in other contexts. *In vivo*, most secreted peptide hormones are converted to primary amides by PAM (peptidyl glycine alpha-amidating monooxygenase) using copper-catalyzed hydroxyl radical attack and subsequent backbone cleavage to yield shortened C-terminally amidated peptides^17^. *In vitro*, hydroxyl radicals can trigger peptide bond cleavage by a similar mechanism, giving rise to C-terminal amide fragments (**Fig. 4a**)^29^. We therefore hypothesized that CTAP degradation via FBXO31 could serve to eliminate protein fragments following mis-trafficking of secretory proteins or oxidative damage.

**Fig 4.**
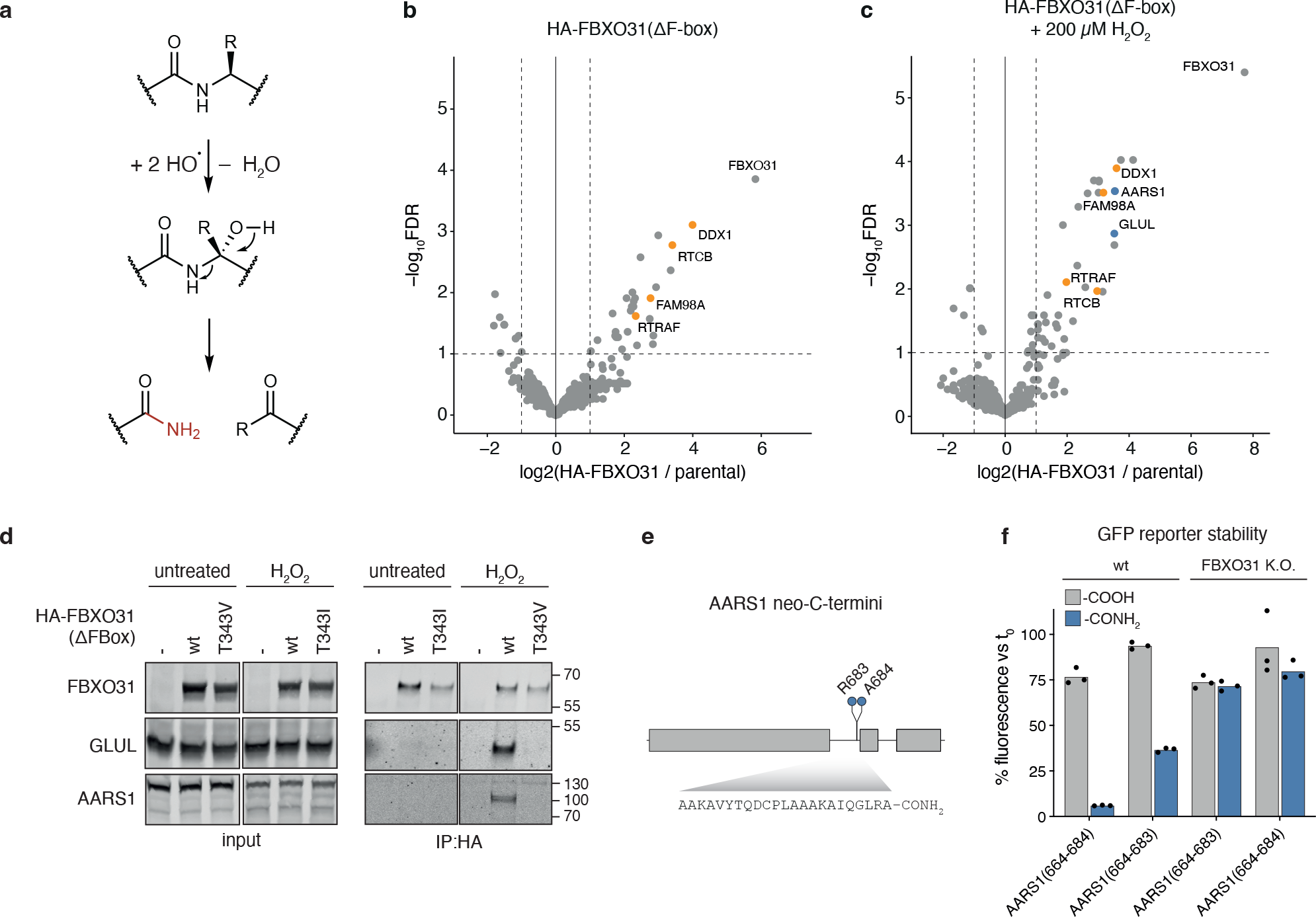
FBXO31 recognizes endogenous CTAPs following oxidative stress. (**a**) Proposed model of C-terminal amidation by PAM and hydroxyl radicals. (**b**) IP-MS of HA-tagged FBXO31(ΔF-box) from FBXO31 knockout HEK293T cells. Components of the tRNA ligase complex are shown in orange. (**c**) IP-MS as in (b) from cells treated for 20 min with 200 µM H2O2 in PBS containing 500 nM epoxomicin. Stress-specific interactors selected for follow-up are highlighted in blue, tRNA ligase complex components in orange. (**d**) Validation of FBXO31 clients by co-IP of cells treated as in (c). (**e**) Identification of CTAP cleavage sites by IP-MS of FLAG-AARS1 from FBXO31 knockout HEK293T cells treated as in (c). The map indicates putative cleavage sites along AARS1 in blue. The sequence below corresponds to a detected peptide with non-enzymatic C-terminus and terminal amide. (**f**) Validation of degron-activity for AARS1-derived CTAP-fragments. Indicated AARS1 residues were conjugated to sfGFP by sortylation. Protein levels were quantified by flow cytometry in wt and FBXO31 knockout HEK293T cells as in Fig. 1b, c. Bars represent means of three experiments. Black dots represent individual replicates. FDR - Benjamini-Hochberg corrected significance of enrichment across three experiments (see Methods).

To test whether CTAP-formation occurs in natural contexts, we re-analyzed public deep proteome profiles of healthy human tissues (**Extended Data Fig. 6a, b**). Terminal amides were preferentially found on neo-C-termini, indicating that CTAPs can form upon protein fragmentation (**Extended Data Fig. 6c**). Up to 427 such neo-C-termini affecting up to 165 different proteins were detected per tissue sample (**Extended Data Fig. 6d, e**). Since such fragments are expected to be rare under physiological conditions, potentially due to active clearance by FBXO31, we screened for additional endogenous CTAPs in FBXO31 knockout cells. We stably expressed catalytically inactive affinity-tagged FBXO31 (HA-FBXO31(ΔF-box)) in FBXO31 knockout HEK293T cells and performed co-IP followed by MS. Under basal conditions, FBXO31 co-purified with select proteins, including all core members of the tRNA ligase complex. Notably, the tRNA ligase complex is subject to oxidative hydroxyl radical damage under regular culture conditions^30^ (**Fig. 4b, Supplementary Table 3**).

Following acute oxidative stress (20 min, 200 µM H_2_O_2_), we detected several additional putative FBXO31 clients, such as the aminoacyl tRNA synthetase AARS1 and the glutamine synthetase GLUL (**Fig. 4c, Supplementary Table 4**). Mis-trafficked secreted proteins were not found to co-IP with FBXO31 in any condition. Individual co-IP Western blotting confirmed that FBXO31(ΔF-box) binds certain clients only after H_2_O_2_ treatment (**Fig. 4d**). This binding is furthermore dependent upon threonine 343 in the FBXO31 peptide binding pocket, which we previously found to be critical for amidated peptide recognition. FBXO31 therefore preferentially recognizes multiple endogenous clients under oxidative stress. Notably, the bound client AARS1 exhibited a substantially reduced molecular weight relative to the full-length protein, implying an internal cleavage event.

We tested whether the identified FBXO31 clients undergo primary amide-forming fragmentation in cells. We performed IP-MS on affinity-tagged AARS1 in FBXO31 knockout cells following oxidative challenge (1h, 200 µM H_2_O_2_). A search for *de novo* C-termini of AARS1 identified two internal cleavage sites with a mass shift of −0.98 Da at their C-termini, consistent with amide modification (**Fig. 4e**). Cleavage at these sites would reduce the molecular weight of AARS1 by −30 kDa, corresponding to the fragment of endogenous AARS1 we observed bound by FBXO31 (**Fig. 4d**). Amidated versions of both AARS1 C-terminal peptides induced potent reporter degradation in cells (**Fig. 4f**). This effect was only observed in presence of a C-terminal amide and was entirely rescued by FBXO31 knockout.

Based on these results, we conclude that FBXO31 acts as a protein quality control factor that recognizes and removes endogenous CTAPs formed under oxidative stress, particularly ones arising from oxidative backbone cleavage. This is consistent with the ability of FBXO31 to broadly recognize termini that could theoretically occur anywhere within a protein. It also potentially accounts for FBXO31’s slight preference for hydrophobic amino acids that tend to be found in protein cores instead of at native C-termini. Since random damage across the proteome would evade detection by shotgun MS, the reported CTAPs may represent hotspots of more wide-spread damage by hydroxyl radicals.

### A dominant cerebral palsy-associated mutation in FBXO31 alters substrate selection

Two previous studies reported a dominant *de novo* D334N mutation in FBXO31 among patients with diplegic spastic cerebral palsy^31, 32^. This mutation eliminates the negative charge in the substrate binding pocket of FBXO31 and was speculated to act by increased degradation of Cyclin D1. We asked whether the D334N mutation alters FBXO31 substrate recognition and CTAP clearance.

*In vitro*, neither wildtype nor D334N mutant FBXO31 showed any affinity for the proposed C-terminal degron of Cyclin D1 (**Extended Data Fig. 7a**). However, the D334N mutation did abolish binding to C-terminal amide peptides (**Extended Data Fig. 7b, c**). This held true in a pooled peptide interaction screen covering >1200 peptides, where the D334N mutation displayed globally reduced CTAP binding (**Extended Data Fig. 7d, Supplementary Table 5**). Likewise, FBXO31(D334N, ΔF-box) expressed in FBXO31 knockout cells failed to immuno-precipitate both a model CTAP (mCherry-CONH_2_) and unmodified Cyclin D1 (**Extended Data Fig. 7e**). However, simple loss of function does not account for the dominant inheritance of the D334N mutation. We therefore hypothesized that this mutation might also lead to recognition of neosubstrates.

To identify cellular targets of FBXO31(D334N), we performed co-IP MS of FBXO31(ΔF-box) using both the wildtype and D334N mutant cDNAs. Mutant FBXO31(D334N, ΔF-box) formed detectable interactions with 220 proteins, 195 of which were not detected for the wildtype (**Fig. 5a and Supplementary Table 6**). We tested whether these putative neo-substrates are also removed from cells in response to acute FBXO31(D334N) expression using a ligand-inducible shield-degron system (DD)^33^. Tandem mass tag (TMT) expression proteomics identified a marked reduction in the abundance of many of the bound candidates within 12 h of stabilizing DD-FBXO31(D334N), but their abundance was unaffected when expressing wildtype FBXO31 (**Fig. 5b, Extended Data Fig. 7f, Supplementary Table 7**). Among these newly bound and degraded substrates were core-essential proteins including the SUGT1 central regulator of mitotic spindle assembly^34, 35^.

**Fig 5.**
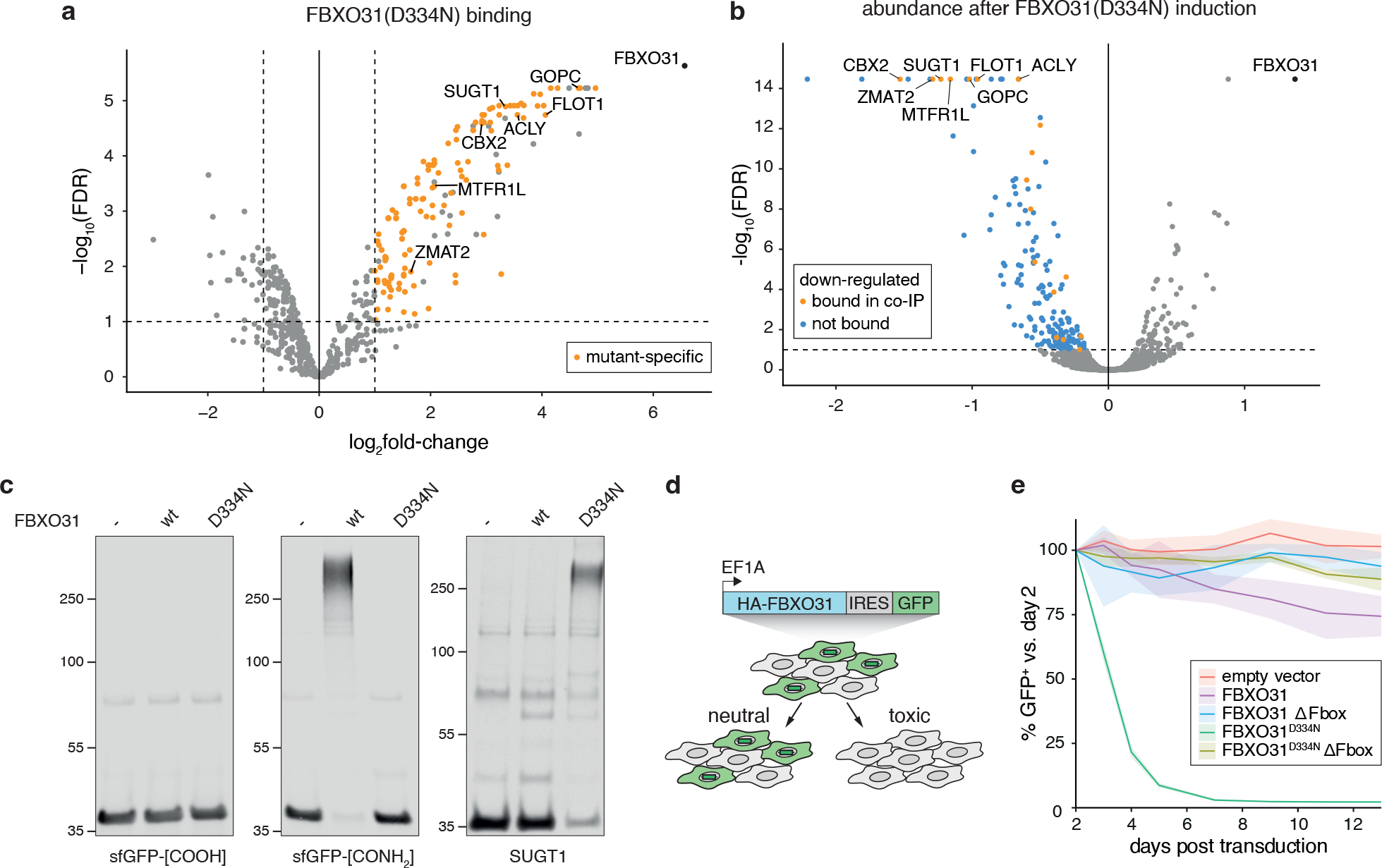
The Cerebral-palsy mutation D334N shifts substrate selectivity of FBXO31. (**a**) Identification of FBXO31(D334N) neosubstrates by IP-MS of HA-tagged FBXO31(ΔF-box, D334N) from FBXO31 knockout HEK293T cells. Proteins that do not co-IP with wt FBXO31 are highlighted in orange. (**b**) Identification of proteome changes following acute induction of FBXO31(D334N) expression. FBXO31 knockout cells were stably transduced with ligand-inducible DD-FBXO31(D334N). Total protein abundance was measured by TMT-MS after 12 h of protein induction (2 µM shield-1) and compared to shield-treated parental cells. Down-regulated proteins (FDR < 0.1, log2FC < 0, three independent replicates) are highlighted. (**c**) *In vitro* ubiquitylation of an unmodified or CTAP model substrate (sfGFP-[X]) or SUGT1 by wt or mutant FBXO31. (**d**) Schematic of a competitive proliferation assay in FBXO31 knockout HEK293 cells. The fraction of cells stably expressing GFP-linked cDNAs is followed over time by flow cytometry. Lines indicate mean of three experiments. Ribbons indicate the standard deviation. (**e**) Results of the proliferation assay shown in (d) for indicated cDNAs. FDR - Benjamini-Hochberg corrected significance of enrichment across three experiments (see Methods).

*In vitro* reconstitution of SCF/FBXO31 ubiquitylation confirmed that the D334N mutation abolished its CTAP-directed activity (**Fig. 5c**). Conversely, FBXO31(D334N) robustly ubiquitylated the neosubstrate SUGT1. Degradation of SUGT1 and the other candidate FBXO31(D334N) neosubstrates would be expected to impair cell survival or proliferation. We therefore performed a competitive growth assay to quantify the impact of FBXO31(D334N) expression on cell fitness (**Fig. 5d**). While wildtype FBXO31 cDNA expression was well tolerated in FBXO31 knockout HEK293 cells, the D334N mutant was rapidly depleted from co-culture (**Fig. 5e**). Deletion of the F-box motif required for SCF complex assembly fully abolished this effect, demonstrating that SCF/FBXO31(D334N) exerts a toxic ubiquitin ligase activity. Based on these results we conclude that D334 is required for CTAP recognition, and the cerebral-palsy associated mutation is dominant because it redirects ubiquitin ligase activity away from C-terminal amide substrates and towards essential cellular proteins.

## Discussion

To maintain protein homeostasis, cellular machinery must survey thousands of proteins for their chemical integrity. We propose that this can be achieved by modified amino acid degrons (MAADs) that mark proteins for removal by reader proteins and downstream effectors. We identified a C-terminal amide as a *bona fide* MAAD found on oxidatively damaged proteins, marking them for degradation by SCF/FBXO31. A role of SCF/FBXO31 in surveilling oxidative protein damage would parallel the surveillance of oxidative genome damage by enzymes such as OGG1 and MUTYH^36^.Notably, FBXO31 is highly expressed in metabolically active cells, including skeletal muscles and the brain^37^. By removing irreversibly damaged protein fragments from these tissues, FBXO31 could prevent accumulation of protein decay products.

It remains possible that CTAPs are also formed by means other than oxidative damage, for example, by a so-far unknown PAM-like lyase in the cytoplasm. We found no evidence for secretory proteins in our immunoprecipitation or proteomic datasets, but proteins that are amidated by PAM in the secretory pathway might also become FBXO31 clients if internalized or mis-localized to the cytoplasm under certain conditions.

FBXO31’s recognition of a C-terminal amide could reconcile conflicting reports on its role in Cyclin D1 degradation. An earlier study of stress-induced CyclinD1 downregulation suggested a direct role of FBXO31 in its destabilization^38^, partly supported by reports of a direct yet low affinity interaction^28^. Subsequent studies in other cell types and states could not confirm a universal role of FBXO31 in CyclinD1 regulation and instead implicated the ligase AMBRA^39, 40^. We also found no evidence for unmodified Cyclin D1 degradation by SCF/FBXO31. However, C-terminal amidation could render Cyclin D1 an FBXO31 client in other cell types or following stress, and we found that an amidated Cyclin D1 peptide could direct recognition by FBXO31.

The activity of sequence-based degrons can be modulated by PTMs that facilitate or block binding of ubiquitin ligases to defined amino acid motifs. Examples include some of the first described degrons, such as the cell cycle-regulated phospho-degron of Sic1^41^, or the hydroxyproline-degron recognized by VHL^42^. These are strictly dependent on a defined sequence context and typically have regulatory functions. In contrast, the C-terminal amide MAAD acts in a largely sequence-independent manner as an apparent quality control factor. While a great deal of effort has been put into finding sequence-based targets for the ∼600 ubiquitin ligases in the human genome, it may be that a portion of these orphan ligases recognize MAADs. The human proteome is extensively modified, with at least 50% of peptides in deep MS scans carrying one of more than 300 catalogued modifications^9, 43^. While we focused on the effect of candidate MAADs on protein turnover, our semi-synthetic screening approach can be extended to chart the impact of chemical modifications on all steps in a protein’s life cycle.

MAADs have so far been described in very few cases, but with remarkable potential for therapeutic use. It has recently been appreciated that the CRBN ligase, often harnessed for targeted protein degradation, recognizes cyclized C-terminal imides^6^. Given that MAAD readers must recognize a well-defined chemical ligand while surveilling a broad range of putative clients, it is possible that MAADs are entry points for developing degradation-inducing compounds. Indeed, we found that addition of a C-terminal amide was sufficient to induce the degradation of all host proteins we tested. Harnessing native chemical recognition instead of repurposing sequence-based recognition could provide a promising alternative path to future targeted protein degradation drugs.

## Methods

### Peptide synthesis

Detailed synthetic methods and characterization of all peptides, pooled peptide libraries, custom peptide building blocks, and protein-peptide conjugates generated for this study are provided in Supplementary File 1. Peptide sequences are provided in (**Extended Data Table 1**).

### General method for the preparation of peptides for sortase reaction

Amino acids were loaded on chlorotrityl resin to access C-terminal carboxylic acids or Rink amide to access C-terminal amides (see Supplementary File 1). Automated peptide elongation was carried out on a Multisyntech Syro I parallel synthesizer according to the general peptide methods (Supplementary File 1). The peptide was cleaved from the resin using TFA/DODT/H_2_O (95:2.5:2.5, v/v) for 1 h. The resin was removed by filtration and the filtrate concentrated under reduced pressure. The solution was triturated with Et_2_O and centrifuged to obtain crude peptide. The crude peptide was dissolved in H_2_O/CH_3_N (1:1, v/v) + 0.1% (v/v) TFA and purified using preparative HPLC. The peptide series pep_AARS1, pep_lib_VIN and pep_lib_YNR were obtained from Craftide Co. Ltd (Nagoya).

### Sortase-mediate protein conjugation

Peptides were dissolved in sortase reaction buffer (50 mM Tris, 150 mM NaCl, pH 7.4, at 4 °C). pH was adjusted to 7-8. Peptide (final concentration 1 mM) was mixed with reporter protein (sfGFP, mCherry, BFP) (final concentration 75 µM). SortA 7M was added (final concentration 2 µM) and incubated at 4 °C for 18 h. Unreacted reporter protein and cleaved sortag were removed by Ni-NTA purification. The flow-through was collected and immediately buffer exchanged to ion exchange buffer (25 mM Tris, pH 8.5) using a desalting column (Cytiva). The sample was further purified by anion exchange using a MonoQ column (Cytiva) with a gradient of 0-25% high salt buffer (25 mM Tris, 1 M NaCl, pH 8.5) in 25CV. Fractions containing the product were pooled, buffer-exchanged to sortase reaction buffer supplemented with 0.5 mM TCEP and concentrated.

### General cell culture methods

HEK293 and HEK293T cells were cultured in DMEM containing Glutamax (Thermo Fisher Scientific) and K562 cells were cultured in RPMI 1640 containing Glutamax (Thermo Fisher Scientific) under standard conditions. Growth media were further supplemented with 10% (v/v) Fetal Bovine Serum (Thermo Fisher Scientific) and 100 U/ml Penicillin-Streptomycin (Thermo Fisher Scientific). Stably transduced cells were selected using 2 µg/ml Puromycin (Thermo Fisher Scientific), 500 µg/ml Geneticin (Thermo Fisher Scientific) or 10 µg/ml Blasticidin (Thermo Fisher Scientific) starting two days after transduction.

Induction of doxycycline-dependent vectors was performed by addition of 500 ng/ml doxycycline (Merck) every 48 hours. Cells were treated with epoxomicin (Merck), folimycin A (concanamycin A, Merck), TAK243 (MedChem Express) or MLN4924 (MedChem Express) as indicated in figure legends. H_2_O_2_ was freshly diluted in serum-free DMEM growth medium to 200 µM immediately before use and was kept protected from direct light. Cell lines were regularly tested for mycoplasma contamination using the MycoAlert Mycoplasma Detection Kit (Lonza).

### Virus production

Lentiviral vectors were packaged in HEK293T cells using standard methods. In brief, cells were incubated with plasmid DNA (transfer plasmid, pCMV-dR8.2 dvpr and pCMV-VSV-G at a weight-ratio of 4:2:1) and PEI (Polyethyleneimine, Mw ∼25’000 Da) at a ratio of 1:3 (w/w) of total plasmid DNA to PEI. Viral supernatant was harvested at 48-72 hours post nucleofection by ultrafiltration and supplemented with 4 µg/ml polybrene. Gene knockdown was performed in a sub-clonal K562 cell line stably expressing dCas9-KRAB-BFP (Addgene plasmid #102244), generously provided by Eric J. Aird (Corn Lab, ETH Zürich). For CRISPRi, sgRNAs targeting near transcription start sites were chosen based on previously optimized design rules (21) and cloned into vector pCRISPRia. Puromycin was used to select for cells stably expressing sgRNAs. All custom vectors generated for this study will be made available through Addgene upon publication of this manuscript. A complete list of plasmids used in this study is provided in (**Extended Data Table 2**).

### Flow cytometry assays and cell sorting

Analytical flow cytometry was performed on an Attune NxT Flow Cytometer (Thermo Fisher Scientific). Single-cell isolation and sorting for CRISPR-screen analysis was performed on a Sony SH-800 cell sorter.

For measuring stability of recombinant or semi-synthetic proteins, 1.5-2.0 x 10^5^ cells were electroporated with 200 pmol of protein in 20 µl cuvettes using a 4D nucleofector device (Lonza) according to the manufacturer’s recommendations. Following protein delivery, cells were left to recover at 37°C for 30-45 minutes and washed with PBS before measuring initial mean fluorescence intensity values (t_0_) by flow cytometry. Subsequent measurements were performed at indicated time points. Baseline-fluorescence values of mock-electroporated cells receiving no protein were measured in parallel and subtracted for each time point. Background-corrected fluorescence values were normalized to t_0_.

Cell-surface expression of CD55 was assessed by immunostaining and flow cytometry. In brief, cells were pelleted and resuspended in primary antibody diluted 1:200 in PBS containing 5 % FBS. (APC-anti-human-CD55[JS11], Biolegend). After 20 minutes of incubation on ice, cells were washed three times with PBS followed by data acquisition. For competitive growth assays, cells were transduced with lentiviral vectors delivering FBXO31-IRES-GFP cDNA variants or an empty vector control (IRES-GFP) at a multiplicity of infection (MOI) of about 0.3. The fraction of transduced cells was measured every two days by flow cytometry and normalized to initial transduction levels (day 2) as a measure of relative fitness.

### Generation of clonal cell lines

For genome-wide CRISPR screening, K562 cells were made competent for inducible gene knockout (iCas9) by transduction with vectors SRPB (pHR-SFFV-rtTA3-PGK-Bsr) and 3GCasT (pHR-TRE3G-hSpCas9-NLS-FLAG-2A-Thy1.1). Cas9-P2A-Thy1.1 expression was induced with doxycycline for 2 days and cells staining positive for Thy1.1 were isolated by single-cell sorting. Clonal lines were screened for cells that show no evidence of CD55 knockout following viral delivery of sgCD55 (**Extended Data Table 3**) by antibody staining and flow cytometry, as well as efficient knockout following addition of doxycycline for 9 days.

FBXO31 knockout cells were generated by electroporation of cells with Cas9/sgRNA ribonucleoprotein particles (RNP) as described previously (41) using sgRNAs listed in (**Extended Data Table 3**) based on previously optimized design rules (17, 42). In brief, *in vitro* transcription templates were generated by PCR using Q5 polymerase (New England Biolabs) and primers listed in (**Extended Data Table 3**) and used for *in vitro* transcription by T7 RNA polymerase (New England Biolabs). Resulting RNA was purified using a spin-column kit (RNeasy mini kit, QIAGEN) and 120 pmol of sgRNA were complexed with 100 pmol of recombinant SpCas9 protein at RT for 20 minutes. Cas9 protein was provided by GEML (FGCZ of Uni Zürich and ETH). Assembled sgRNA/Cas9 complexes were delivered to cells using a 4D nucleofector kit (Lonza) according to the manufacturer’s instructions. Clonal cell lines were isolated by single-cell sorting and characterized by genomic DNA extraction (Lucigen QuickExtract), PCR amplification of edited loci using Q5 polymerase (NEW ENGLAND BIOLABS) with genotyping and NGS primers (**Extended Data Table 3**). Pooled next generation sequencing of edited loci was performed by GEML (FGCZ of Uni Zürich and ETH) on a MiSeq sequencer (Illumina) with 150 bp paired-end reads. Deep sequencing reads were analyzed using CRISPResso2 (43) to identify compound heterozygous knockout clones.

### Western blotting & antibodies used in this study

Unless for co-IP studies, immunoblotting was performed on whole cell lysates in radioimmunoprecipitation buffer (RIPA, 50 mM Tris-HCl, 150 mM NaCl, 0.25% deoxycholic acid, 1% NP-40, 1 mM EDTA, pH 7.4) supplemented with protease inhibitors (Halt protease inhibitor cocktail, Thermo Fisher Scientific). Proteins were analyzed by polyacrylamide gel electrophoresis (PAGE) and wet transfer onto nitrocellulose membranes (0.2 µm pore size) using standard methods. Membranes were blocked with TBS-T (150 mM NaCl, 20 mM Tris, 0.1% (w/v) Tween-20, pH 7.4) containing 5% skimmed milk powder and incubated with primary antibodies diluted in TBST containing 5% bovine serum albumin (BSA) and 0.05 % (w/v) sodium azide. Primary western blotting antibodies used in this study are FBXO31 (Abcam, ab86137), GFP (Abcam, ab6556), mCherry (Abcam, ab183628), SKP1 (Cell Signaling Technology, 2156), CUL1 (Invitrogen, 71-8700), AARS1 (Fortis/Bethyl Life Science, A303-473A), GLUL (Fortis/Bethyl Life Science, A305-323A), Cyclin D1 (Abcam, ab134175), HA (Cell Signaling Technology, 3724), and SUGT1 (Bethyl, A302-944A). Detection of primary antibodies was performed using far-red fluorescently labeled secondary antibodies (LI-COR Biosciences, cat. nr. 926-32213 and 926-68072) on an Odyssey CLx scanner (LI-COR Biosciences).

### CRISPR screening and analysis

iCas9 cells were transduced with the pooled lentiviral sgRNA library TKOv3 (17) at a multiplicity of infection (MOI) of about 0.3 as measured by serial dilution, puromycin selection and viability assay (CellTiter-Glo 2.0, Promega). Two pools of 1.2 · 10^8^ cells each were transduced, yielding a ca 500-fold library coverage, which was maintained throughout all cell culture steps. Cas9-expression was induced by addition of doxycycline for 5 days prior to delivery of reporter proteins. To isolate CTAP-clearance deficient cells ≥ 4 large-scale nucleofections were performed by combining 5 · 10^7^ cells, 2000 p mol sfGFP-pep1-CONH_2_ (sfGFP-GGGKDLEGKGGSAGSGSAGGSKYPYDVPDYAKS-[CONH_2_]) and 2000 p mol of a degron-tagged control protein (mTagBFP2-RxxG) in a 100 µl nucleofection reaction (4D nucleofector kit SE plus supplement SF1, Lonza) 14 hours following nucleofection, cells were transferred on ice and sorted into a CTAP clearance-deficient population (sfGFP^+^ mTagBFP2^-^) and a control population (sfGFP^-^ mTagBFP^-^). Genomic DNA was extracted from snap-frozen sorted cells using the Gentra Pure kit (QIAGEN). sgRNA cassettes were isolated by two rounds of PCR using a previously published strategy (17) with NEBNext Ultra II Q5 Master Mix (New England Biolabs) and primer pairs listed in **Extended Data Table 2**.

Protospacers were quantified by deep sequencing using 21 initial dark cycles on a NextSeq 2000 device (Illumina) by the Genome Engineering and Measurement Lab of the Functional Genomics Center Zürich (GEML, FGCZ). sgRNA counts were retrieved using mageck count (MAGeCK v0.5.9.3) with default parameters (44). Raw sgRNA counts are provided as a supplementary table (**Supplementary Table 8**). Enrichment of sgRNAs targeting the same gene in CTAP clearance-deficient cells versus the control population was estimated with mageck test using a paired design for screening duplicates with option --remove-zero both and otherwise default parameters.

### Recombinant protein production

Codon-optimized cDNAs of SKP1(A2P, Δ38-43, Δ71-82) and H_6_-Smt3-StrepII-FBXO31.1(Δ1-65) were generated by gene synthesis (Integrated DNA Technologies) based on previously optimized expression constructs (26, 45) and cloned into the bacterial expression vector pET28b (**Extended Data Table 2**). Expression was performed at 18°C overnight in presence of 0.5 mM IPTG in E. coli strain BL21(DE3) in Terrific Broth (Faust AG) in presence of 50 µg/ml kanamycin. Following bacterial cell lysis by sonication in buffer NBA (200mM NaCl, 50 mM HEPES, 25 mM imidazole, 1 mM TCEP, pH 8.0), recombinant protein was enriched on a HisTrap FF column (Cytiva) by elution with 250 mM imidazole in the same buffer. The H_6_-Smt3 tag was removed by digestion with purified SUMO protease Ulp1 (46) at 4°C overnight. Binary SKP1/FBXO31 complexes were further purified by anion exchange chromatography (HisTrap Q HP, Cytiva) in 50 mM HEPES (pH7.5) and 5 mM DTT on a linear gradient of 100-1000 mM NaCl followed by gel filtration (HiPrep Sephacryl S-100 HR, Cytiva) in 200mM NaCl, 50 mM HEPES and 1mM TCEP. Sortase (SortA 7M) was prepared by recombinant expression *E. coli* using a previously reported method^44^.

### Pooled library IP and TIMS-TOF analysis

For each IP, 1 pmol of peptide library was incubated with an equimolar amount of recombinant Strep II-tagged FBXO31/SKP1 complex in 1 ml of binding buffer (200mM NaCl, 25mM HEPES pH 7.5, 1 mM TCEP) with pre-equilibrated magnetic Strep-Tactin XT beads (240 µl MagStrep type 3 XT bead slurry, IBA life sciences). Following 1 h of binding at room temperature on a rotator, beads were washed twice with binding buffer, twice with wash buffer (150 mM NaCl, 25 mM HEPES, 1 mM TCEP) and eluted in 150 mM Nacl, 50 mM HEPES, 100 mM biotin (pH 7.7). Eluted peptides and an equal amount of input library (1 pmol) were diluted 1:1 with triethylammonium bicarbonate (50 mM, pH 8.0). Peptides were isobarically-labeled with TMT 2-plex (Thermo Fisher Scientific) labels according to the manufacturer’s instructions. Input library was modified with TMT-126 and eluted peptides with TMT-127. Following labeling, peptides were combined 1:1. Combined peptides were desalted using 100 µL C18 ZipTips and dried (Savant SpeedVac). The dried peptides were resuspended in 5 vol% acetonitrile, 0.1 vol% formic acid.

Tandem MS experiments were performed using ESI-TIMS-QTOF-MS system (TimsTOF Pro, Bruker Daltonics, Germany) with collision-induced dissociation (CID) and N_2_ as the collision gas. Peptides were pressure loaded onto a reversed phase 25 cm × 75 µm i.d. C18 1.6 μm column (Ionoptics Ltd., Australia) with a reversed phase 5 mm × 0.3 mm i.d. C18 5 μm column (Thermo Scientific, Lithuania) as guard column at 40 °C. The mobile phase consisted of (A) water with 0.1% formic acid and (B) acetonitrile with 0.1% formic acid. The gradient started at 2% of B and was linearly increased to 35% in 120 min at flowrate of 300 µl min^−1^. A second gradient profile, started at 35% of B and was linearly increased to 95% in 2 min at flowrate of 300 µl min^−1^. Followed by isocratic conditions of 95% B at flowrate of 300 µl min^−1^ for 8 min. Total run time, including the conditioning of the column to the initial conditions, was 163 min. Further data processing was performed with Data Analysis 5.3 software (Bruker Daltonics, Germany) using a processing script to generate export files and reports.

Mascot Server 2.8.1. (Matrix Science) was used to match spectra against a custom reference of common contaminants concatenated with all potential peptides generated by pooled SPPS (10 ppm peptide mass tolerance, 0.05 Da fragment mass tolerance and for peptide-amide libraries −0.98 Da at peptide termini as variable modification). For each peptide, reporter ion intensities were summed across multiple detections and scaled to the sum of reporter ion intensities for all input library peptides yielding relative reporter ion intensities (RRI) for cross-comparisons between runs. For each experiment, two parallel IPs were analyzed. For wt FBXO31 co-IP, two MS-measurements were run for each IP for improved library coverage.

### *In vitro* ubiquitylation assay

2 µM of recombinant FBXO31/SKP1 complexes were combined with 0.5 µM nM of substrate, 50 µM recombinant ubiquitin, 50 nM UBE1, 500 nM UBE2R1, 500 nM UBE2D3, and 500 nM CUL1∼NEDD8/RBX1 complex in 50 mM Tris/HCl (pH 7.5), 100 mM NaCl, 10 mM MgCl_2_, 10 mM ATP and 0.5 mM TCEP. Reactions were incubated at 30°C for 1h unless indicated otherwise and analyzed by immunoblotting. All protein components except for FBXO31/SKP1 and model substrates were sourced commercially (R&D Systems).

### HA-FBXO31 co-IP

Co-IP of semi-synthetic model substrates (mCherry-GGGRRLEGKEEDEKGSRASDDFRDLR-[COOH/CONH_2_]) was performed in FBXO31 knockout HEK293 cells stably transduced with pLenti-EF1A-HA-FBXO31-PGK-Neo or indicated mutant derivatives (**Extended Data Table 2**). Cells were nucleofected 2 hours prior to harvest and treated with 2 µM MLN4924 (MedChemExpress) and 500 nM epoxomicin (Merck) to block protein degradation and stabilize CRL complexes. 2 µM MLN4924 and 500 nM epoxomicin were added at all subsequent steps until cell lysis.

Co-IP of endogenous FBXO31 clients was performed in FBXO31 knockout HEK293T cells stably transduced with pLenti-EF1A-HA-FBXO31(ΔF-box)-IRES-GFP or mutant derivatives of it (**Extended Data Table 2**). Cells were cultured in serum-free DMEM media supplemented with 500 nM epoxomicin with or without addition of H_2_O_2_ at a final concentration of 200 µM for 1 h prior to cell harvest.

For both, endogenous and synthetic client IP, cells were harvested by centrifugation and washed in ice cold PBS followed by lysis in SCF-IP buffer (150mM NaCl, 50 mM Tris-HCl, 1mM EDTA, 0.1% Igepal CA-630, 5% (v/v) glycerol, pH 7.5) supplemented with Halt protease inhibitor cocktail (Thermo Fisher Scientific) by rotation for 30 min at 4°C. Debris was removed by centrifugation (4°C, 30 min, 20’000 g) and total protein concentrations measured using the Bradford protein assay (Thermo Fisher Scientific). Typically, 140 µg of protein were incubated with 12 µl of Pierce anti-HA magnetic bead resin (Thermo Fisher Scientific) on a rotator wheel at 4°C overnight. Beads were washed three times for 5 minutes in SCF-IP buffer and bound proteins were eluted twice using 0.1 M Glycine (pH 2), followed by addition of 0.25 volumes neutralization buffer (1.5 M NaCl, 0.5 M Tris-HCl, pH 8.0).

### co-IP MS of HA-tagged FBXO31

IP-MS of HA-tagged FBXO31 was performed with FBXO31 knockout HEK293T cells stably expressing HA-FBXO31(ΔF-box) from vector pLenti-EF1A-HA-FBXO31(ΔF-box)-IRES-GFP or an indicated mutant cDNA. For each condition, three replicate samples were processed from separately passaged cultures. As negative control, we used identical FBXO31 knockout HEK293T cells not expressing bait cDNAs. For IP of oxidatively stressed cells, cultures were incubated in serum-free DMEM media supplemented with 500 nM epoxomicin with or without addition of H_2_O_2_ at a final concentration of 200 µM for 1 h prior to cell harvest.

Cells were trypsinized and washed twice with ice cold PBS containing 500 nM epoxomicin and resuspended in lysis buffer (300 mM NaCl, 50 mM Tris-HCl, 0.5% (v/v) Igepal CA-630) freshly supplemented with Halt Protease Inhibitor Cocktail (Thermo Fisher Scientific). Cell lysates were obtained by rotation for 30 minutes at 4°C and were cleared by centrifugation (4 °C, 30 min, 20’000 g). For IP, 500 µg protein were incubated with 80 µl of pre-equilibrated bead slurry (Pierce anti-HA magnetic beads, Thermo Fisher Scientific) in a total volume of 250 µl of lysis buffer on a rotator wheel at 4°C for 4 h. Magnetic beads were washed four times with lysis buffer and twice with PBS followed by elution with 80 µl 0.1 M Glycine (pH 2.0) at RT with gentle shaking for 10 min. Supernatant was neutralized with 0.25 volumes of 1.5 M NaCl, 0.5 M TrisHCl pH 7.5.

Eluates were processed by the proteomics group of FGCZ (Uni Zürich/ETH), by in-solution tryptic digest and LC-MS/MS on an Orbitrap Exploris 480 Mass Spectrometer (Thermo Fisher Scientific) in data-dependent acquisition mode. Peptide search was performed using MSFRagger 3.4 (47) matching spectra against the UniProt human proteome (UP000005640) and common contaminants and filtering peptides for FDR ≤ 0.01. Enrichment of interactors in HA co-IP experiments was calculated using limma 3.54, by fitting a linear model with emperical bayes moderation to spectral counts of control-IP and HA-IP samples with Benjamini-Hochberg correction for multiple hypothesis testing.

### IP-MS of FLAG-tagged AARS1

IP-MS of 3xFLAG-tagged AARS1 was performed on FBXO31 knockout HEK293T cells transiently transfected with pCDNA4TO-CMV-3xFLAG-Halo-AARS1 using 30 µg of polyethyleneimine (M_W_∼25’000) and 10 µg of plasmid DNA. Two days after transfection, cells were challenged with 200 µM H_2_O_2_ in serum-free DMEM. Five minutes following H_2_O_2_ treatment cells were additionally supplemented with 500 nM epoxomicin. Cells were washed three times in PBS supplemented with 500 nM epoxomicin and snap-frozen. Cell pellets were lysed by resuspension in RIPA buffer (50 mM Tris-HCl, 150 mM NaCl, 0.25% deoxycholic acid, 1% NP-40, 1 mM EDTA, pH 7.4) supplemented with protease inhibitors (Halt protease inhibitor cocktail, Thermo Fisher Scientific). 250 µg of protein were incubated with 25 µl of pre-equilibrated anti-FLAG resin (M2 anti-FLAG beads, Merck) on a rotator at 4°C overnight. Beads were washed once with RIPA buffer, four times with high-salt RIPA buffer (50 mM Tris-HCl, 500 mM NaCl, 0.25% deoxycholic acid, 1% NP-40, 1 mM EDTA, pH 7.4) and twice with TBS (150 mM NaCl, 50 mM Tris pH 7.5) followed by elution in 100 µl TBS + 150 ng/µL competitor peptide (Pierce 3x DYKDDDDK Peptide, Thermo Fisher Scientific) at 37 °C for 30 min.

Eluates were processed by the proteomics group of FGCZ (Uni Zürich/ETH), by in-solution digest with Glu-C protease and LC-MS/MS data dependent acquisition on an Orbitrap Exploris 480 Mass Spectrometer (Thermo Fisher Scientific). Peptide search was performed using MSFRagger 3.4 (47) matching against the UniProt human proteome (UP000005640) and common contaminants with precursor and fragment mass tolerances of 20 ppm, up to 1 missed cleavage, semi-specific cleavage and addition of a C-terminal peptide amide (−0.98 Da) as variable modification. For identification of amide-bearing de novo C-termini, internal AARS1-matching peptides were filtered for modified C-termini not matching a Glu-C cleavage site (D or E).

### Expression proteomics by tandem mass tag (TMT) MS

For inducible expression of DD-3xFLAG-FBXO31, FBXO31 knockout HEK293 cells were transduced with pLenti-EF1A-DD-3xFLAG-FBXO31-PGK-Neo or its D334N mutated variant (**Extended Data Table 2**). Cells were treated with 2 µM Shield-1 (MedChem Express) for 12 h prior to harvest. All experiments were performed with three replicate samples treated and harvested on separate days. Cell pellets were processed for whole proteome quantification by FGCZ (Uni Zürich/ETH). In brief, cell pellets were lysed in 4 % Sodium dodecyl sulfate (SDS) in 100 mM Tris/HCl pH 8.2 by boiling and mechanical homogenization. Per sample, 50 µg of total protein were reduced (2 mM TCEP) and alkylated (15 mM 2-Chlroacetamide) and further processed using a fully automated SP3 purification, digest and clean-up workflow. Samples were isobarically labeled with TMT10plex (Thermo Fisher Scientific), pre-fractionated by RP-HPLC on an XBridge Peptide BEH C18 column (Waters) and pooled into eight fractions. LC-MS/MS acquisition was performed on an Orbitrap Exploris 480 mass spectrometer (Thermo Fisher Scientific) in DDA mode. Raw MS data was processed using ProteomeDiscoerer 2.4 (Thermo Fisher Scientific). Spectra were searched against the UniProt human reference proteome (downloaded on 29.4.2022) and common contaminants with Sequest HT with a peptide-level FDR cutoff of 0.01. Detection of differentially abundant proteins was performed on reporter ion intensities for D334N or wt FBXO31-expressing cells compared to Shield-1-treated control cells lacking FBXO31 cDNA.

### Re-analysis of public proteomes for CTAP detection

Label-free MS-based proteomics of tryptic digests of healthy human tissues from single donors were downloaded from PRIDE archive PXD010154, samples 1277 (brain), 1499 (adipose tissue), 1306 (thyroid), and 1296 (heart)^45^. Raw spectra were transformed into mzML files using Thermo Raw File Parser^46^. Spectra were searched against the human proteome (Uniprot reference UP000005640) using the MSFragger 3.7 search engine^47^ through the Philosopher software suite^48^ using a precursor mass tolerance of 20 ppm and a fragment mass tolerance of 0.05 Da. Peptide identification was based solely on b- and y-series ions allowing for up to 1 missed cleavage and up to 1 non-enzymatic peptide terminus with variable peptide C-terminal amidation (−0.984016 Da), M oxidation (+15.99490 Da), protein C-terminal acetylation (+ 42.01060 Da) and fixed C carbamidomethylation (+57.021464 Da). Filtering for high-confidence peptide spectrum matches (PSMs), peptides and proteins was performed using default parameters (FDR ≤ 0.01). Fully enzymatic peptides were defined as having termini that match trypsin cleavage sites or annotated N and C-termini, accounting for clipping of N-terminal M. Semi-enzymatic peptides were defined as having one cleavage that does not match an annotated protein terminus or trypsin cleavage site.

### Visualization of protein structures and alignments

All protein structures were downloaded from the RCSB Protein Data Bank (https://www.rcsb.org/) under accession numbers noted in respective figure legends. Structures and electrostatic potential maps were rendered using ChimeraX-1.5 (48). The composite structure of SCF/FBXO31 was generated from substructures 5VZU (FBXO31) and 6TTU (NEDD8-CUL1-RBX1-SKP1). Structures were aligned by their shared SKP1 substructure in using MatchMaker with default settings. Alignment and conservation analysis of select FBXO31 orthologs was performed using JalView (49).

### FP assays and analysis

Increasing concentrations of FBXO31-SKP1 complex (0–5 µM) were incubated with fluorescein-labelled peptides (2-10 nM) in 25 mM HEPES, 150 mM NaCl, pH 7.5 and incubated at room temperature for 30 min. Fluorescence polarization was measured on a Victor Nivo Plate Reader (Perkin Elmer). Following subtraction of baseline signal, polarization levels were normalized to the highest signal of each pair of amidated and non-amidated peptides for wt FBXO31. Binding curves were fitted by least squares regression and half-maximal binding concentrations were extracted using Prism 9 (Graphpad) assuming one-site binding, maximal binding at 100% of the measured signal and no contribution from unspecific interactions. Dissociation constants for the highest affinity probes reported in this study are close to the probe concentrations used for FP assays and are therefore only reported as approximate values.

## Acknowledgements

We thank Eric J. Aird, Marija Banović, John Fielden, M. Erman Karasu and Jin Rui Liang and all other members of the Corn and Bode labs for critical discussions. Tobias Kockmann, Laura Kunz, Chia-wei Lin, Sibylle Pfammatter and Paolo Nanni of the Functional Genomics Center Zürich (FGCZ, ETH/University of Zürich) supported this work through proteomics sample processing, acquisition and data analysis. Louis Bertschi, Michael Meier, and Bertran Rubi of the Molecular and Biomolecular Analysis Service (MoBiAS) in the Department of Chemistry and Applied Bioscience (ETHZ) provided HRMS and assisted with multiplex peptide binding screens. Susanne Kreutzer and Zacharias Kontarakis of the Genome Engineering and Measurement Lab (GEML, ETH/University of Zürich) provided recombinant SpCas9 and a viral sgRNA library and performed deep sequencing. We thank Jason Greenwald (ETH Zürich, D-CHAB) for helpful discussions and support regarding protein expression.

This work was supported by an ETH Zurich Postdoctoral Fellowship program and an EU Horizon 2020, Marie Skłodowska-Curie grant 898175 (MM), by the Scholarship Fund of the Swiss Chemical Industry (RH), and by the NOMIS Foundation and the Lotte and Adolf Hotz-Sprenger Stiftung (JEC).

## Author contributions

Conceptualization: MM, JF, RH, JWB, JEC; Writing – original draft: MM, JF, JWB, JEC; Writing – original, review and editing: MM, JF, RH, JWB, JEC; Investigation: MM, JF, RH, MC, NDS

## Materials & Correspondence

Deep sequencing read counts are provided as supplementary table. Complete lists of proteins identified by IP-MS and TMT-MS are provided as supplementary tables and corresponding raw data will be made available upon publication through the Proteomics Identification Database (PRIDE, www.ebi.ac.uk/pride/). Plasmids generated for this study will be made available through Addgene (www.addgene.org/). Correspondence regarding any materials used in this study should be addressed to JWB and JEC.

---

**Extended Data Fig. 1.**
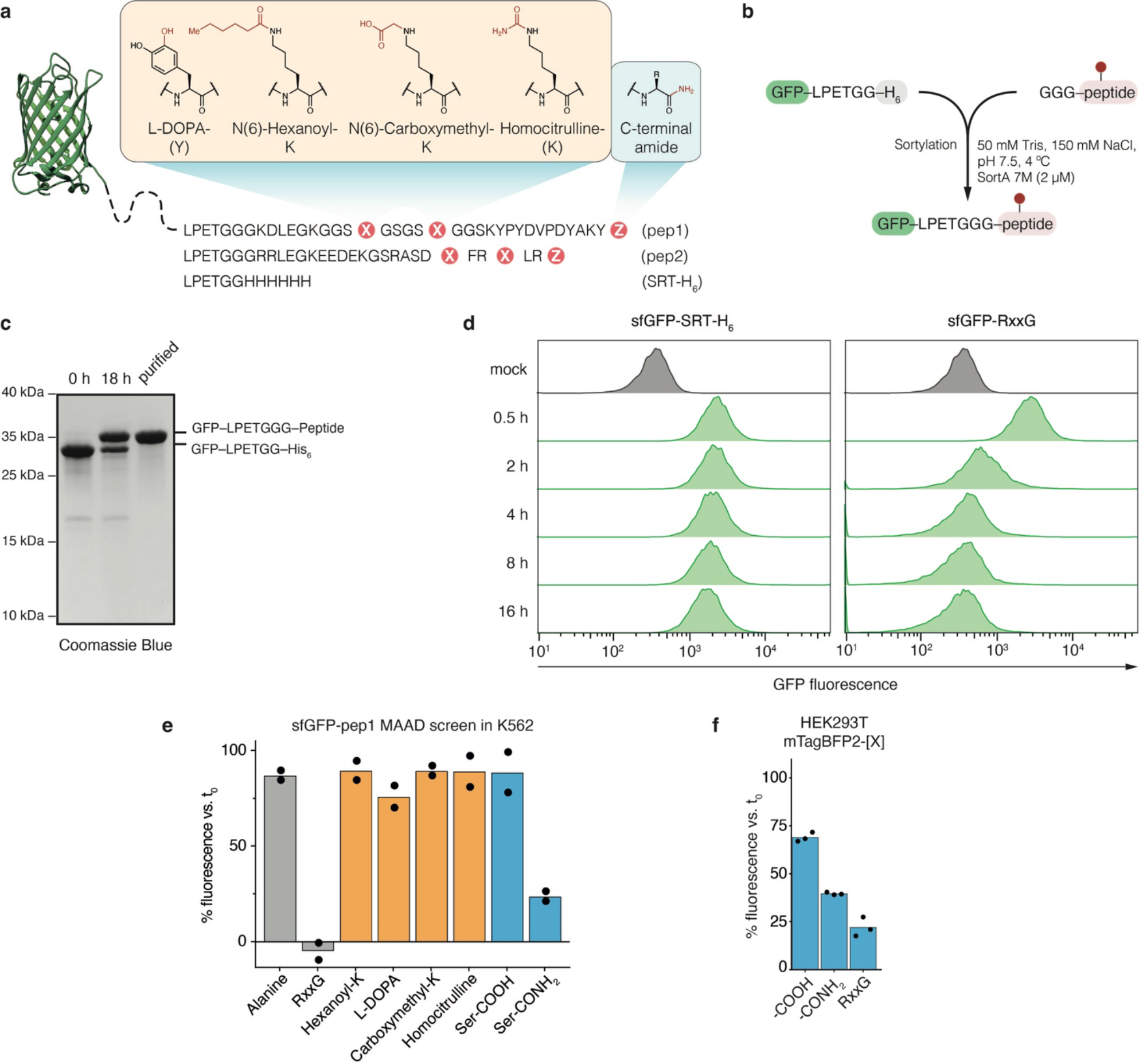
A screen for modified amino acid degrons (MAAD). (**a**) Schematic showing sfGFP bearing various PTMs tested for MAAD activity. Modified sequences are shown and modified positions are highlighted. The modification on each amino acid is show in red. Corresponding unmodified amino acids are indicated in brackets. (**b**) Schematic showing modification of sfGFP with a C-terminal sortylation tag. “SortA 7M” conjugates sfGFP and peptide containing an N-terminal GGG-motif. (**c**) Exemplary SDS-PAGE analysis of sortase reaction showing sfGFP, crude reaction mixture and purified sfGFP-conjugate. Conjugate was purified by Ni-NTA to remove unreacted sfGFP, cleaved sortag and SortA 7M, followed by ion exchange chromatography. (**d**) Exemplary time-course experiment measuring sfGFP turnover in K562 cells. Cells received sfGFP carrying a C-terminal sortase tag (sfGFP-SGGLPETGGHHHHHHV) or its conjugated form carrying an RxxG degron motif (sfGFP-SGGLPETGGGRRLEGKEEDEKGSRASDRFRGLR). (**e**) Results of a screen for MAAD activity. Pep1 MAAD reporters shown in (a) were used in an in-cell reporter assay as in (d) in K562 cells. (**f**) Validation of C-terminal amidation as a MAAD as in (e) in HEK293T cells using mTagBFP conjugated to amidated (-CONH2) or unmodified (-COOH) pep2 shown in (a) or the positive control degron (RxxG) as in (d). Bars represent mean of independent replicate experiments. Black dots show individual measurements.

**Extended Data Fig. 2.**
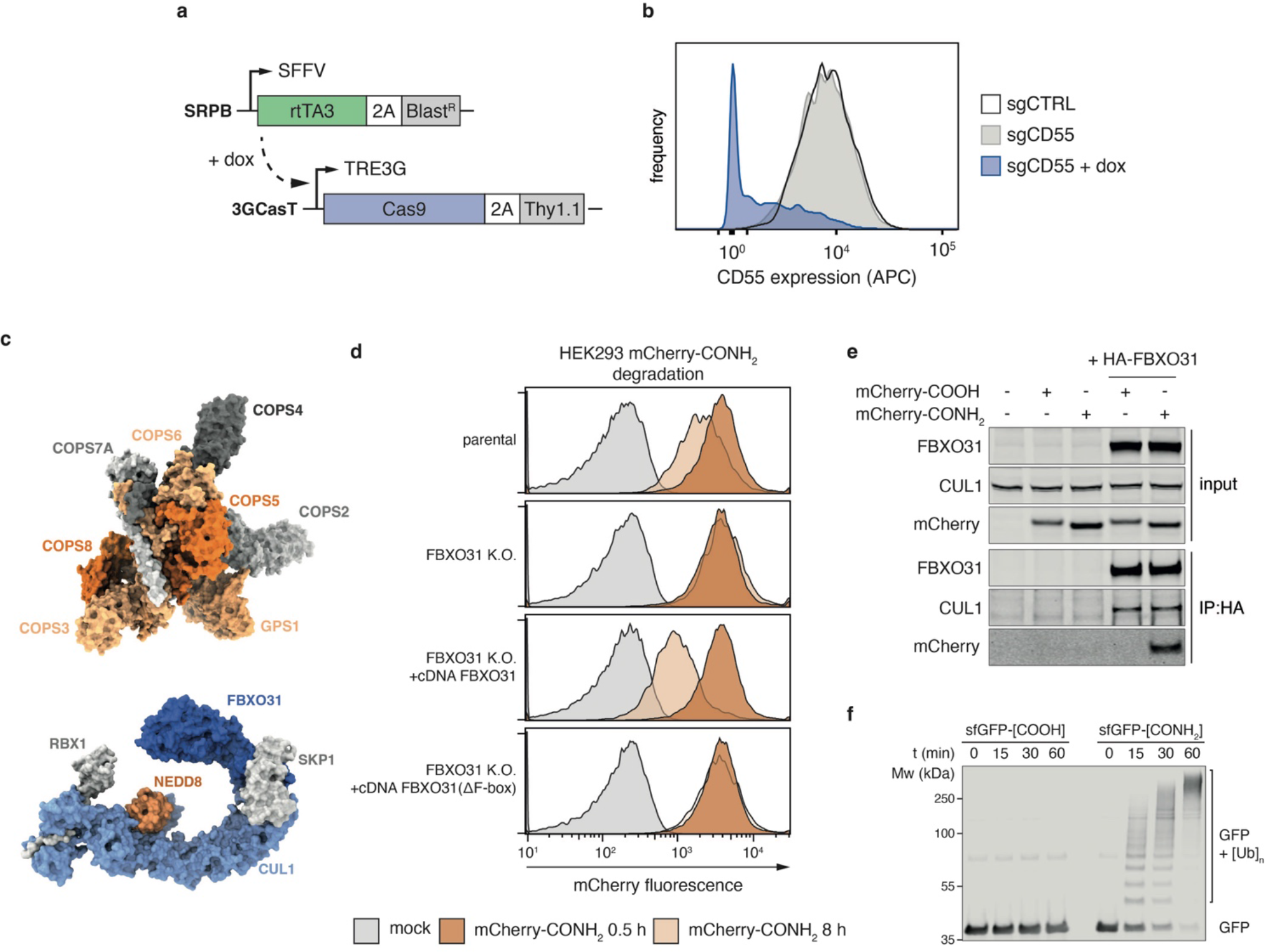
Inducible CRISPR screen identifies SCF/FBXO31 as a CTAP clearance factor. (**a**) Schematic of the vectors used to establish a doxycyline (dox)-inducible Cas9 expression system (iCas9) for CRISPR screening. (**b**) Validation of inducible gene disruption in iCas9 cells using an sgRNA targeting cell surface marker CD55. Cells stably expressing sgCD55 were left untreated or on dox (500 ng/ml) for 9 days. Histogram shows CD55 surface expression measured by flow cytometry. (**c**) Rendering of the COP9 signalosome complex from PDB:4D10 (top) and SCF/FBXO31 based on previously published structures (bottom, see Methods). Screening hits identified for CTAP clearance and NEDD8 are highlighted in blue or orange. (**d**) CTAP degradation assay in FBXO31 knockout cells with or without stable transduction of indicated cDNA rescue constructs, as well as non-edited HEK293 cells (parental). (**e**) Co-IP of HA-FBXO31 from FBXO31 knockout HEK293 cells electroporated with the model substrate mCherry-pep2 (mCherry-GGGRRLEGKEEDEKGSRASDDFRDLR) with indicated C-termini. Cells were co-treated with the neddylation inhibitor MLN4924 (2 µM) and 500 nM epoxomicin and harvested after 2 h to sttabilize CRL complexes and to prevent protein degradation. HA-FBXO31 was stably expressed as cDNA. (**f**) *In vitro* ubiquitylation time course of sfGFP-pep1 (sfGFP-GGGKDLEGKGGSAGSGSAGGSKYPYDVPDYAKS) with indicated C-termini incubated with SCF/FBXO31, E1 and E2 enzymes.

**Extended Data Fig. 3.**
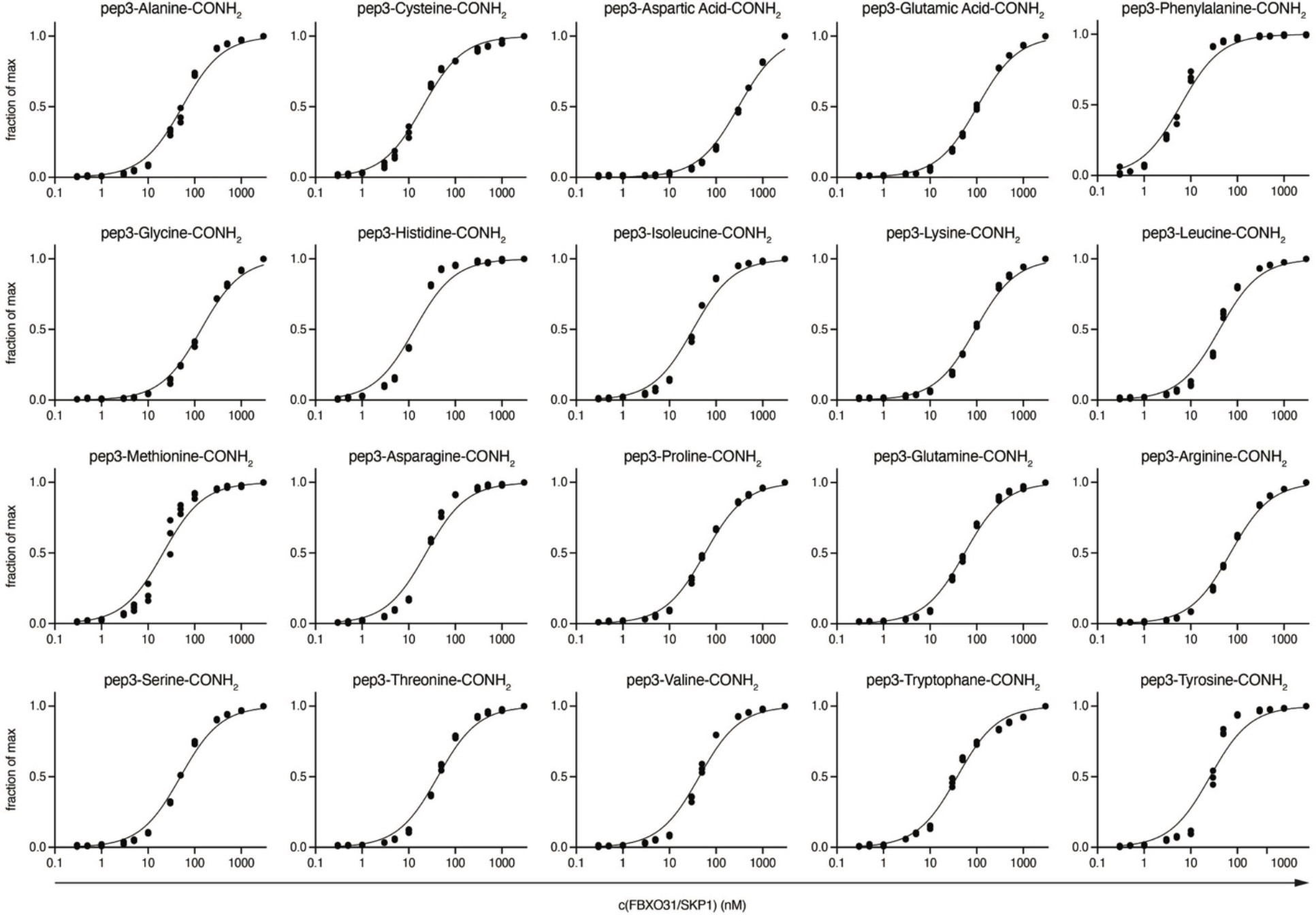
CTAP binding across terminal amino acid contexts by FBXO31. Individual binding curves of pep3 (KKYRYDVPDYSA[X]-CONH2) with each of the 20 canonical proteinogenic amino acids in the C-terminal positions. Corresponding dissociation constants are summarized in Fig. 3b.

**Extended Data Fig. 4:**
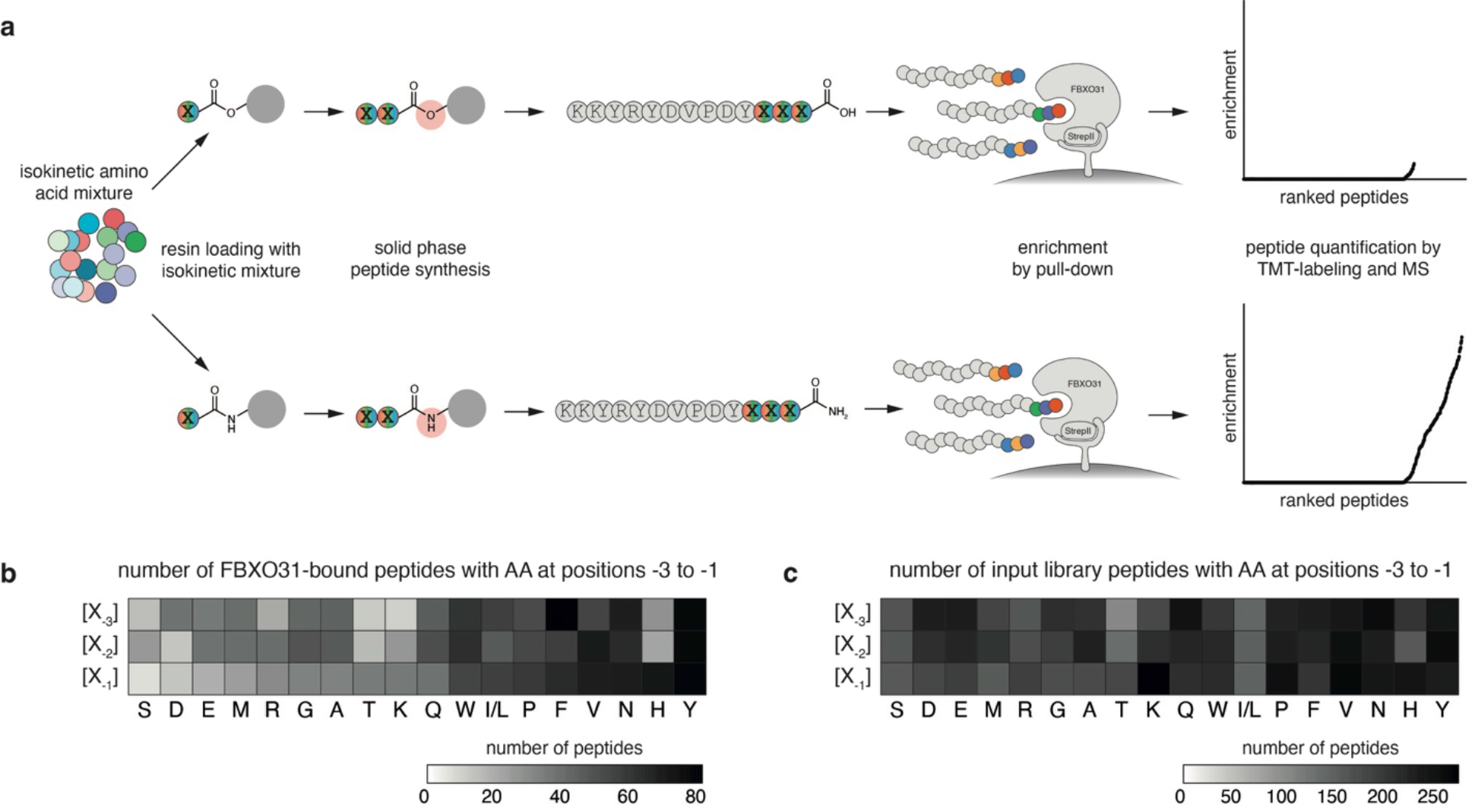
A pooled *in vitro* interaction screen for CTAP binding preferences of FBXO31. (**a**) Schematic showing analysis of peptide library with variable C-terminal amino acids by FBXO31 pull-down and subsequent MS/MS analysis. Peptide libraries are prepared using isokinetic amino acid mixtures. (**b**) Heatmaps showing absolute amino acid frequencies in the three terminal positions of 841 unique C-terminally amidated peptides co-precipitated with FBXO31. (**c**) Heatmap as in (b) for all 3817 unique peptides identified in the input library.

**Extended Data Fig. 5:**
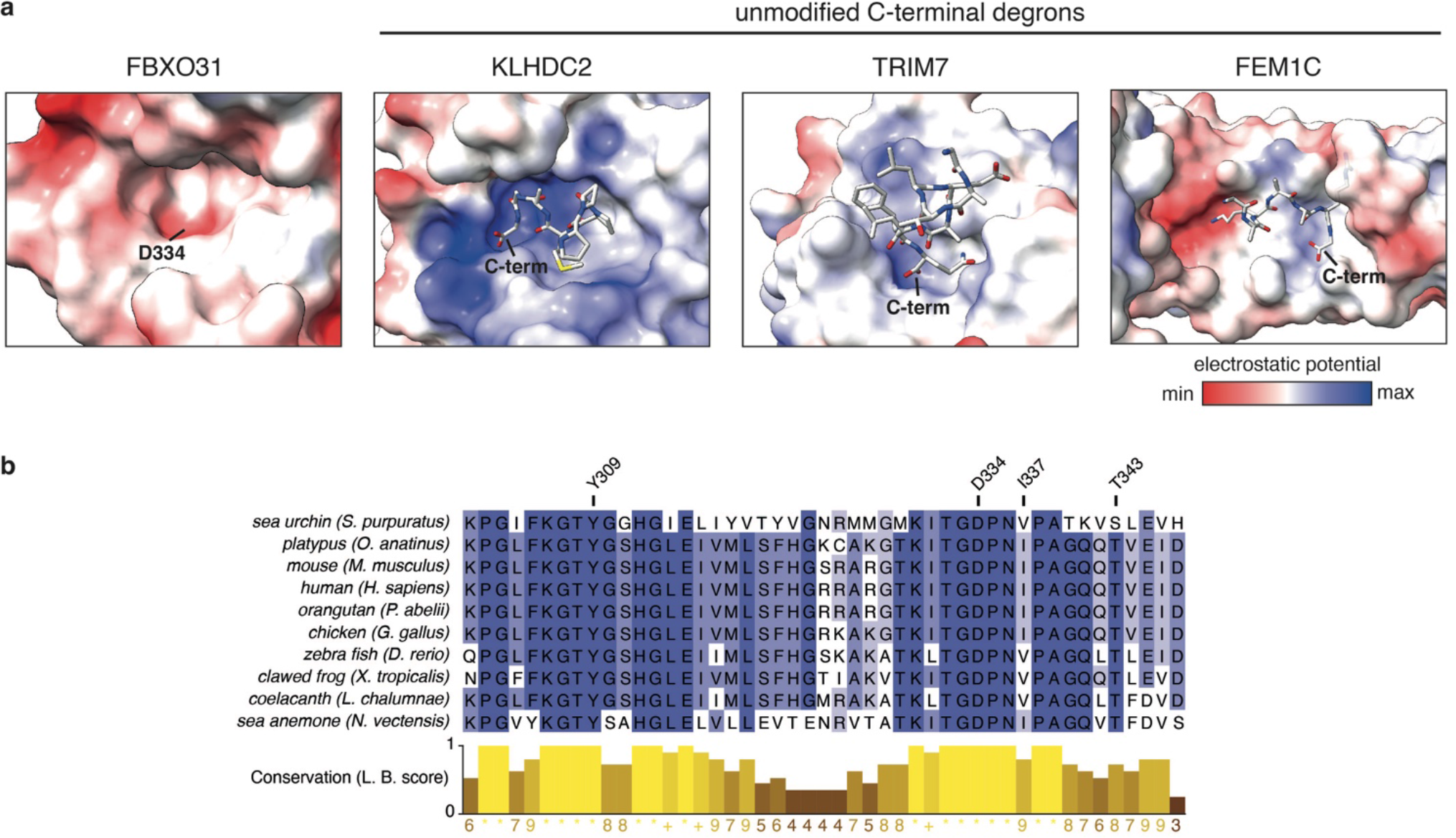
A conserved binding pocket enables amide-recognition by FBXO31. (**a**) Comparison of the negatively charged binding substrate pocket of FBXO31 with readers of unmodified C-terminal degrons. Electrostatic potential maps were rendered based on published structures with PDB accessions 5VZT (FBXO31), 6DO3 (KLHDC2), 7Y3A (TRIM7) and 6LEY (FEM1C). (**b**) Protein sequence alignment of the FBXO31 substrate binding pocket from orthologs of indicated species. Conservation is measured with the Linvingston-Barton conservation score.

**Extended Data Fig. 6:**
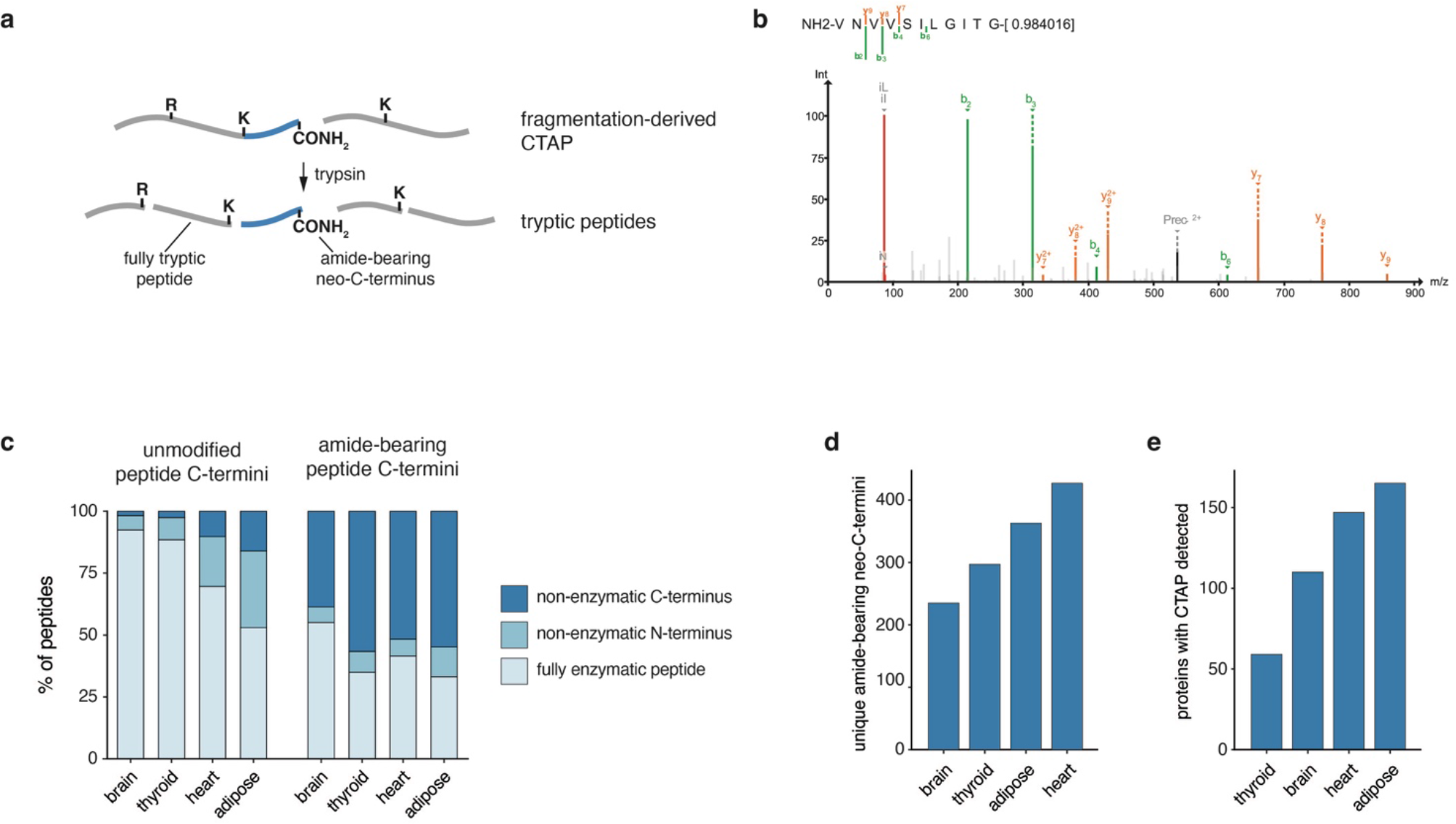
Evidence of CTAP formation in human tissue proteomes. (**a**) Schematic showing a CTAP stemming from alpha-amidating protein fragmentation and resulting peptides following tryptic digest for mass spectrometry. (**b**) Sample spectrum of a peptide indicative of an amide-bearing neo-C-terminus identified in a public proteome dataset from healthy human brain tissue (PRIDE Project PXD010154). (**c**) Re-analysis of public proteome data from indicated tissues (see Methods). Peptides were separated into such with unmodified C-termini or a putative C-terminal amide (−0.984016 Da). Bar graphs show the percentage of peptides that are fully enzymatic or have one non-enzymatic N-or C-terminus. (**d**) Number of unique amide-bearing neo-C-termini in indicated tissues as analyzed in (c). (**e**) Number of proteins showing evidence of CTAP-formation in (d).

**Extended Data Fig. 7:**
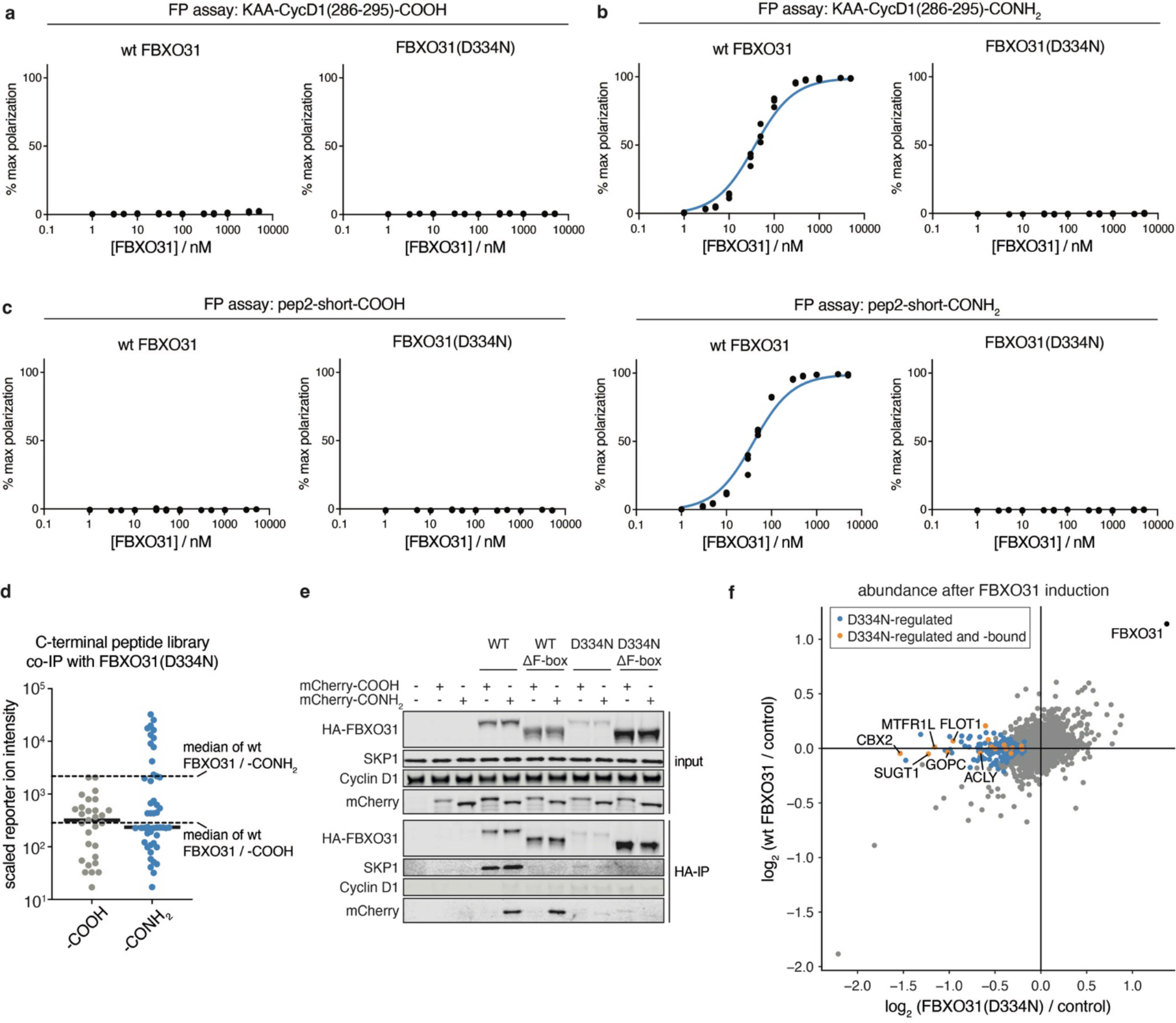
The D334N mutation disrupts CTAP recognition by FBXO31. (**a**) FP assay of for a peptide derived from the C-terminus of Cyclin D1 ([fluorescein]-KAATPTDVRDVDI-COOH) with wildtype or D334N mutant FBXO31/SKP1. (**b**) FP assay as in (a) with the same peptide bareing a C-terminal amide. (**c**) FP assays as in (a) and (b) for a second peptide pair (pep2-short, KEEDEKGSRASDDFRDLR). (**d**) Results of a pooled peptide interaction screen as in Extended Data Fig. 4. Points represent individual peptides co-purifying with FBXO31(D334N). Ordinate values indicate relative enrichment measured by TMT reporter ion intensitiy scaled to the input library. Dotted lines indicate median enrichments for unmodified (-COOH) and amidated (-CONH2) libraries with wt FBXO31. (**e**) co-IP of indicated HA-tagged FBXO31 cDNAs in FBXO31 knockout HEK293 cells electroporated with a model substrate (mCherry-GGGRRLEGKEEDEKGSRASDDFRDLR). Cells were co-treated with 2 μM MLN4924 and 500 nM epoxomicin and harvested after 2h. (**f**) Comparison of proteome-wide responses to FBXO31 induction for wildtype or D334N mutant DD-3xFLAG-FBXO31 induced by 12 h of treatment with shield-1 measured by TMT-MS. Proteins are highlighted in blue if they are down-regulated only by mutant FBXO31 expression (FDR < 0.1) and in orange if they are also detected by FBXO31(D334N, ΔF-box) coIP-MS (FDR < 0.1).

