## Supplementary figures, tables and methods for "C-terminal amides mark proteins for degradation via SCF/FBXO31"

### Supplementary File 1

### Supplementary Figures

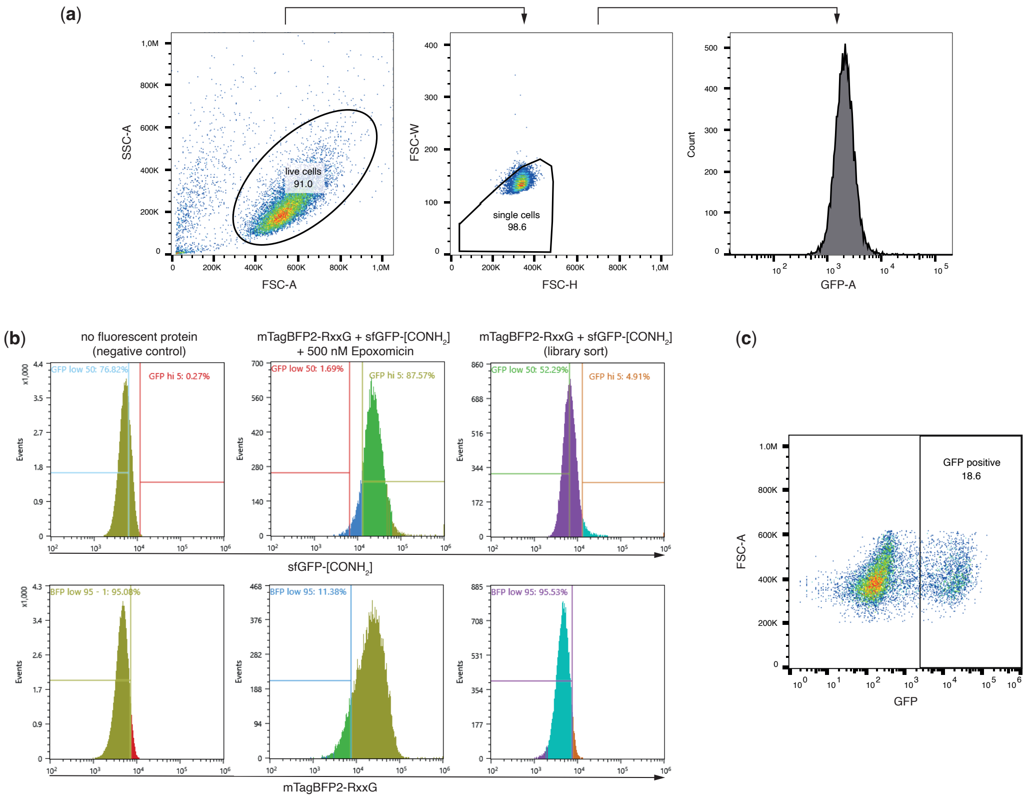

**Supplementary Fig. 1: Gating strategies for flow cytometry experiments.** (**a**) Representative flow cytometry plots showing the gating strategy for measuring reporter protein levels in K562 cells. Total events were gated to include live cells and exclude duplets before extracting median GFP fluorescence values. (**b**) Gating strategy used for isolating CTAP-clearance deficient cells (BFP low & GFP high) and a CTAP-proficient background population following live-cell gating and doublet exclusion as in (a). Set-up control samples received no fluorescent reporter proteins (negative control) or BPF-RxxG and GFP-CONH$_2$ followed by proteasome inhibition (500 nM epoxomicin) to assess fluorescence levels of cells deficient for reporter degradation. Gates were manually adjusted during the sort to maintain initial gating precentages. (**c**) Gating strategy used for quantifying cDNA overexpressing cells (GFP+) in competitive growth assays following live-cell gating and doublet exclusion as in (a).

### Extended Data Tables

**Extended Data Table 1 - Peptide sequences used in this study**

| name | sequence | use |
|---|---|---|
| pep1 [homocitrulline (HCT)]–D–COOH | GGGKDLEGKGGS[HCT]GSGS[HCT]GGSKYPYDVPDYAKD–[COOH] | conjugation |
| pep1 [L-DOPA (LDO)]–D–COOH | GGGKDLEGKGGS[LDO]GSGS[LDO]GGSKYPYDVPDYAKD–[COOH] | conjugation |
| pep1 [carboxymethyllysine (CML)]–D–COOH | GGGKDLEGKGGS[CML]GSGS[CML]GGSKYPYDVPDYAKD–[COOH] | conjugation |
| pep1 [A]–S–COOH | GGGKDLEGKGGSAGSGSAGGSKYPYDVPDYAKS-[COOH] | conjugation |
| pep1 [A]–S–CONH2 | GGGKDLEGKGGSAGSGSAGGSKYPYDVPDYAKS-[CONH2] | conjugation |
| pep1 [A]–D–COOH | GGGKDLEGKGGSAGSGSAGGSKYPYDVPDYAKD–[COOH] | conjugation |
| pep1 [hexanoyllysine (KHL)]–D–COOH | GGGKDLEGKGGS[KHL]GSGS[KHL]GGSKYPYDVPDYAKD–[COOH] | conjugation |
| pep2 [DxxD]–R–COOH | GGGRRLEGKEEDEKGSRASDDFRDLR–[COOH] | conjugation |
| pep2 [DxxD]–R–CONH2 | GGGRRLEGKEEDEKGSRASDDFRDLR–[CONH2] | conjugation |
| pep2 [RxxG]–R–COOH | GGGRRLEGKEEDEKGSRASDRFRGLR–[COOH] | conjugation |
| pep2 [RxxG]–R–CONH2 | GGGRRLEGKEEDEKGSRASDRFRGLR–[CONH2] | conjugation |
| pep_lib_VIN-COOH | GGGKYRYDVPDYVIN-[COOH] | conjugation |
| pep_lib_VIN-CONH2 | GGGKYRYDVPDYVIN-[CONH2] | conjugation |
| pep_lib_YNR-COOH | GGGKYRYDVPDYYNR-[COOH] | conjugation |
| pep_lib_YNR-CONH2 | GGGKYRYDVPDYYNR-[CONH2] | conjugation |
| pep_AARS1(664-684)-COOH | KAVYTQDCPLAAAKAIQGLRA-[COOH] | conjugation |
| pep_AARS1(664-684)-CONH2 | KAVYTQDCPLAAAKAIQGLRA-[CONH2] | conjugation |
| pep_AARS1(664-683)-COOH | KAVYTQDCPLAAAKAIQGLR-[COOH] | conjugation |
| pep_AARS1(664-683)-CONH2 | KAVYTQDCPLAAAKAIQGLR-[CONH2] | conjugation |
| fluorescein-pep2_short–R–COOH | [Fluorescein]–KEEDEKGSRASDDFRDLR–[COOH] | FP assays |
| fluorescein-pep2_short–R–CONH2 | [Fluorescein]–KEEDEKGSRASDDFRDLR–[CONH2] | FP assays |
| fluorescein-pep2_short–R–CH2OH | [Fluorescein]-KEEDEKGSRASDDFRDLR–[CH2OH] | FP assays |
| fluorescein-pep2_short–R–COOMe | [Fluorescein]- KEEDEKGSRASDDFRDLR–[COOMe] | FP assays |
| fluorescein-KAA-CycD1(286-295)–COOH | [Fluorescein]– KAATPTDVRDVDI–[COOH] | FP assays |
| fluorescein-KAA-CycD1(286-295)–CONH2 | [Fluorescein]– KAATPTDVRDVDI–[CONH2] | FP assays |
| fluorescein-pep3–**X**–CONH2 | [Fluorescein]-KKYRYDVPDYSAX–[CONH2] | FP assays |
| fluorescein-pep3–N–COOH | [Fluorescein]-KKYRYDVPDYSAN–[COOH] | FP assays |
| fluorescein-pep3–Q–COOH | [Fluorescein]-KKYRYDVPDYSAQ–[COOH] | FP assays |
| pep_lib_COOH | GGGKYRYDVPDYXXX-[COOH] | pooled library |
| pep_lib_CONH2 | GGGKYRYDVPDYXXX-[CONH2] | pooled library |

**Extended Data Table 2 - Plasmids used in this study**

| short name | description | reference | Addgene ID | type |
|---|---|---|---|---|
| pCMV-VSV-G | helper plasmids for lentiviral packaging | Stewart et al. 2003 | 8454 | transient transfection |
| pCMV-dR8.2 dvpr | helper plasmids for lentiviral packaging | Stewart et al. 2003 | 8455 | transient transfection |
| pCDNA4TO-CMV-3xFLAG-Halo-AARS1 | expression of FLAG-AARS1 for IP | this study | tba | transient transfection |
| pHR-SFFV-rtTA3-PGK-Bsr ("SRPB") | overexpression of Tet-transactivator (rtTA3) | this study | tba | lentiviral cDNA vector |
| pHR-TRE3G-hSpCas9-NLS-FLAG-2A-Thy1.1 ("3GCasT") | tet-responsive expression of Cas9 | this study | tba | lentiviral cDNA vector |
| pLentiX-EF1A-HA-FBXO31-PGK-Neo | FBXO31 cDNA expression | this study | tba | lentiviral cDNA vector |
| pLentiX-EF1A-HA-FBXO31(ΔF-box)-PGK-Neo | FBXO31 cDNA expression | this study | tba | lentiviral cDNA vector |
| pLentiX-EF1A-HA-FBXO31(Y309A)-PGK-Neo | FBXO31 cDNA expression | this study | tba | lentiviral cDNA vector |
| pLentiX-EF1A-HA-FBXO31(I337D)-PGK-Neo | FBXO31 cDNA expression | this study | tba | lentiviral cDNA vector |
| pLentiX-EF1A-HA-FBXO31(T343V)-PGK-Neo | FBXO31 cDNA expression | this study | tba | lentiviral cDNA vector |
| pLentiX-EF1A-HA-FBXO31(D334N)-PGK-Neo | FBXO31 cDNA expression | this study | tba | lentiviral cDNA vector |
| pLentiX-EF1A-HA-FBXO31(ΔF-box, D334N)-PGK-Neo | FBXO31 cDNA expression | this study | tba | lentiviral cDNA vector |
| pLentiX-EF1A-IRES-GFP | empty vector control | this study | tba | lentiviral cDNA vector |
| pLentiX-EF1A-HA-FBXO31-IRES-GFP | FBXO31 cDNA expression | this study | tba | lentiviral cDNA vector |
| pLentiX-EF1A-HA-FBXO31(ΔF-box)-IRES-GFP | FBXO31 cDNA expression | this study | tba | lentiviral cDNA vector |
| pLentiX-EF1A-HA-FBXO31(D334N)-IRES-GFP | FBXO31 cDNA expression | this study | tba | lentiviral cDNA vector |
| pLentiX-EF1A-HA-FBXO31(ΔF-box, D334N)-IRES-GFP | FBXO31 cDNA expression | this study | tba | lentiviral cDNA vector |
| pLentiX-EF1A-HA-FBXO31(ΔF-box, K330A)-IRES-GFP | FBXO31 cDNA expression | this study | tba | lentiviral cDNA vector |
| pLentiX-EF1A-HA-FBXO31(ΔF-box, T343V)-IRES-GFP | FBXO31 cDNA expression | this study | tba | lentiviral cDNA vector |
| pLentiX-EF1A-DD-3xFLAG-FBXO31-IRES-GFP | FBXO31 cDNA expression | this study | tba | lentiviral cDNA vector |
| pLentiX-EF1A-DD-3xFLAG-FBXO31(D334N)-IRES-GFP | FBXO31 cDNA expression | this study | tba | lentiviral cDNA vector |
| TKOv3 | genome-wide human sgRNA library | Hart et al. 2017 | 125517 | lentiviral sgRNA vector |
| pCRISPRia-v2 | expression of individual sgRNAs for CRISPRi | Horlbeck et al. 2016 | 84832 | lentiviral sgRNA vector |
| pLenti-U6-sgCD55-EFS-eBFP2-P2A-Puro | expression of individual sgRNA for CD55 knockout | this study | tba | lentiviral sgRNA vector |
| pET28-sfGFP-SRT-His6 | recombinant production of sortase-tagged sfGFP | this study | tba | bacterial protein production |
| pET28-mCherry-SRT-His6 | recombinant production of sortase-tagged mCherry | this study | tba | bacterial protein production |
| pET28-mTagBFP2-SRT-His6 | recombinant production of sortase-tagged mTagBFP2 | this study | tba | bacterial protein production |
| pET28-SKP1(A2P, Δ38-43, Δ71-82)-FBXO31(Δ1-65) | recombinant expression of core FBXO31/SKP1 complex | this study | tba | bacterial protein production |
| pET28-SKP1(A2P, Δ38-43, Δ71-82)-FBXO31(Δ1-65, D334N) | recombinant expression of core FBXO31/SKP1 complex | this study | tba | bacterial protein production |
| pFGET19_Ulp1 | production of UlpI for Smt3-tag cleavage | Guerrero et al. 2015 | 64697 | bacterial protein production |

**Extended Data Table 3 - Oligonucleotide sequences used in this study**

| sequence name | sequence (5`-3`) | notes | type |
|---|---|---|---|
| sgFBXO31.KO3 | ATCAGGTGGATCCTGAACAG | protospacer of sgRNA targeting the CDS of FBXO31, delivered as RNP | protospacer |
| sgFBXO31.VC1 | GCCGTACATGCACTCCACTG | protospacer of sgRNA targeting the CDS of FBXO31, delivered as RNP | protospacer |
| sgCD55 #1 | CTGGGCATTAGGTACATCTG | protospacer of sgRNA targeting the CDS or CD55, expressed from pLenti-U6-sgCD55-EFS-eBFP2-P2A-Puro | protospacer |
| sgNT | GCTGAAGCACTGCACGCCAT | protospacer of non-targeting control sgRNA, expressed from pCRISPRia-v2 | protospacer |
| sgFBXO31 #1 | GTCTCCACCAGCAGCTCGGG | protospacer of sgRNA for CRISPRi targeting FBXO31, expressed from pCRISPRia-v2 | protospacer |
| sgFBXO31 #2 | GCGGTGTGTGCTCGCCTTTG | protospacer of sgRNA for CRISPRi targeting FBXO31, expressed from pCRISPRia-v2 | protospacer |
| T7FwdVar-FBXO31.KO3 | GGATCCTAATACGACTCACTATAGATCAGGTGGATCCTGAACAGGTTTTAGAGCTAGAA | template for extension PCR to generate IVT template of sgFBXO31.KO3 | PCR template |
| T7FwdVar-FBXO31.VC1 | GGATCCTAATACGACTCACTATAGCCGTACATGCACTCCACTGGTTTTAGAGCTAGAA | template for extension PCR to generate IVT template of sgFBXO31.VC1 | PCR template |
| T7RevLong | AAAAAAGCACCGACTCGGTGCCACTTTTTCAAGTTGATAACGGACTAGCCTTATTTTAACTTGCTATTTCTAGCTCTAAAAC | primer for extension PCR of T7FwdVar oligos | PCR primer |
| T7FwdAmp | GGATCCTAATACGACTCACTATAG | primer for extension PCR of T7FwdVar oligos | PCR primer |
| T7RevAmp | AAAAAAGCACCGACTCGG | primer for extension PCR of T7FwdVar oligos | PCR primer |
| FBXO31.KO3-VC1-GT.F | CTTTCCCTACACGACGCTCTTCCGATCTTTCATCATCGGGTGGATGTAC | amplification of target sites for sgFBXO31.KO3 and sgFBXO31.VC1 with binding sites for NGS primers | PCR primer |
| FBXO31.KO3-VC1-GT.R | GGAGTTCAGACGTGTGCTCTTCCGATCTTCTGCAGCTGAGAATAGCAC | amplification of target sites for sgFBXO31.KO3 and sgFBXO31.VC1 with binding sites for NGS primers | PCR primer |
| TKOv3_PCR1_pLCKO2_fwd | GAGGGCCTATTTCCCATGATTC | PCR1 of TKOv3 sgRNA library from genomic DNA | PCR primer |
| TKOv3_PCR1_pLCKO2_rev | CAAACCCAGGGCTGCCTTGGAA | PCR1 of TKOv3 sgRNA library from genomic DNA | PCR primer |
| TS_amplicon_NGS_fwd | AATGATACGGCGACCACCGAGATCTACAC**[i5-index]**ACACTCTTTCCCTACACGACGCTCTTCCGATCT | generation of NGS libraries from genomic DNA amplicons (FBXO31.KO3-VC1-GT.F/R) | PCR primer |
| TS_amplicon_NGS_rev | CAAGCAGAAGACGGCATACGAGAT**[i7-index]**GTGACTGGAGTTCAGACGTGTGCTCTTCCGATC | generation of NGS libraries from genomic DNA amplicons (FBXO31.KO3-VC1-GT.F/R) | PCR primer |
| TKOv3_PCR2_i5_fwd | AATGATACGGCGACCACCGAGATCTACAC**[i5-index]**ACACTCTTTCCCTACACGACGCTCTTCCGATCTTTGTGGAAAGGACGAGGTACCG | PCR2 of TKOv3 sgRNA library for deep sequencing | PCR primer |
| TKOv3_PCR2_i7_rev | CAAGCAGAAGACGGCATACGAGAT**[i7-index]**GTGACTGGAGTTCAGACGTGTGCTCTTCCGATCTACTTGCTATTTCTAGCTCTAAAAC | PCR2 of TKOv3 sgRNA library for deep sequencing | PCR primer |

### General Synthetic procedures

#### Peptide and protein synthesis methods

##### Peptide synthesis reagents and solvents

Fmoc-amino acids with suitable side-chain protecting groups, HATU (1-[bis(dimethylamino)methylene]-1H-1,2,3-triazolo[4,5-b]pyridinium 3-oxide hexafluorophosphate) were purchased from Peptides International (Louisville, KY, USA), ChemImpex (Wood Dale, IL, USA) and Merck Milipore. HPLC grade $CH_3CN$ from Sigma-Aldrich was used for analytical and preparative HPLC purification. Trifluoroacetic acid for HPLC analytical and preparative HPLC purification was purchased from ABCR. DMF (> 99.8%) from Sigma-Aldrich and N-methylpyrrolidine from ABCR were directly used without further purification for solid phase peptide synthesis. Other commercially available reagents and solvents were purchased from Sigma-Aldrich (Buchs, Switzerland), Acros Organics (Geel, Belgium) and TCI Europe (Zwijndrecht, Belgium).

##### Peptide and protein conjugate characterization

High-resolution mass spectra were recorded by the Molecular and Biomolecular Analysis Service (MoBiAS) at ETH Zurich with a Bruker maXis instrument (ESI-MS measurements) equipped with an ESI source and a Q-TOF detector. Reaction monitoring was performed on a Bruker microFLEX instrument (MALDI-TOF) using 4-hydroxy-α-cyanocinnamic acid as matrix. Advanced raw data processing for bioanalytical interpretations were performed using PEAKS Studio software (Bioinformatics Solutions Inc., Canada).

##### Peptide purification

Peptides were analyzed and purified by reverse phase high performance liquid chromatography (RP-HPLC) on JASCO analytical and preparative instruments equipped with dual pump, mixed and in-line degasser, a variable wavelength UV detector (simultaneous detection of the eluent at 220 nm, 254 nm and 301 nm) and a Rheodyne injector with a 200 µL or 10 mL injection loop. Columns were heated to 60 ºC using a Jetstream 2 column heater (analytical) or a $H_2O$ water bath (preparative). The mobile-phase for RP-HPLC was Milipore-$H_2O$ containing 0.1% (v/v) TFA and HPLC grade $CH_3N$ containing 0.1% (v/v) TFA. Analytical HPLC was performed on Shiseido Capcell Pak C18 (5 µm, 4.6 mm I.D. x 250 mm) columns at a flow rate of 1 mL/min. Preparative HPLC was performed on Shiseido Capcell Pak MGIII (5 µm, 20 mm I.D. x 250 mm) at a flow rate of 40 mL/min.

General *analytical* HPLC methods:

- flow 1 mL/min, isocratic 10% $CH_3CN$ for 3 min, then gradient from 10% to 95% $CH_3CN$ in 14 min

General *preparative* HPLC methods:

- flow 40 mL/min, isocratic 5% $CH_3CN$ for 5 min, then gradient from 10% to 65% $CH_3CN$ in 28 min.

##### Solid phase peptide synthesis

Loading of amino acids on solid support was performed as followed:

- Chloro-trityl resin: The amino acid (1.20 equiv of desired loading) was dissolved in $CH_2Cl_2$ (200 mM). NMM (2 equiv) was added to the solution. The solution was given to preswollen chlorotrityl resin and shaken for 1h. The resin was washed with $CH_2Cl_2$ and DMF. Remaining chloro-trityl moieties were capped with $CH_2Cl_2$/MeOH/NMM (17:2:1, v:v:v) for 1 min. The capping step was repeated once. The resin was washed with $CH_2Cl_2$ and DMF. The resin was dried using a $N_2$ stream prior to usage.

- Rink amide resin: Fmoc-Rink amide resin was deprotected using 20 vol% piperidine in DMF for 2x5 min. The resin was washed thoroughly. Amino acid (1.2 equiv of desired loading) and HCTU (0.95 equiv of amino acid) were dissolved in DMF (200 mM). NMM (2 equiv of amino acid) was added. The solution was added to the preswollen Rink amide resin and shaken for 18 h. The resin was washed with $CH_2Cl_2$ and DMF. The resin was dried using a $N_2$ stream prior to usage.

Peptides were synthesized on a Multisyntech Syro I parallel synthesizer using Fmoc-SPPS chemistry. The following Fmoc amino acids with side-chain protection groups were used: Fmoc-Ala-OH, Fmoc-Arg(Pbf)-OH, Fmoc-Asn(Trt)-OH, Fmoc-Asp(OtBu)-OH, Fmoc-Gln(Trt)-OH, Fmoc-Glu(OtBu)-OH, Fmoc-Gly-OH, Fmoc-His(1-Trt)-OH, Fmoc-Ile-OH, Fmoc-Leu-OH, Fmoc-Lys(Boc)-OH, Boc-Lys(Fmoc)-OH Fmoc-Met-OH, Fmoc-Phe-OH, Fmoc-Pro-OH, Fmoc-Ser(tBu)-OH, Fmoc-Thr(tBu)-OH, Fmoc-Trp(Boc)-OH, Fmoc-Tyr(tBu)-OH, Fmoc-Val-OH.

General methods on Multisyntech Syro I parallel synthesizer:

- Amino acids were dissolved in DMF to a concentration of 0.5 M. HATU was dissolved in DMF to a concentration of 0.5 M. DIPEA was dissolved in NMP to a concentration of 2 M. Amino acid, HATU and DIPEA are mixed to a final concentration of 0.2 M, 0.2 M and 0.4 M, respectively, and added to the resin. The resin was agitated for 45 min. Coupling steps were repeated once.
- Capping was performed with acetic anhydride. 20 vol% acetic anhydride in DMF was mixed with 2 M DIPEA at a ratio of 3:2 and added to the resin. The resin was agitated for 5 min. The capping step was repeated once.
- Fmoc deprotection was performed with 20 vol% piperidine in DMF for 10 min. The deprotection step was repeated once.

Preparation of Fmoc-arginol (Pbf)

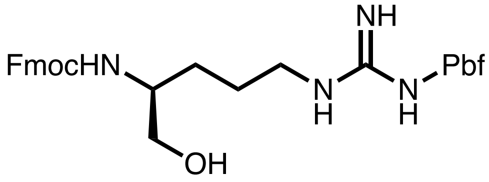

Fmoc-Arg(Pbf)-OH (2.00 g, 3.10 mmol) was dissolved in dimethoxyethane (20 mL) and cooled to –15 ºC using an ice-salt bath under inert atmosphere. N-methyl morpholine (440 µL, 3.40 mmol) was added followed by isobutyl-chloroformate (373 µL, 3.40 mmol) and stirred for 30 min. After 30 min the reaction was filtered and the filtrate was immediately cooled to –15 ºC using an ice-salt bath under inert atmosphere. $NaBH_4$ (350 mg, 9.25 mmol) was added in one portion and stirred for 15 min. The reaction was quenched by addition of saturated aqueous $NaHCO_3$. The mixture was extracted with EtOAc (3x). The organics were washed with 0.5 M HCl, brine, dried using $MgSO_4$ and concentrated to provide Fmoc-arginol-(Pbf) (1.9 g) as a white foam. The product was used without further purification.

HRMS (ESI): calculated for $[C_{34}H_{43}N_4O_6S]^+$: m/z 635.2898, found: m/z 635.2890
**$^1$H-NMR** (500 MHz, $CDCl_3$) d 7.73 (2H, d), 7.57 (2H, d), 7.37 (2H, t), 7.25 (2H, m), 6.30 (3H, brs), 5.64 (1H, brs), 4.36 (2H, d), 4.15 (1H, t), 3.71-3.53 (3H, m), 3.23 (2H, brs), 2.91 (2H, s), 2.58 (3H, s), 2.51 (3H, s), 2.08 (3H, s), 1.67-1.49 (4H, m), 1.44 (6H, s)
**$^{13}$C-NMR** (125 MHz, $CDCl_3$) d 159.07, 157.13, 156.39, 144.01, 143.93, 141.39, 138.55, 132.46, 127.82, 127.20, 125.25, 124.89, 120.07, 117.79, 86.62, 66.84, 64.83, 47.34, 43.30, 41.18, 28.69, 25.70, 19.45, 18.10, 12.61

**Synthesis of fluorescent peptides for fluorescence polarization measurement**

General method

Amino acids were loaded on chlorotrityl resin to access C-terminal carboxylic acids or Rink amide to access C-terminal amides according to the general methods. Automated peptide elongation was carried out on a Multisyntech Syro I parallel synthesizer according to the general peptide methods. N-terminal Boc-Lysine(Fmoc)-OH was coupled manually (2 equiv, 90 min). Fmoc group was removed by treatment with 20 vol% piperidine. The resin was washed thoroughly. All following steps were performed in the dark. FITC (3 equiv) and NMM (6 equiv) were dissolved in DMF, given to the resin and shaken for 2 h. The peptide was cleaved from the resin using TFA/DODT/$H_2O$ (95:2.5:2.5, v/v) for 1 h. The resin was removed by filtration and the filtrate concentrated under reduced pressure. The solution was triturated with $Et_2O$ and centrifuged to obtain crude peptide. The crude peptide was dissolved in $H_2O/CH_3N$ (1:1, v/v) + 0.1% (v/v) TFA and purified using preparative HPLC unless stated otherwise.

Preparation of peptide alcohol

Fmoc-arginol (Pbf) (190 mg, 0.3 mmol) in 2 vol% DBU in DMF was added to 2-chlorotrityl chloride resin (0.1 mmol) and shaken overnight. The resin was washed with DMF and $CH_2Cl_2$ and capped according to the general procedure. Fmoc-Leu (212 mg, 0.6 mmol) was dissolved in DMF/$CH_2Cl_2$ (1:1, v/v) at 0 ºC. DIC (47 µL, 0.3 mmol) and DMAP (3 mg) were added and then given to the resin. The resin was shaken for 6 h and the coupling was repeated two additional times. Following standard Fmoc-SPPS, Boc-Lys(Fmoc)-OH and FITC were coupled as described above. Following resin cleavage, the crude peptide was dissolved in PBS 1 M $NaHCO_3$ pH 8.0 was added and the pH adjusted to 8.0. The solution as incubated for 2 h at room temperature, acidified using TFA and purified by preparative RP-HPLC.

##### 1.1.1. Preparation of peptide methyl-ester

Preparation of peptide methyl-ester

Cyanosulfurylide resin was prepared according to previously reported procedures and Fmoc-Arg(Pbf)-OH was loaded. Following standard Fmoc-SPPS, Boc-Lys(Fmoc)-OH and FITC were coupled as described above. The resin was thoroughly washed with DMF and $CH_2Cl_2$ and dried. The resin was resuspended in THF/MeOH (1:1, v/v) and N-chlorosuccinimide (40 mg) was added and the mixture incubated for 20 min. The filtrate was removed, and the resin washed with THF. The filtrate was concentrated and standard cleavage cocktail was added. The solution was triturated with $Et_2O$ and centrifuged to obtain crude peptide. The crude peptide was dissolved in $H_2O/CH_3N$ (1:1, v/v) + 0.1% (v/v) TFA and purified using preparative HPLC.

**Peptide library synthesis**

Isokinetic amino acid mixture

Ala (3.4 mol%), Arg (6.5 mol%), Asn (5.3 mol%), Asp (3.5 mol%), Gln (5.3 mol%), Glu (3.6 mol%), Gly (2.9 mol%), His (3.5 mol%), Ile (17.4 mol%), Leu (4.9 mol%), Lys (6.2 mol%), Met (3.8 mol%), Phe (2.5 mol%), Pro (4.3 mol%), Ser (2.8 mol%), Thr (4.8 mol%), Trp (3.8 mol%), Tyr (4.1 mol%), Val (11.3 mol%)

Amino acids were dissolved in DMF to a total concentration of 0.5 M and used as is for resin loading and peptide elongation steps.

Synthesis of three variable peptide library with C-terminal carboxylic acid

**GGGKYRYDVPDYXXX-COOH**

Resin loading and peptide synthesis was performed as described in the general methods using the isokinetic amino acid mixture for positions X. Automated peptide elongation was carried out on a Multisyntech Syro I parallel synthesizer according to the general peptide methods. The peptide was cleaved form the resin using TFA/DODT/$H_2O$ (95:2.5:2.5, v/v) for 1 h. The resin was removed by filtration and the filtrate concentrated under reduced pressure. The solution was triturated with $Et_2O$ and centrifuged to obtain crude peptide. $Et_2O$ was repeated three times. The crude peptide was dissolved in $H_2O/CH_3N$ (1:1, v/v) and lyophilized to obtain an off-white powder.

Synthesis of three variable peptide library with C-terminal amide

**GGGKYRYDVPDYXXX-CONH$_2$**

Resin loading and peptide synthesis was performed as described in the general methods using the isokinetic amino acid mixture for positions X. Automated peptide elongation was carried out on a Multisyntech Syro I parallel synthesizer according to the general peptide methods. The peptide was cleaved form the resin using TFA/DODT/$H_2O$ (95:2.5:2.5, v/v) for 1 h. The resin was removed by filtration and the filtrate concentrated under reduced pressure. The solution was triturated with $Et_2O$ and centrifuged to obtain crude peptide. $Et_2O$ was repeated three times. The crude peptide was dissolved in $H_2O/CH_3N$ (1:1, v/v) and lyophilized to obtain an off-white powder.

### Individual QC of synthesis products

#### Characterization of fluorescein-conjugated SPPS products

Synthesis of fluorescein-pep2–R–COOH

**Fluorescein–KEEDEKGSRASDDFRDLR–COOH**

The peptide was obtained as a yellow-orange solid.
HRMS (ESI): calculated for $[C_{108}H_{154}N_{30}O_{40}S]^{2+}$: m/z 1271.5324, found: m/z 1271.5347

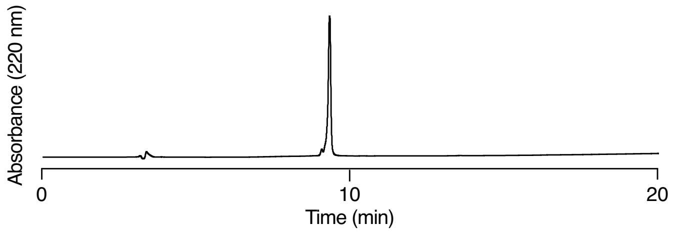

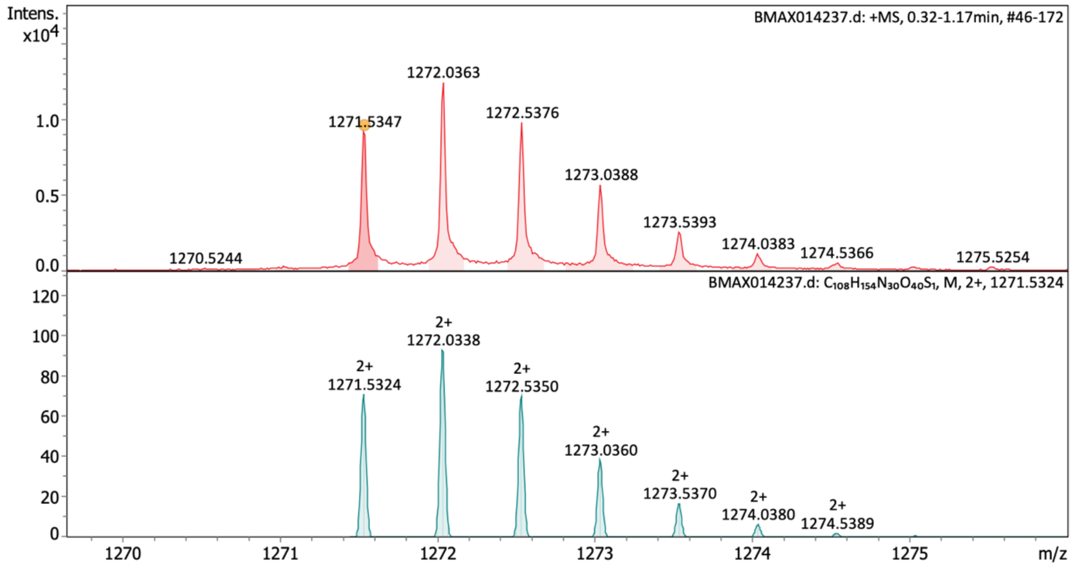

**Characterization of fluorescein-pep2–R–COOH** Analytical RP-HPLC of purified peptide. HRMS (ESI) spectrum of purified peptide showing recorded mass spectrum (upper panel) and calculated spectrum (lower panel).

Synthesis of fluorescein-pep2–R–$CONH_2$

**Fluorescein–KEEDEKGSRASDDFRDLR–$CONH_2$**

The peptide was obtained as a yellow-orange solid.
HRMS (ESI): calculated for $[C_{108}H_{155}N_{31}O_{39}S]^{2+}$: m/z 1271.0404, found: m/z 1271.0423

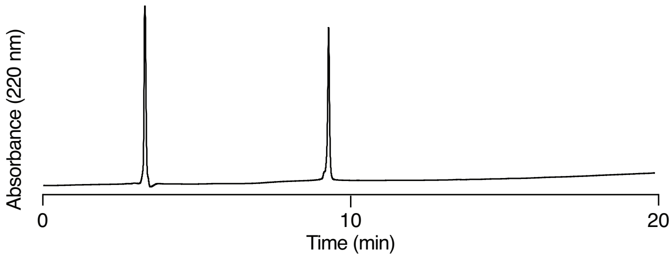

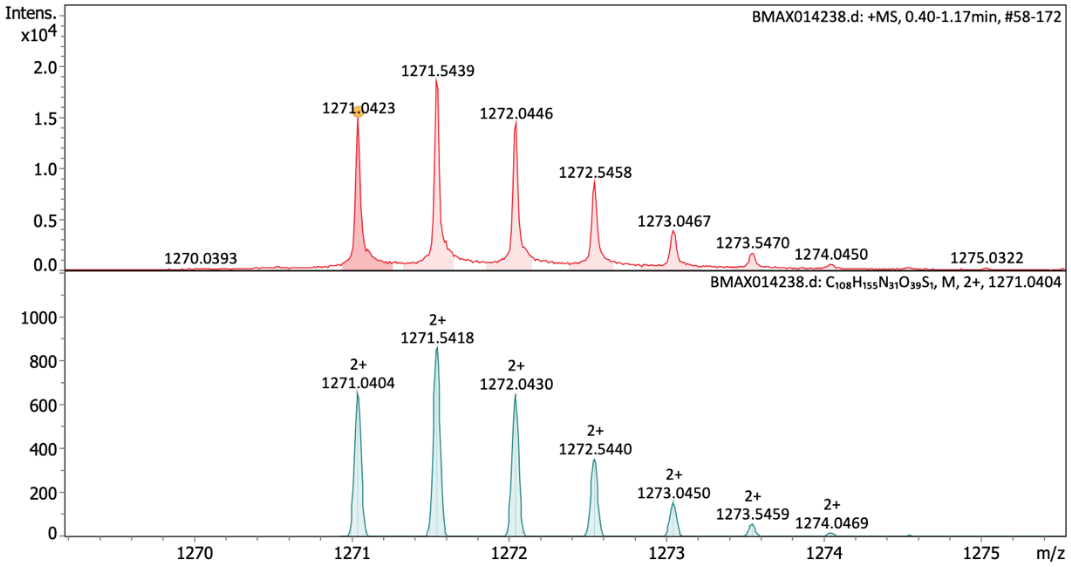

**Characterization of fluorescein-pep2–R–$CONH_2$** Analytical RP-HPLC of purified peptide. HRMS (ESI) spectrum of purified peptide showing recorded mass spectrum (upper panel) and calculated spectrum (lower panel).

Synthesis of fluorescein-KAA-CycD1(286-295)–COOH

**Fluorescein– KAATPTDVRDVDI–COOH**

The peptide was obtained as a yellow-orange solid.
HRMS (ESI): calculated for $[C_{80}H_{114}N_{18}O_{27}S]^{2+}$: m/z 895.3905, found: m/z 895.3916

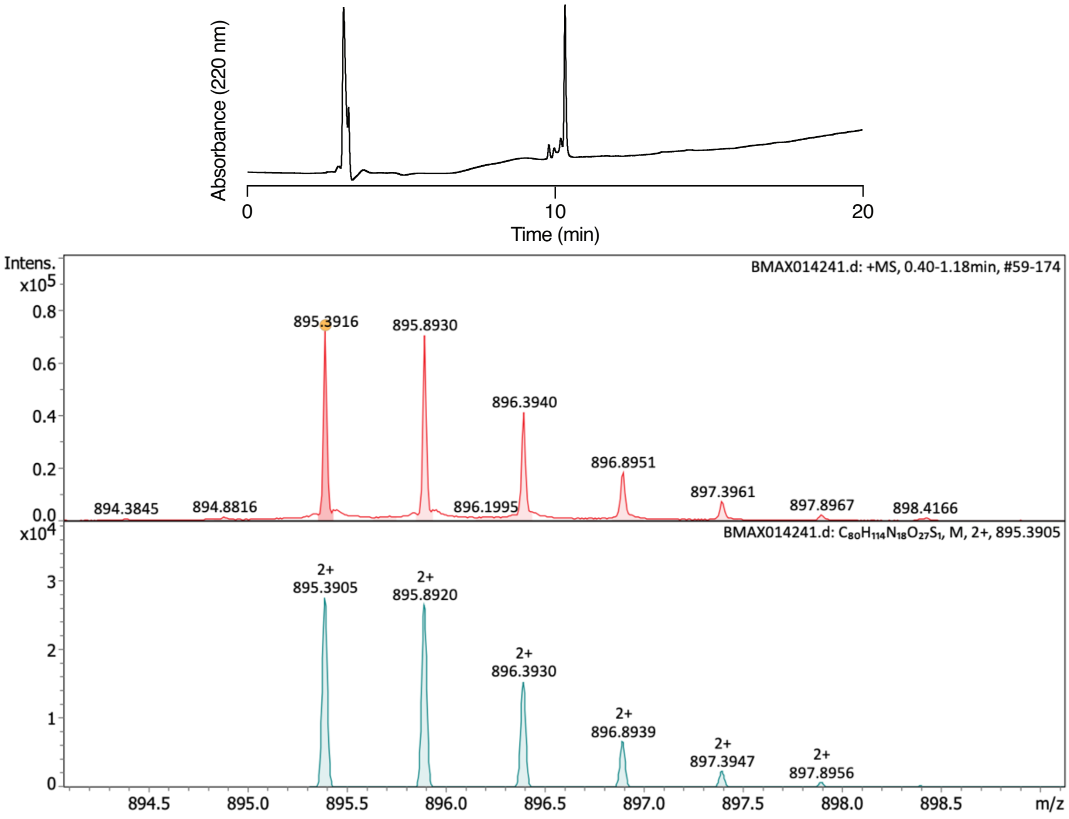

**Characterization of fluorescein-KAA-CycD1(286-295)–COOH** Analytical RP-HPLC of purified peptide. HRMS (ESI) spectrum of purified peptide showing recorded mass spectrum (upper panel) and calculated spectrum (lower panel).

Synthesis of fluorescein-KAA-CycD1(286-295)–$CONH_2$

**Fluorescein– KAATPTDVRDVDI–$CONH_2$**

The peptide was obtained as a white solid.
HRMS (ESI): calculated for $[C_{80}H_{115}N_{19}O_{26}S]^{2+}$: m/z 894.8985, found: m/z 894.901

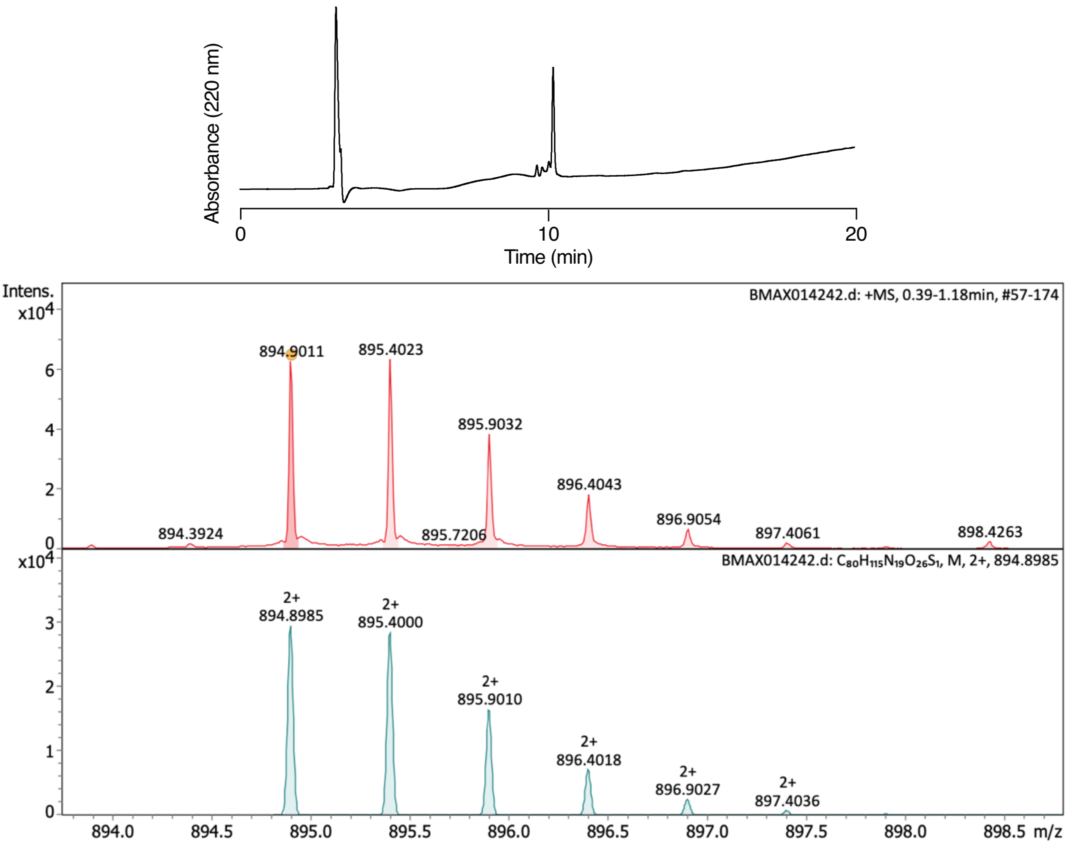

**Characterization of fluorescein-KAA-CycD1(286-295)–$CONH_2$** Analytical RP-HPLC of purified peptide. HRMS (ESI) spectrum of purified peptide showing recorded mass spectrum (upper panel) and calculated spectrum (lower panel).

Synthesis of fluorescein-pep3–A–$CONH_2$

**Fluorescein-KKYRYDVPDYSAA–$CONH_2$**

The peptide was obtained as an orange solid and used without preparative RP-HPLC purification. HRMS (ESI): calculated for $[C_{93}H_{120}N_{20}O_{26}S_1]^{2+}$: m/z 982.4196, found: m/z 982.4212

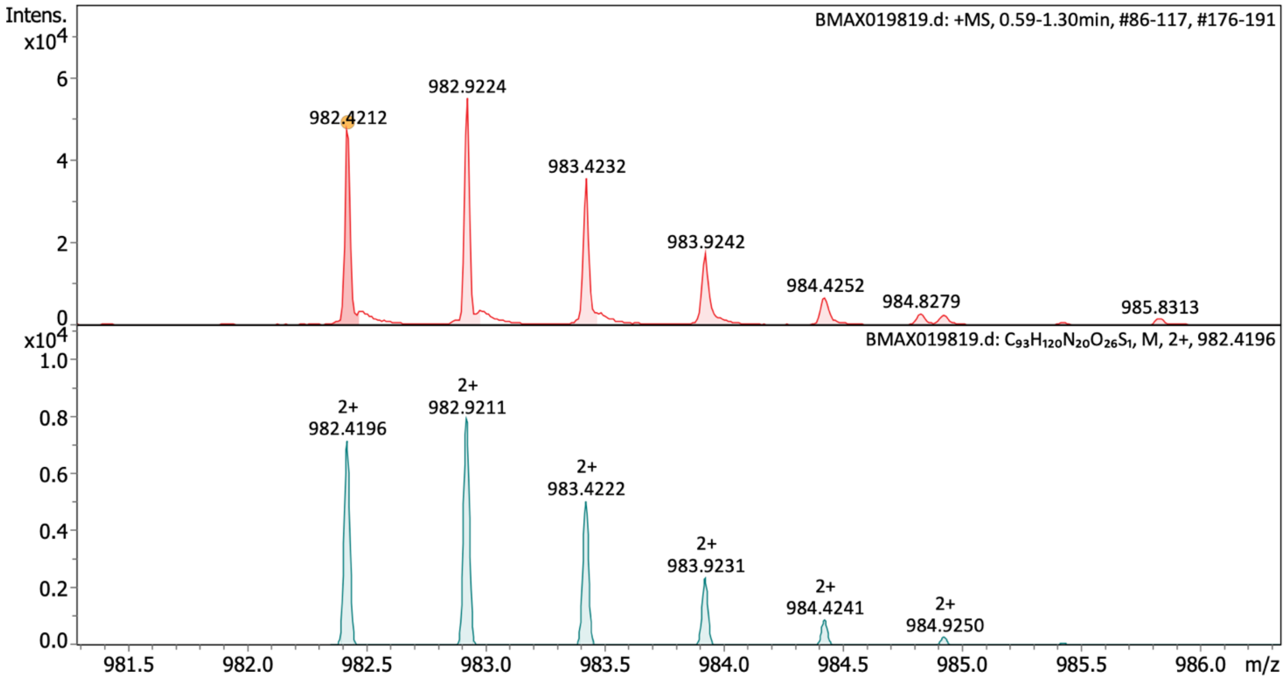

**Characterization of fluorescein-pep3–A–$CONH_2$** HRMS (ESI) spectrum of purified peptide showing recorded mass spectrum (upper panel) and calculated spectrum (lower panel).

Synthesis of fluorescein-pep3–C–$CONH_2$

**Fluorescein-KKYRYDVPDYSAC–$CONH_2$**

The peptide was obtained as an orange solid and used without preparative RP-HPLC purification. HRMS (ESI): calculated for $[C_{93}H_{120}N_{20}O_{26}S_2]^{2+}$: m/z 988.4057, found: m/z 988.4070

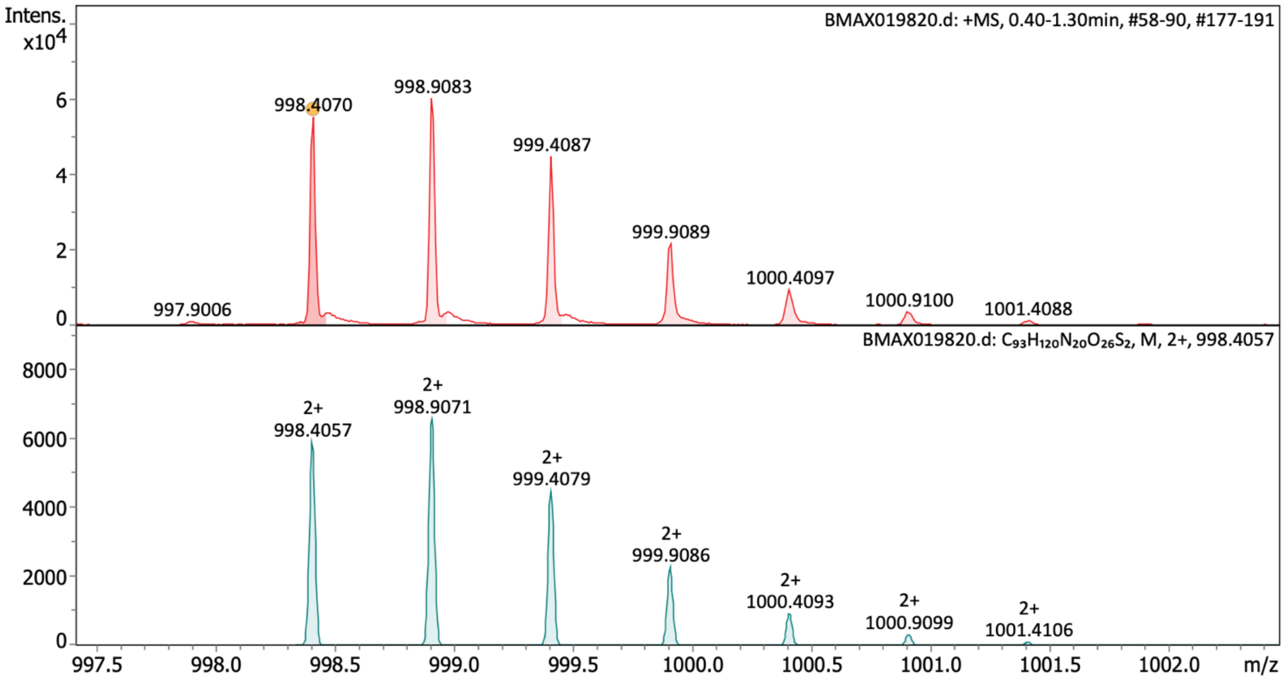

**Characterization of fluorescein-pep3–C–$CONH_2$** HRMS (ESI) spectrum of purified peptide showing recorded mass spectrum (upper panel) and calculated spectrum (lower panel).

Synthesis of fluorescein-pep3–D–$CONH_2$

**Fluorescein-KKYRYDVPDYSAD–$CONH_2$**

The peptide was obtained as an orange solid and used without preparative RP-HPLC purification. HRMS (ESI): calculated for $[C_{94}H_{120}N_{20}O_{28}S_1]^{2+}$: m/z 1004.4145, found: m/z 1004.4164

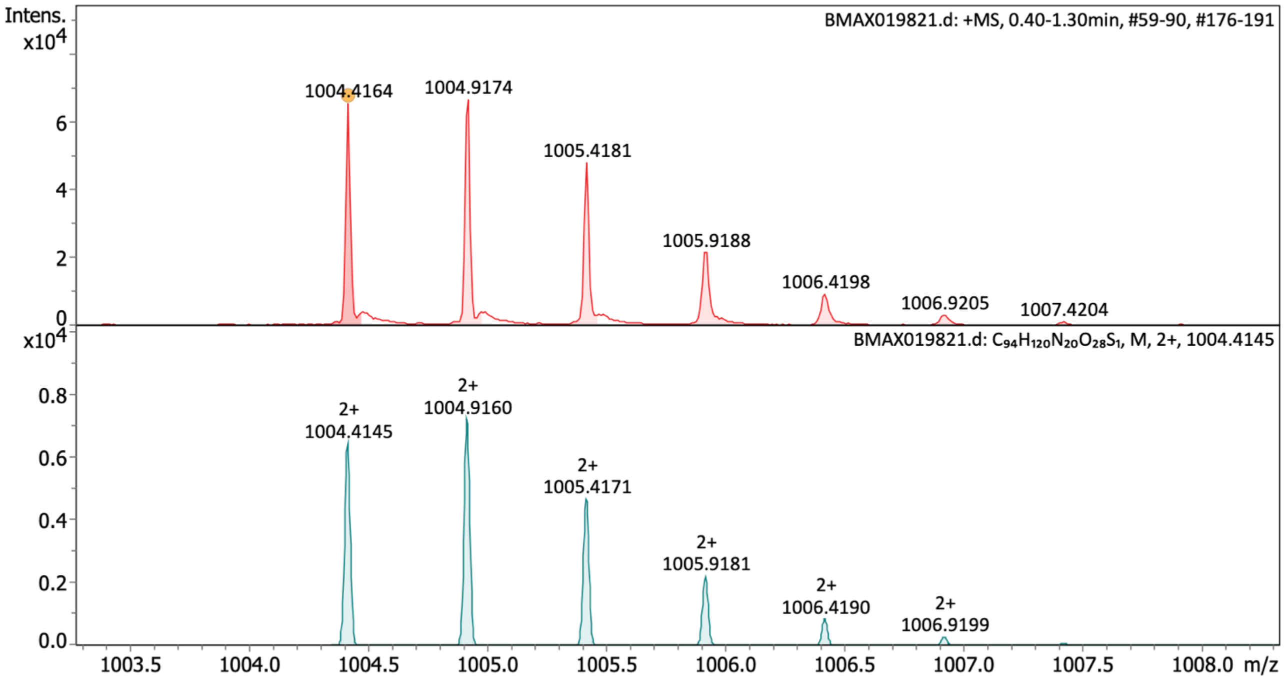

**Characterization of fluorescein-pep3–D–$CONH_2$** HRMS (ESI) spectrum of purified peptide showing recorded mass spectrum (upper panel) and calculated spectrum (lower panel).

Synthesis of fluorescein-pep3–E–$CONH_2$

**Fluorescein-KKYRYDVPDYSAE–$CONH_2$**

The peptide was obtained as an orange solid and used without preparative RP-HPLC purification. HRMS (ESI): calculated for $[C_{95}H_{122}N_{20}O_{28}S_1]^{2+}$: m/z 1011.4224, found: m/z 1011.4241

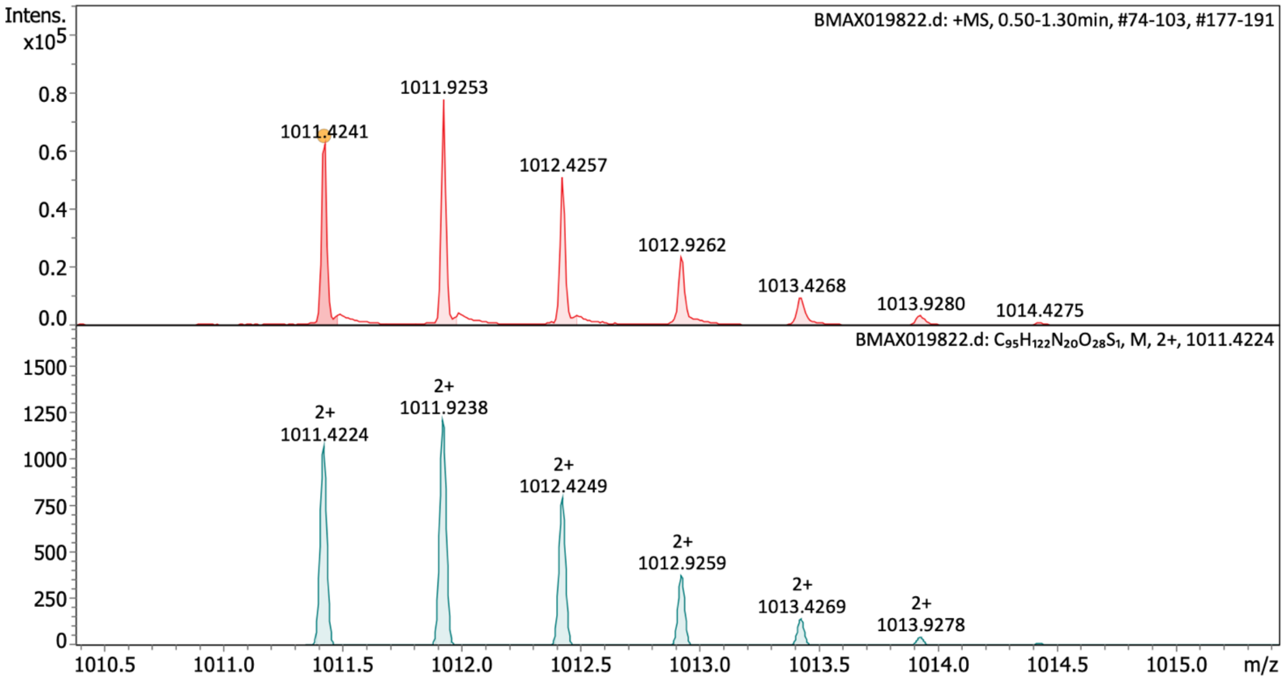

**Characterization of fluorescein-pep3–E–$CONH_2$** HRMS (ESI) spectrum of purified peptide showing recorded mass spectrum (upper panel) and calculated spectrum (lower panel).

Synthesis of fluorescein-pep3–F–$CONH_2$

**Fluorescein-KKYRYDVPDYSAF–$CONH_2$**

The peptide was obtained as an orange solid and used without preparative RP-HPLC purification. HRMS (ESI): calculated for $[C_{99}H_{124}N_{20}O_{26}S_1]^{2+}$: m/z 1020.4353, found: m/z 1020.4373

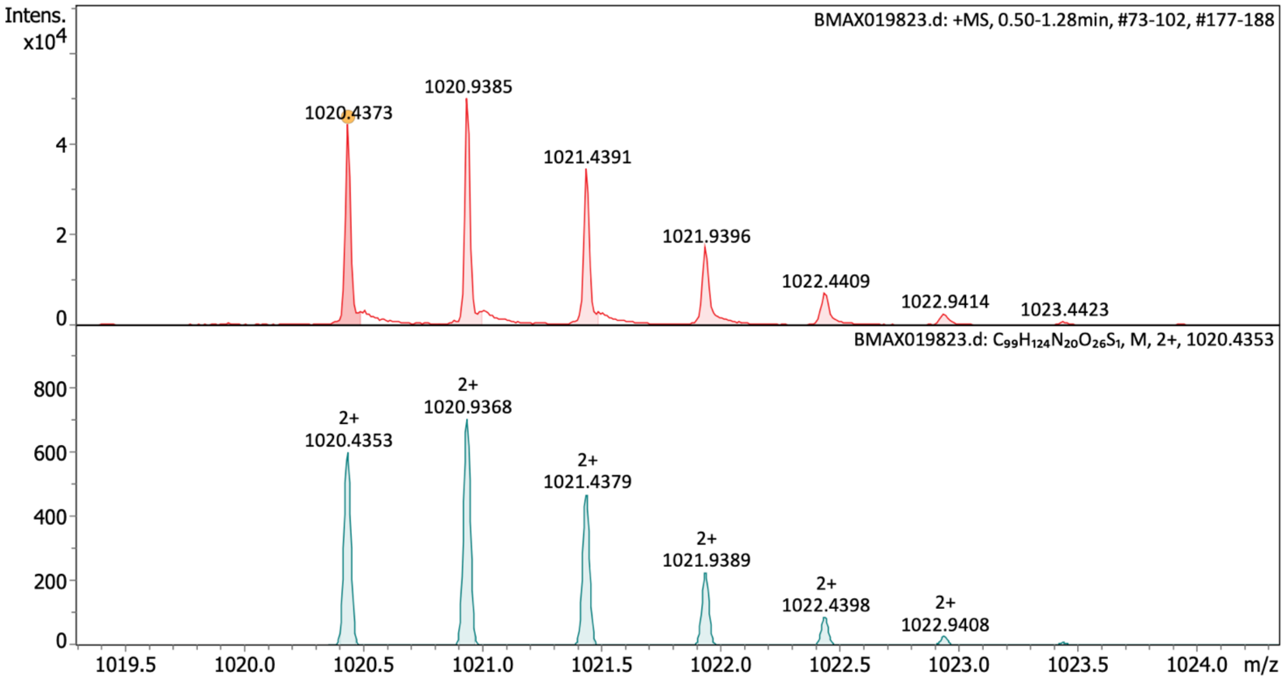

**Characterization of fluorescein-pep3–F–$CONH_2$** HRMS (ESI) spectrum of purified peptide showing recorded mass spectrum (upper panel) and calculated spectrum (lower panel).

Synthesis of fluorescein-pep3–G–$CONH_2$

**Fluorescein-KKYRYDVPDYSAG–$CONH_2$**

The peptide was obtained as an orange solid and used without preparative RP-HPLC purification. HRMS (ESI): calculated for $[C_{92}H_{118}N_{20}O_{26}S_1]^{2+}$: m/z 975.4118, found: m/z 975.4140

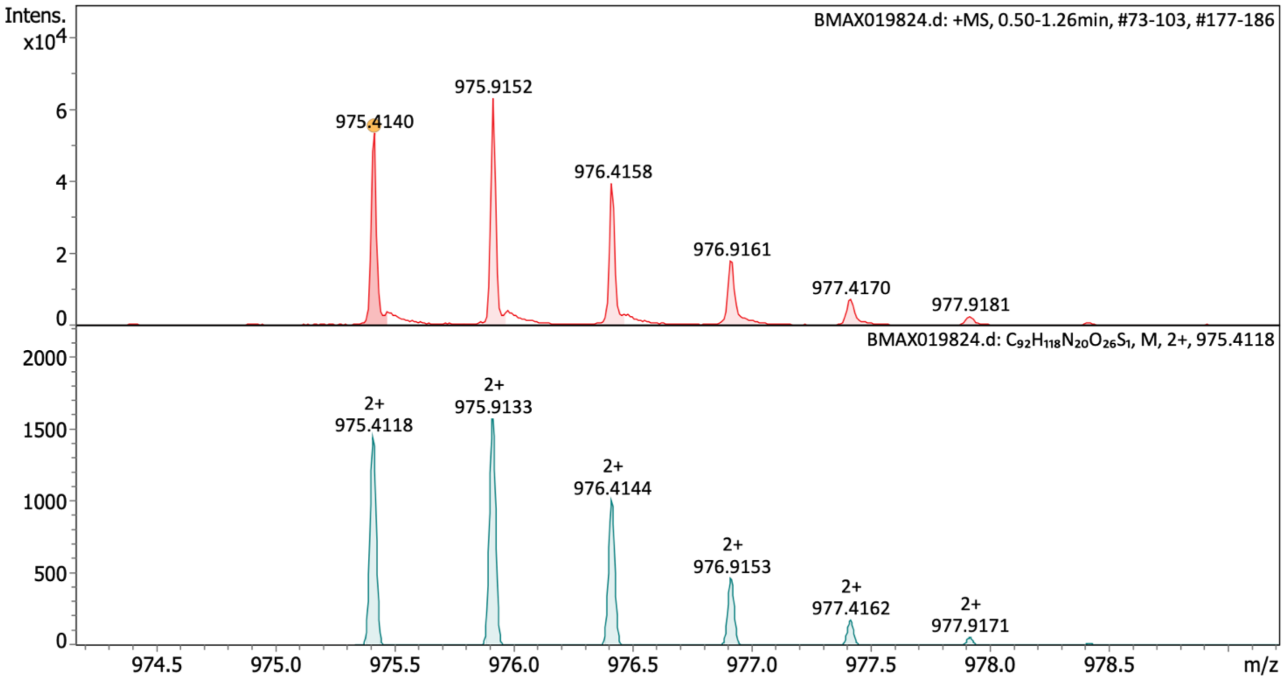

**Characterization of fluorescein-pep3–G–$CONH_2$** HRMS (ESI) spectrum of purified peptide showing recorded mass spectrum (upper panel) and calculated spectrum (lower panel).

Synthesis of fluorescein-pep3–H–CONH$_2$

**Fluorescein-KKYRYDVPDYSAH–CONH$_2$**

The peptide was obtained as an orange solid and used without preparative RP-HPLC purification. HRMS (ESI): calculated for $[C_{96}H_{122}N_{22}O_{26}S_1]^{2+}$: m/z 1015.4305, found: m/z 1015.4316

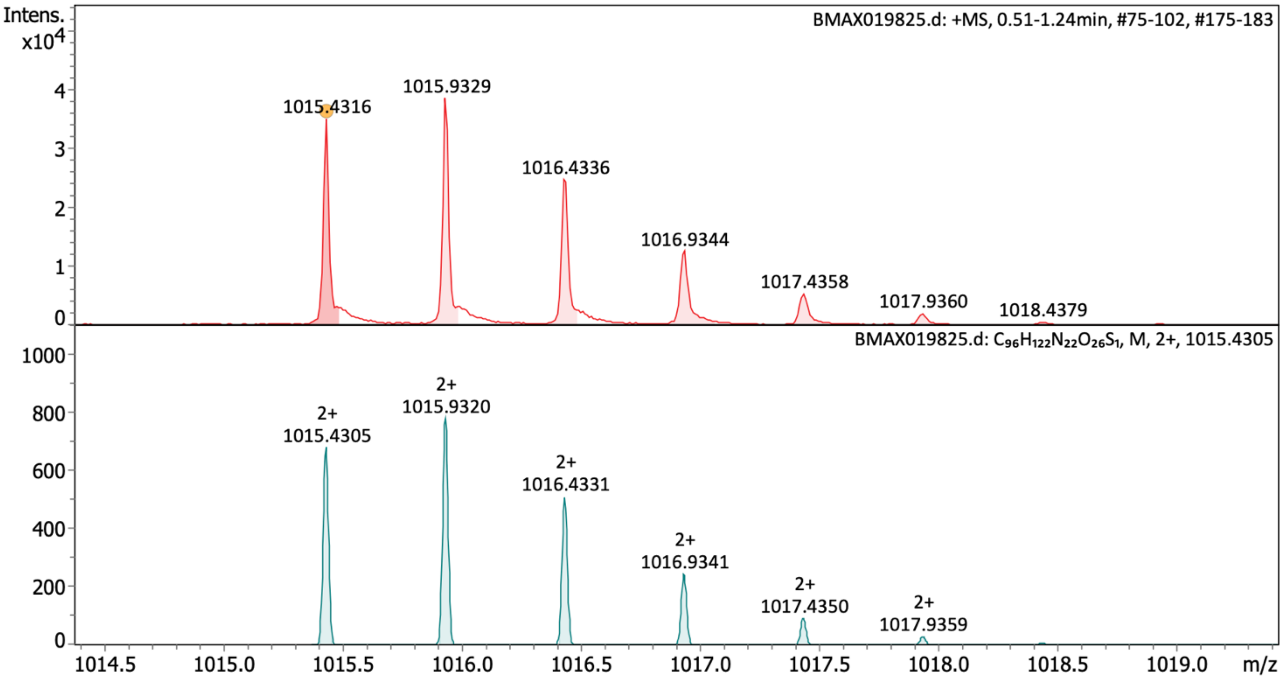

**Characterization of fluorescein-pep3–H–CONH$_2$** HRMS (ESI) spectrum of purified peptide showing recorded mass spectrum (upper panel) and calculated spectrum (lower panel).

Synthesis of fluorescein-pep3–I–$CONH_2$

**Fluorescein-KKYRYDVPDYSAI–$CONH_2$**

The peptide was obtained as an orange solid and used without preparative RP-HPLC purification. HRMS (ESI): calculated for $[C_{96}H_{126}N_{20}O_{26}S_1]^{2+}$: m/z 1003.4431, found: m/z 1003.4445

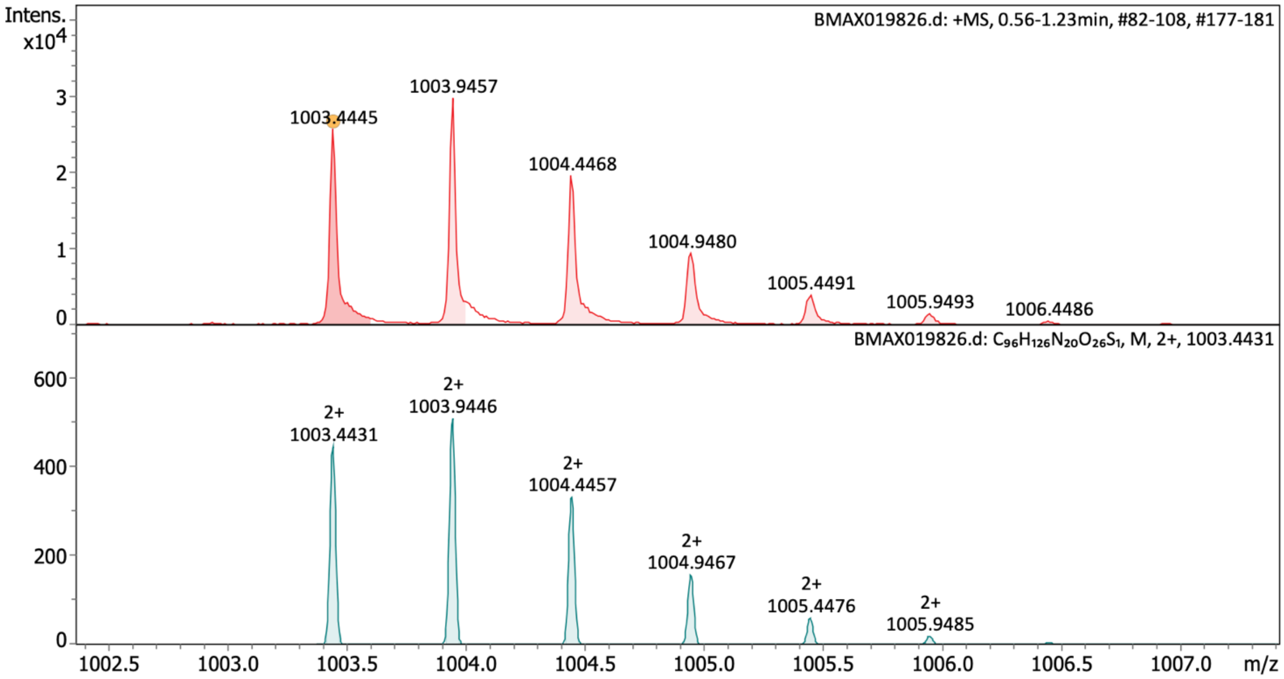

**Characterization of fluorescein-pep3–I–$CONH_2$** HRMS (ESI) spectrum of purified peptide showing recorded mass spectrum (upper panel) and calculated spectrum (lower panel).

Synthesis of fluorescein-pep3–K–$CONH_2$

**Fluorescein-KKYRYDVPDYSAK–$CONH_2$**

The peptide was obtained as an orange solid and used without preparative RP-HPLC purification. HRMS (ESI): calculated for $[C_{96}H_{127}N_{21}O_{26}S_1]^{2+}$: m/z 1010.9485, found: m/z 1010.9484

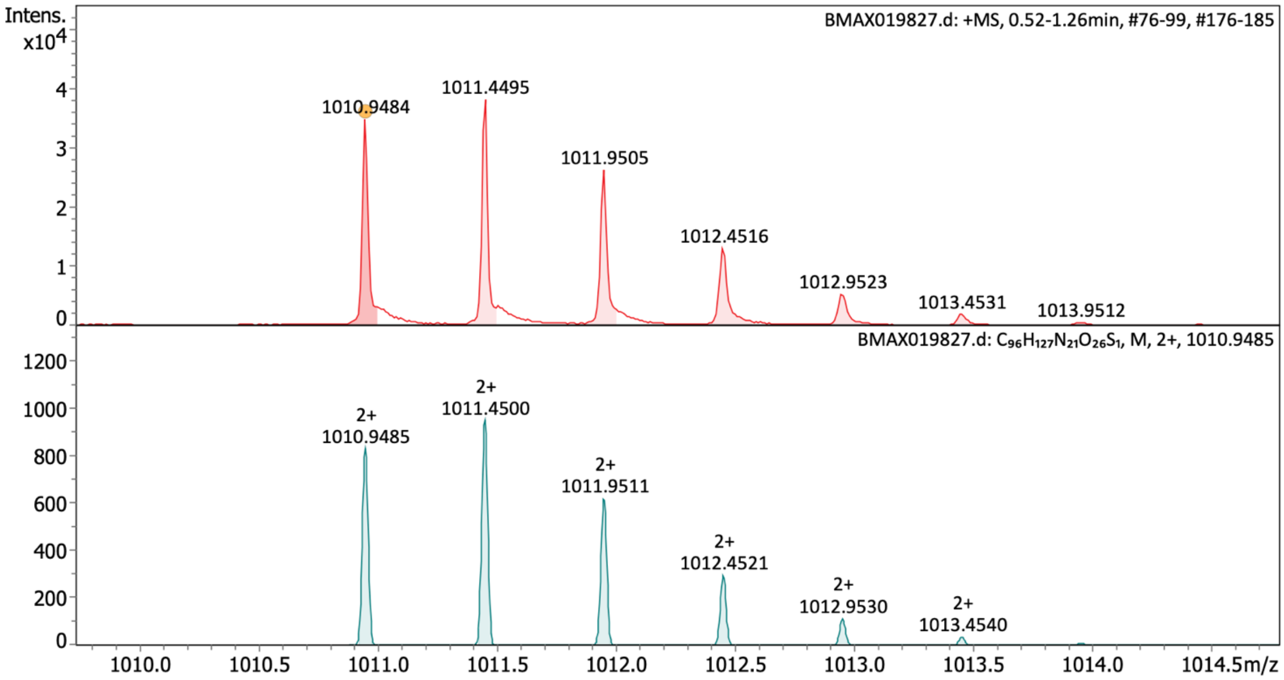

**Characterization of fluorescein-pep3–K–$CONH_2$** HRMS (ESI) spectrum of purified peptide showing recorded mass spectrum (upper panel) and calculated spectrum (lower panel).

Synthesis of fluorescein-pep3–L–$CONH_2$

**Fluorescein-KKYRYDVPDYSAL–$CONH_2$**

The peptide was obtained as an orange solid and used without preparative RP-HPLC purification. HRMS (ESI): calculated for $[C_{96}H_{126}N_{20}O_{26}S_1]^{2+}$: m/z 1003.4431, found: m/z 1003.4442

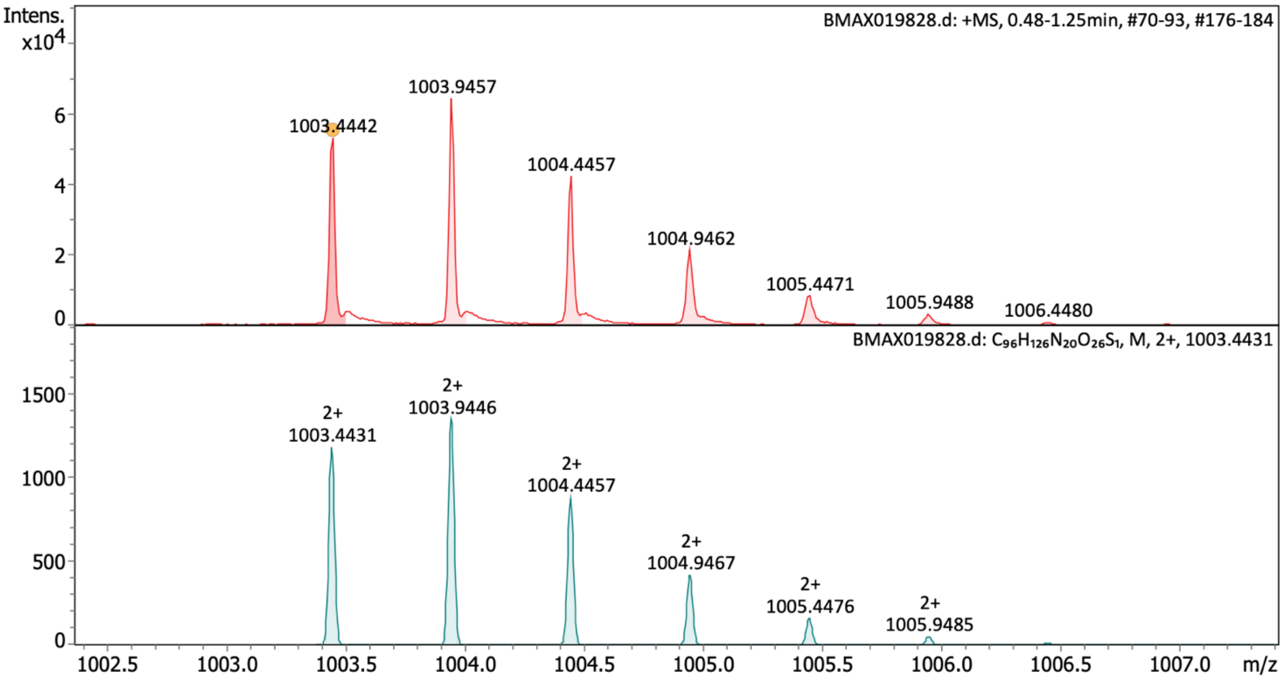

**Figure 14 Characterization of fluorescein-pep3–L–$CONH_2$** HRMS (ESI) spectrum of purified peptide showing recorded mass spectrum (upper panel) and calculated spectrum (lower panel).

Synthesis of fluorescein-pep3–M–$CONH_2$

**Fluorescein-KKYRYDVPDYSAM–$CONH_2$**

The peptide was obtained as an orange solid and used without preparative RP-HPLC purification. HRMS (ESI): calculated for $[C_{95}H_{124}N_{20}O_{26}S_2]^{2+}$: m/z 1012.4213, found: m/z 1012.4243

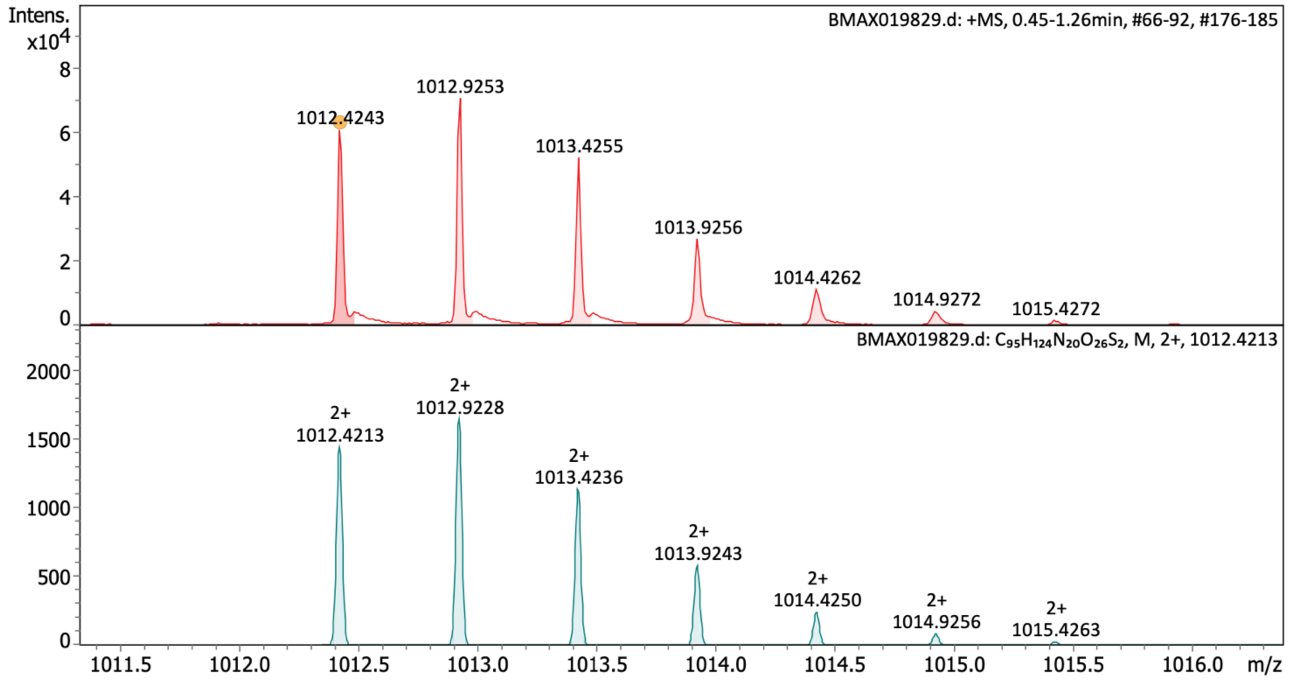

**Characterization of fluorescein-pep3–M–$CONH_2$** HRMS (ESI) spectrum of purified peptide showing recorded mass spectrum (upper panel) and calculated spectrum (lower panel).

Synthesis of fluorescein-pep3–N–$CONH_2$

**Fluorescein-KKYRYDVPDYSAN–$CONH_2$**

The peptide was obtained as an orange solid and used without preparative RP-HPLC purification. HRMS (ESI): calculated for $[C_{94}H_{121}N_{21}O_{27}S_1]^{2+}$: m/z 1003.9225, found: m/z 1003.9235

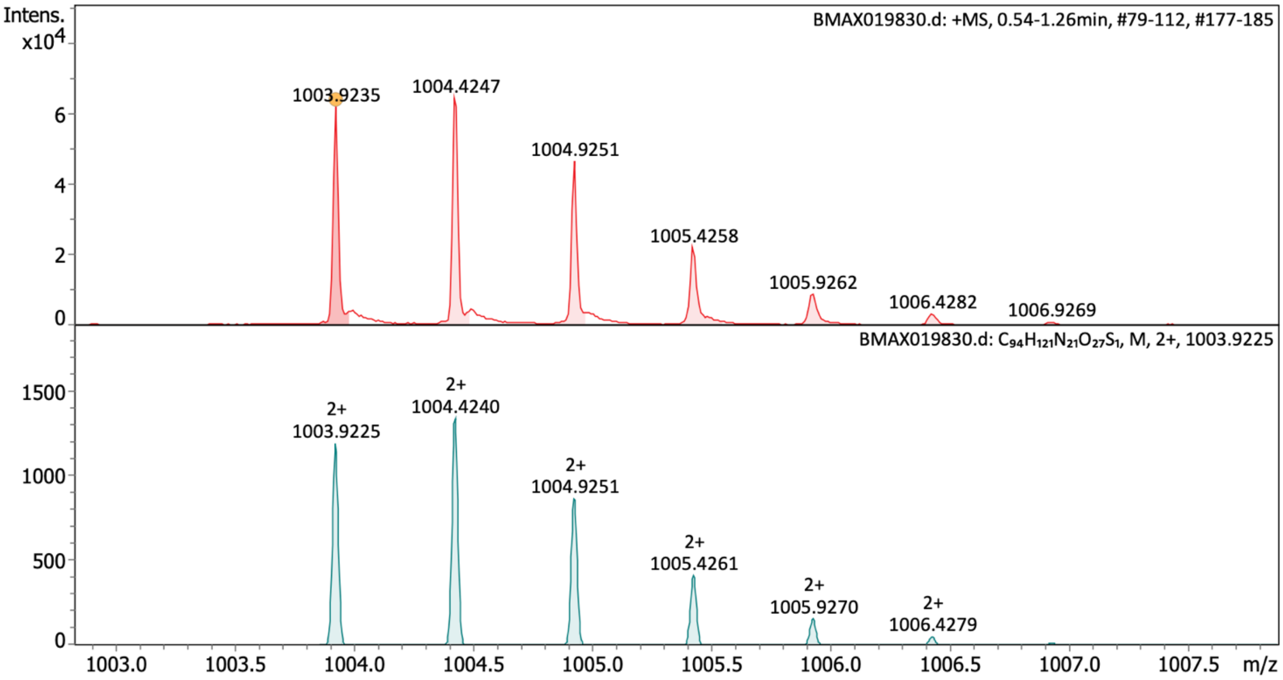

**Characterization of fluorescein-pep3–N–$CONH_2$** HRMS (ESI) spectrum of purified peptide showing recorded mass spectrum (upper panel) and calculated spectrum (lower panel).

Synthesis of fluorescein-pep3–P–$CONH_2$

**Fluorescein-KKYRYDVPDYSAP–$CONH_2$**

The peptide was obtained as an orange solid and used without preparative RP-HPLC purification. HRMS (ESI): calculated for $[C_{95}H_{122}N_{20}O_{26}S_1]^{2+}$: m/z 995.4274, found: m/z 995.4289

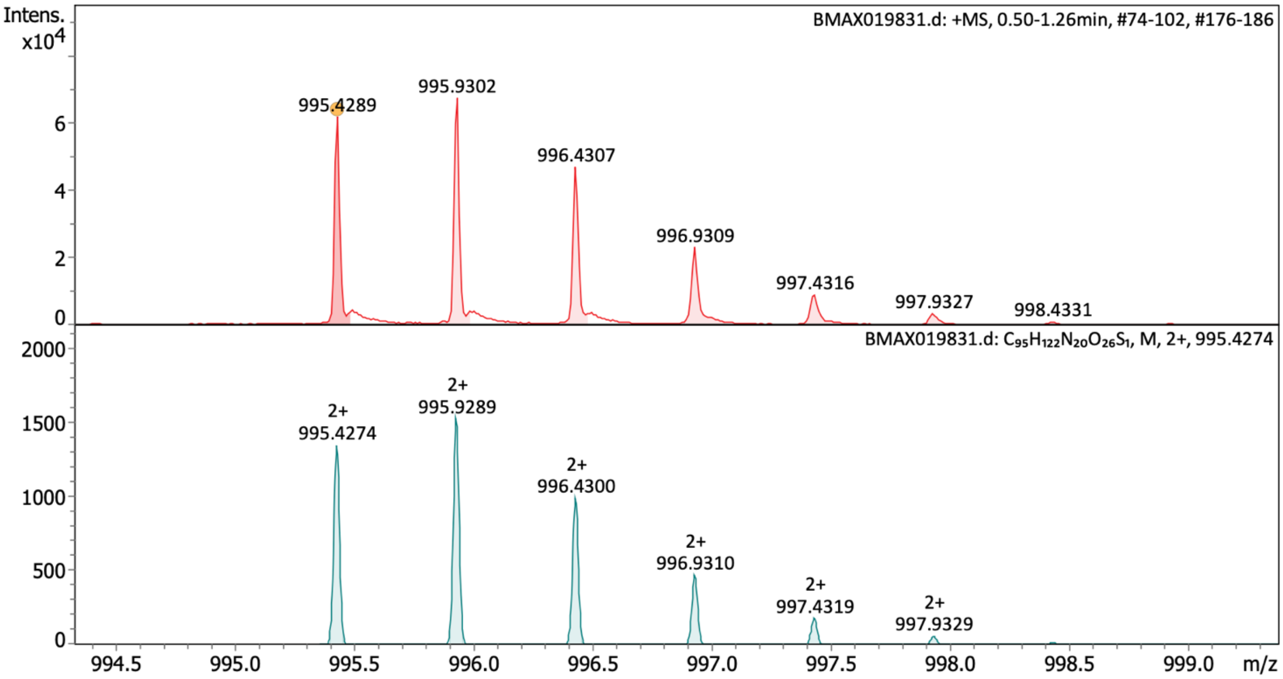

**Characterization of fluorescein-pep3–P–$CONH_2$** HRMS (ESI) spectrum of purified peptide showing recorded mass spectrum (upper panel) and calculated spectrum (lower panel).

Synthesis of fluorescein-pep3–Q–$CONH_2$

**Fluorescein-KKYRYDVPDYSAQ–$CONH_2$**

The peptide was obtained as an orange solid and used without preparative RP-HPLC purification. HRMS (ESI): calculated for $[C_{95}H_{124}N_{21}O_{27}S_1]^{3+}$: m/z 674.2893, found: m/z 674.2893

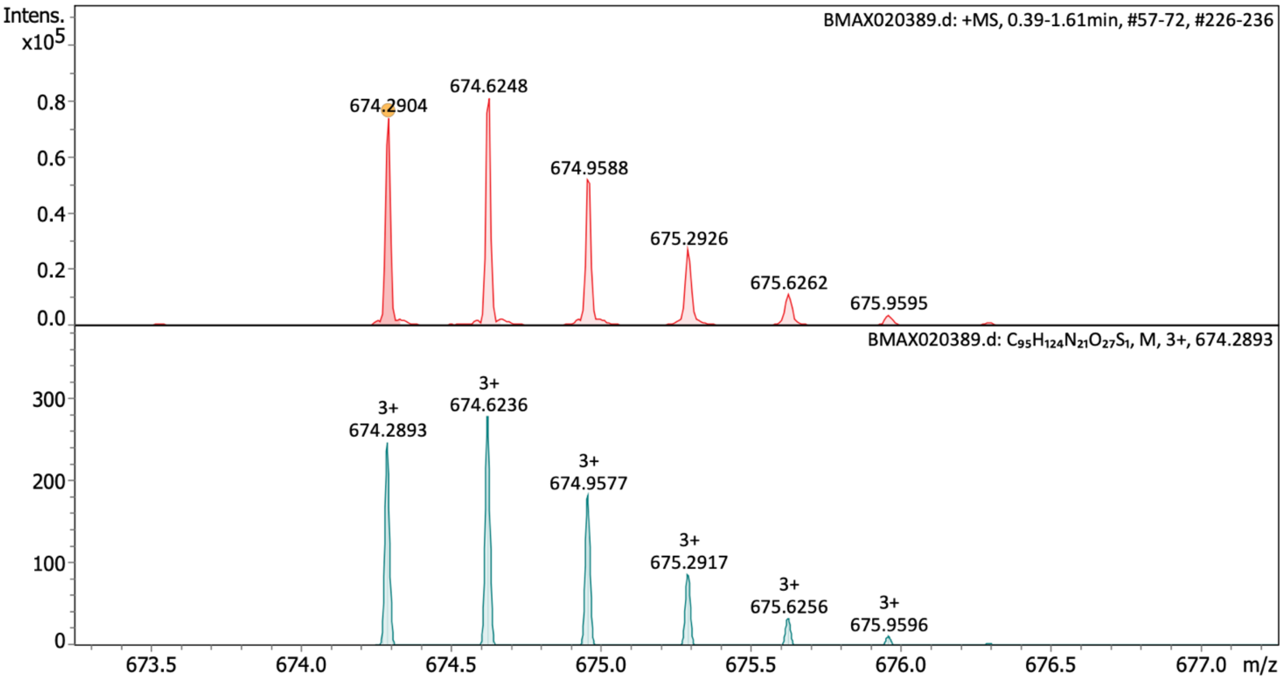

**Characterization of fluorescein-pep3–Q–$CONH_2$** HRMS (ESI) spectrum of purified peptide showing recorded mass spectrum (upper panel) and calculated spectrum (lower panel).

Synthesis of fluorescein-pep3–R–$CONH_2$

**Fluorescein-KKYRYDVPDYSAR–$CONH_2$**

The peptide was obtained as an orange solid and used without preparative RP-HPLC purification. HRMS (ESI): calculated for $[C_{96}H_{127}N_{23}O_{26}S_1]^{2+}$: m/z 1024.9516, found: m/z 1024.9523

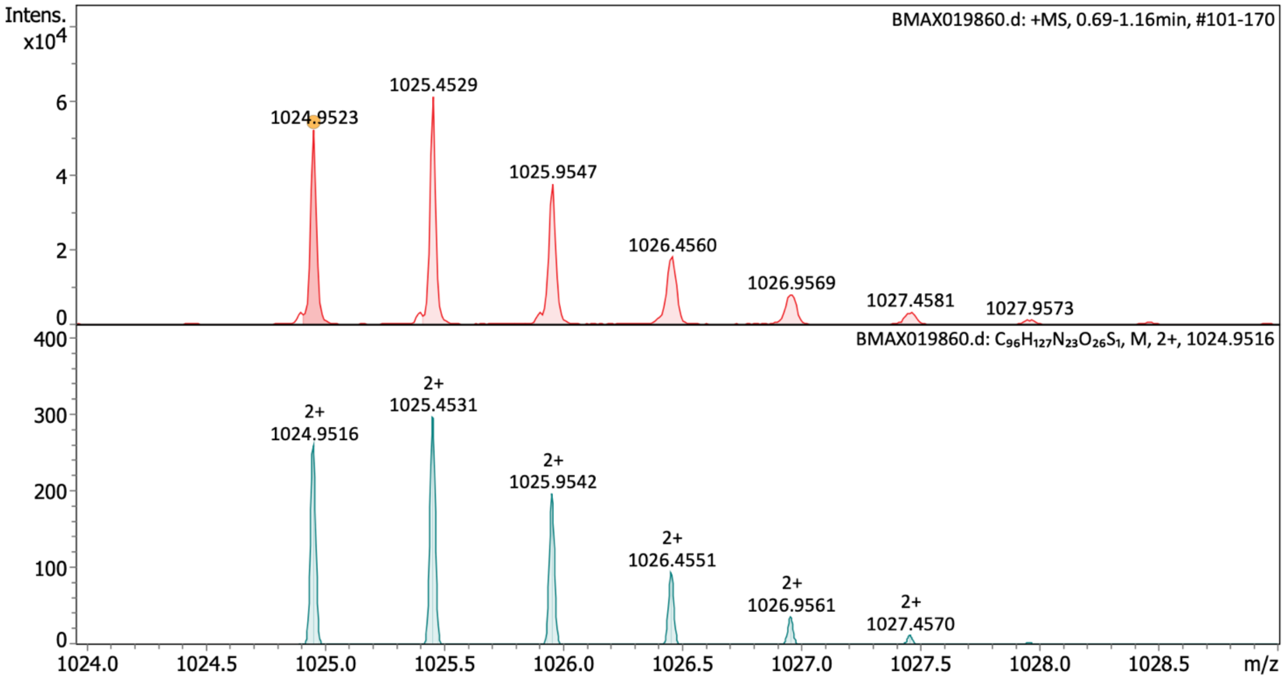

**Characterization of fluorescein-pep3–R–$CONH_2$** HRMS (ESI) spectrum of purified peptide showing recorded mass spectrum (upper panel) and calculated spectrum (lower panel).

Synthesis of fluorescein-pep3–S–$CONH_2$

**Fluorescein-KKYRYDVPDYSAS–$CONH_2$**

The peptide was obtained as an orange solid and used without preparative RP-HPLC purification. HRMS (ESI): calculated for $[C_{93}H_{120}N_{20}O_{27}S_1]^{2+}$: m/z 990.4171, found: m/z 990.4171

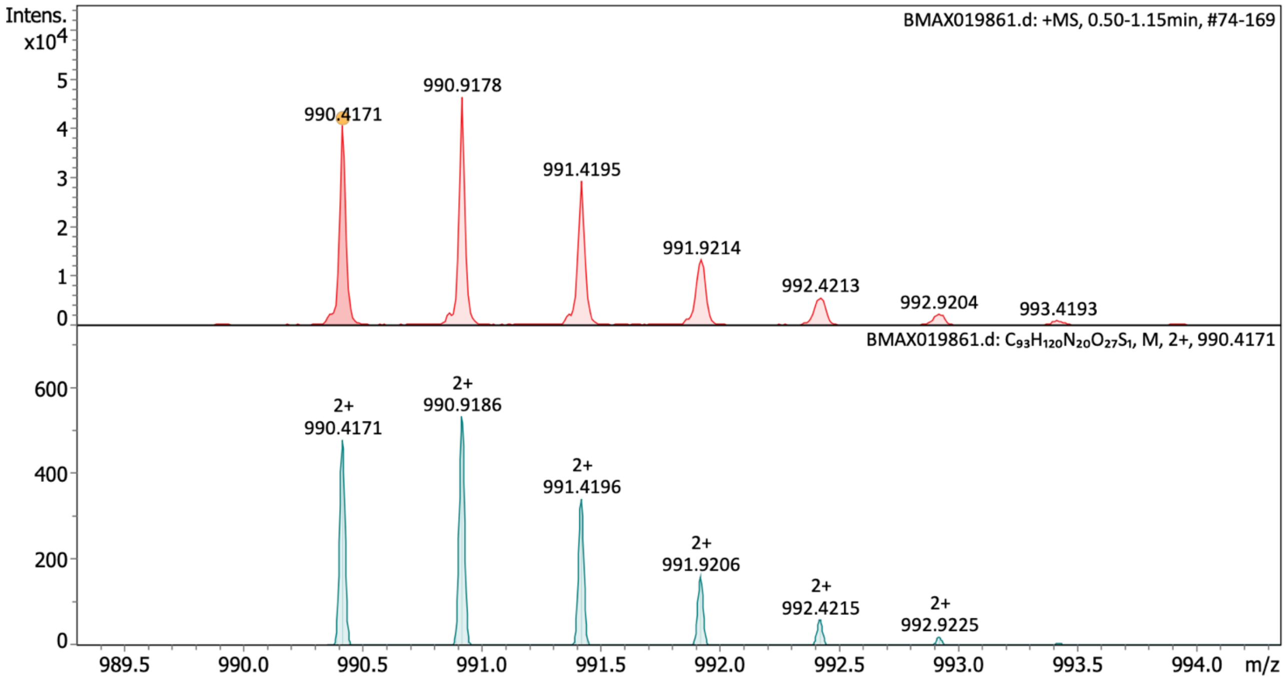

**Characterization of fluorescein-pep3–S–$CONH_2$** HRMS (ESI) spectrum of purified peptide showing recorded mass spectrum (upper panel) and calculated spectrum (lower panel).

Synthesis of fluorescein-pep3–T–$CONH_2$

**Fluorescein-KKYRYDVPDYSAT–$CONH_2$**

The peptide was obtained as an orange solid and used without preparative RP-HPLC purification. HRMS (ESI): calculated for $[C_{94}H_{122}N_{20}O_{27}S_1]^{2+}$: m/z 997.4249, found: m/z 997.4243

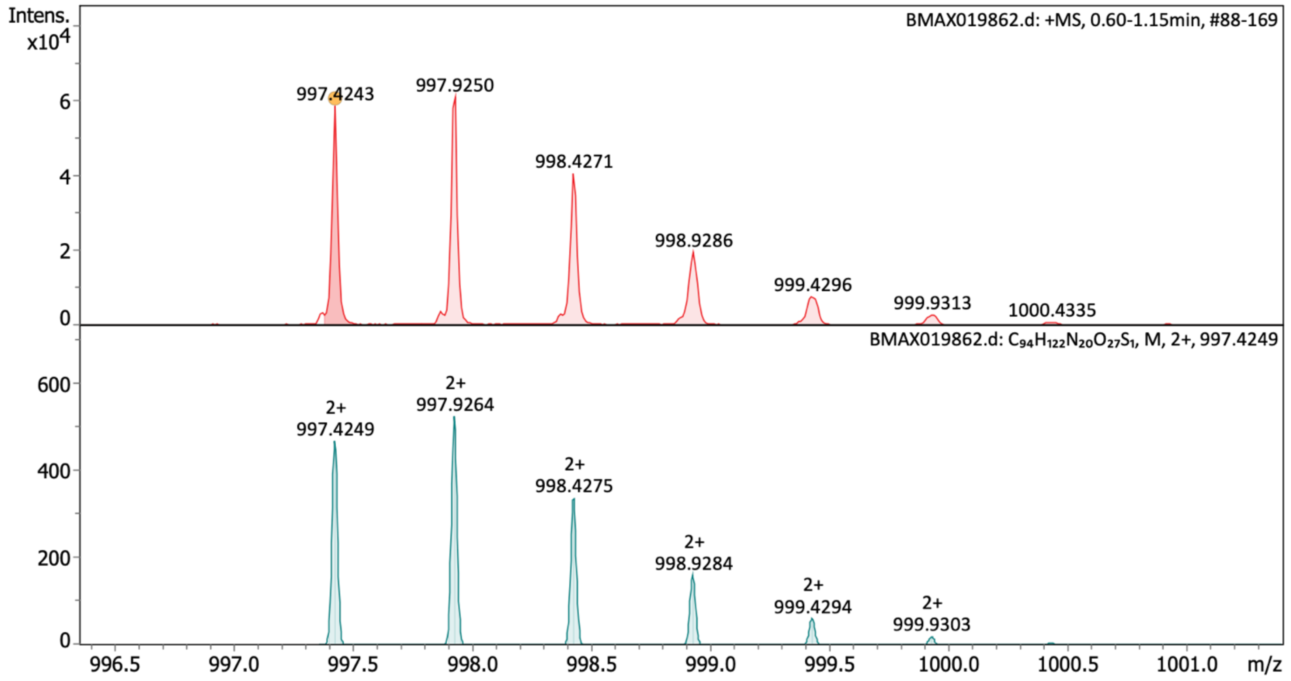

**Characterization of fluorescein-pep3–T–$CONH_2$** HRMS (ESI) spectrum of purified peptide showing recorded mass spectrum (upper panel) and calculated spectrum (lower panel).

Synthesis of fluorescein-pep3–V–$CONH_2$

**Fluorescein-KKYRYDVPDYSAV–$CONH_2$**

The peptide was obtained as an orange solid and used without preparative RP-HPLC purification. HRMS (ESI): calculated for $[C_{95}H_{124}N_{20}O_{26}S_1]^{2+}$: m/z 996.4353, found: m/z 996.4367

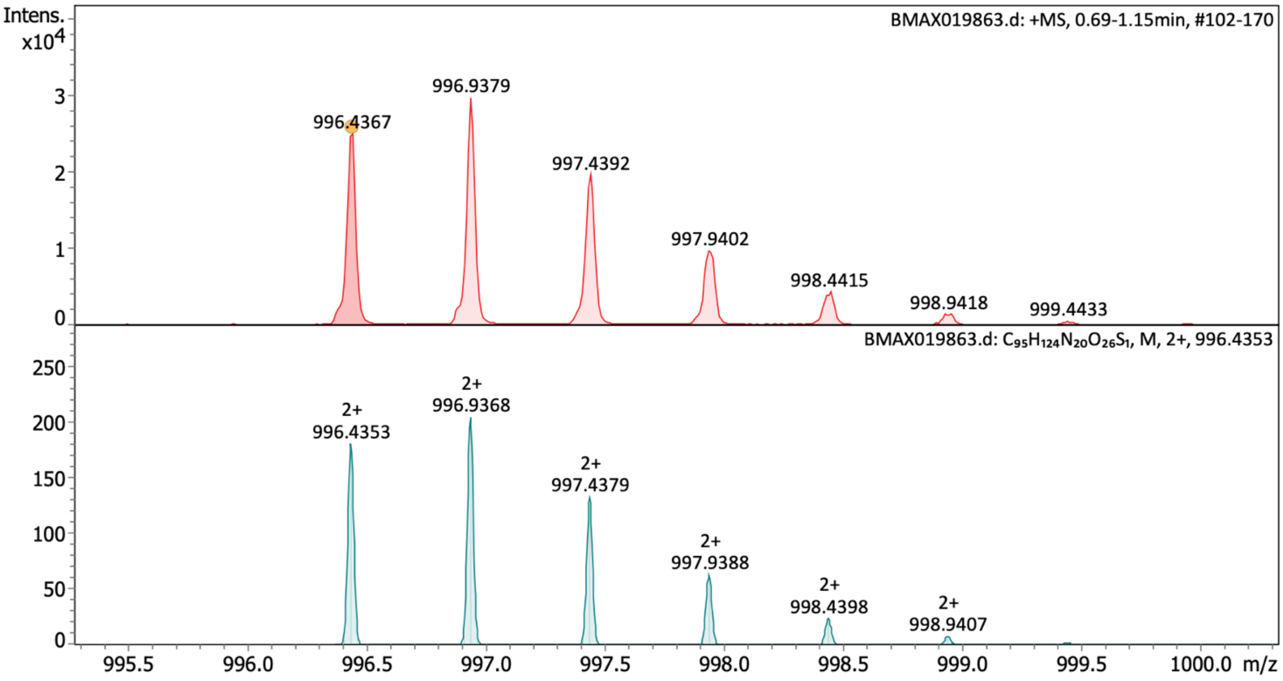

**Characterization of fluorescein-pep3–V–$CONH_2$** HRMS (ESI) spectrum of purified peptide showing recorded mass spectrum (upper panel) and calculated spectrum (lower panel).

Synthesis of fluorescein-pep3–W–$CONH_2$

**Fluorescein-KKYRYDVPDYSAW–$CONH_2$**

The peptide was obtained as an orange solid and used without preparative RP-HPLC purification. HRMS (ESI): calculated for $[C_{101}H_{125}N_{21}O_{26}S_1]^{2+}$: m/z 1039.9407, found: m/z 1039.9385

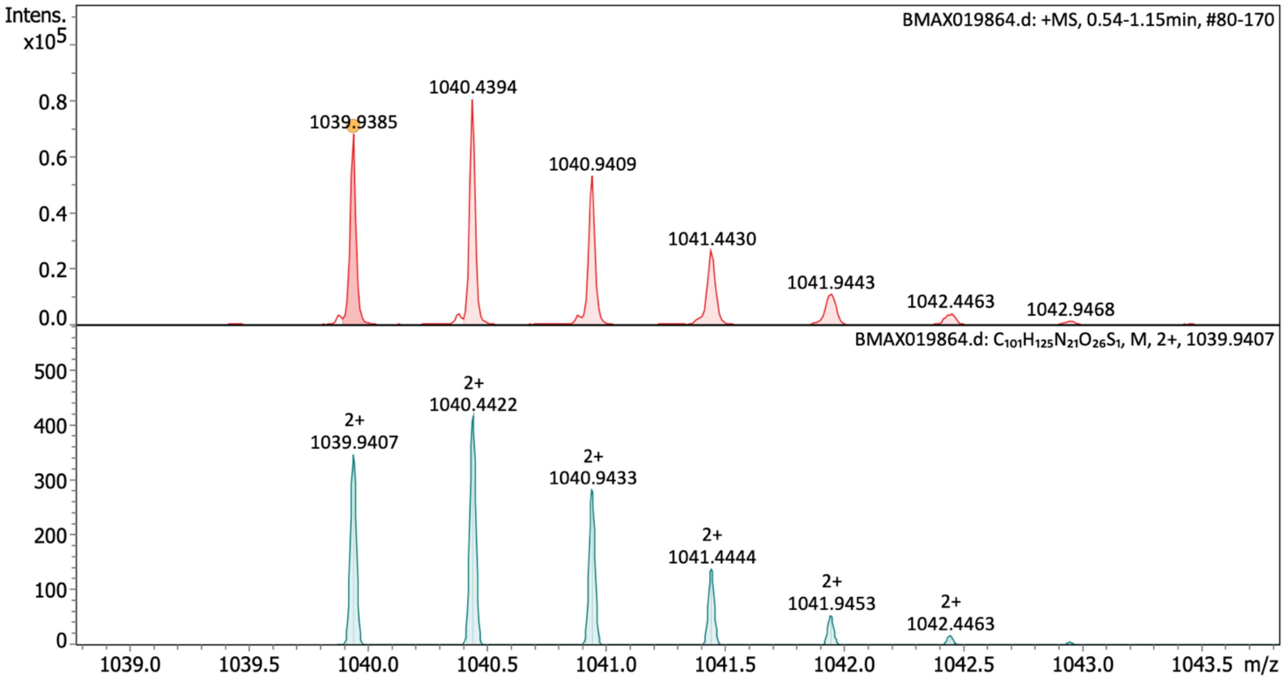

**Characterization of fluorescein-pep3–W–$CONH_2$** HRMS (ESI) spectrum of purified peptide showing recorded mass spectrum (upper panel) and calculated spectrum (lower panel).

Synthesis of fluorescein-pep3–Y–$CONH_2$

**Fluorescein-KKYRYDVPDYSAY–$CONH_2$**

The peptide was obtained as an orange solid and used without preparative RP-HPLC purification. HRMS (ESI): calculated for $[C_{99}H_{124}N_{20}O_{27}S_1]^{2+}$: m/z 1028.4327, found: m/z 1028.4341

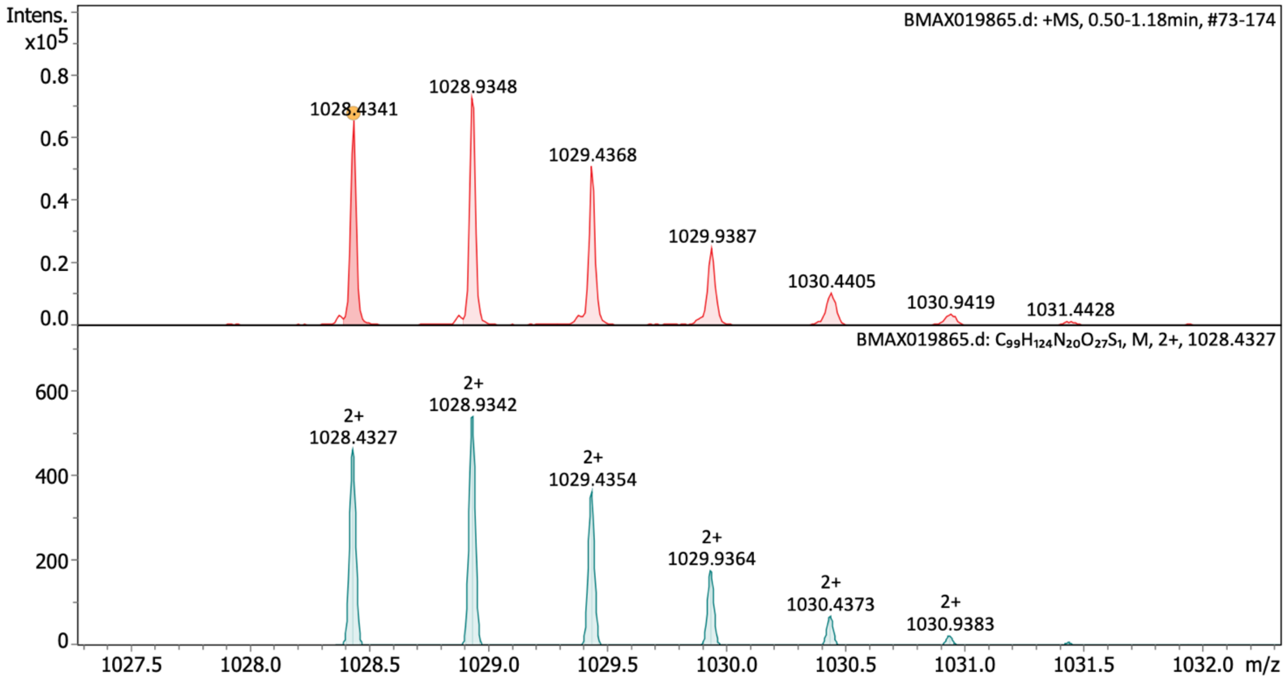

**Characterization of fluorescein-pep3–Y–$CONH_2$** HRMS (ESI) spectrum of purified peptide showing recorded mass spectrum (upper panel) and calculated spectrum (lower panel).

Synthesis of fluorescein-pep3–N–COOH

**Fluorescein-KKYRYDVPDYSAN–COOH**

The peptide was obtained as an orange solid.
HRMS (ESI): calculated for $[C_{94}H_{120}N_{20}O_{28}S_1]^{2+}$: m/z 1004.4145, found: m/z 1004.4155

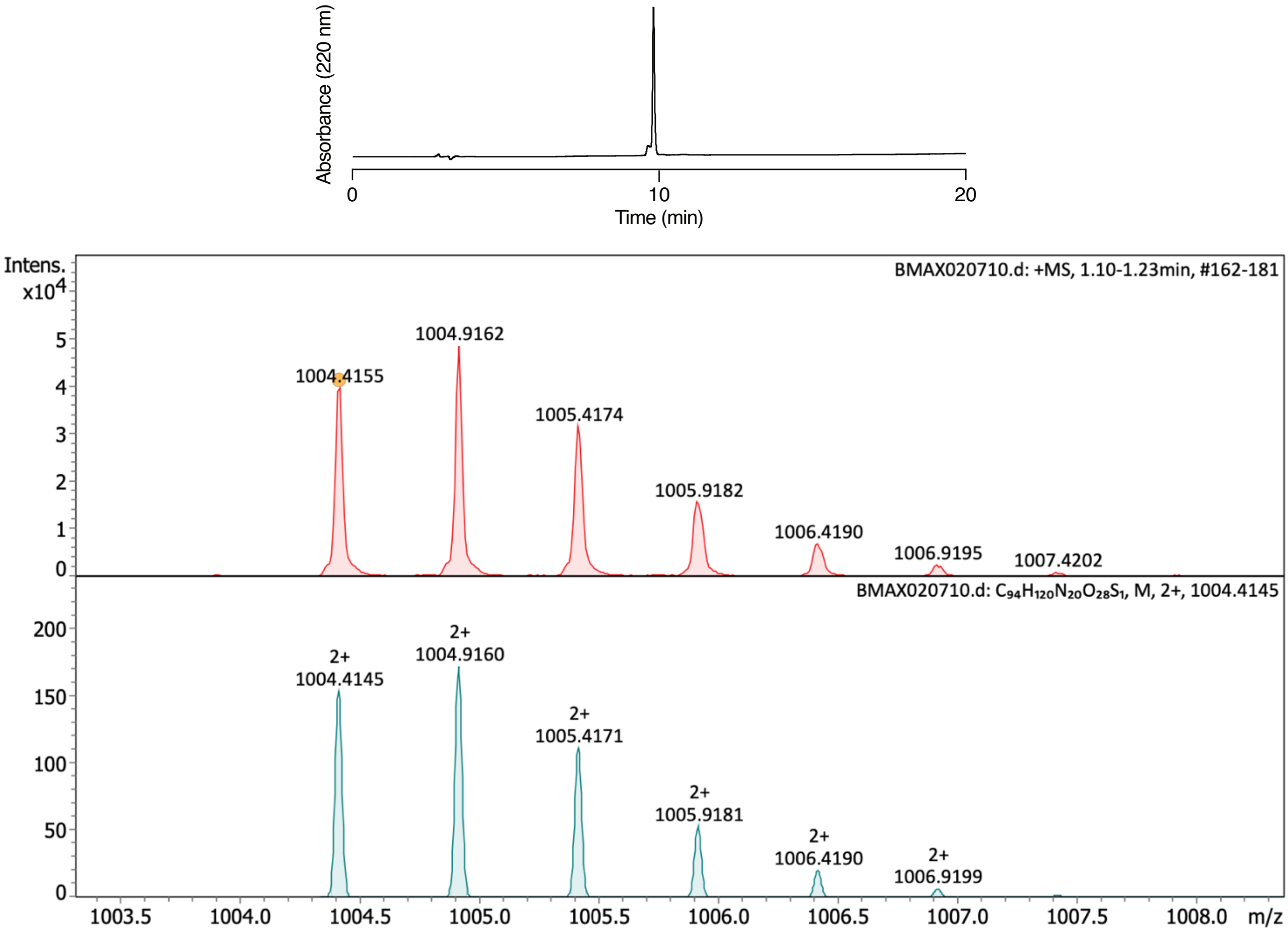

**Characterization of fluorescein-pep3–N–COOH** Analytical RP-HPLC of purified peptide. HRMS (ESI) spectrum of purified peptide showing recorded mass spectrum (upper panel) and calculated spectrum (lower panel).

Synthesis of fluorescein-pep3–Q–COOH

**Fluorescein-KKYRYDVPDYSAQ–COOH**

The peptide was obtained as an orange solid.
HRMS (ESI): calculated for $[C_{95}H_{122}N_{20}O_{28}S_1]^{2+}$: m/z 1011.4224, found: m/z 1011.4222

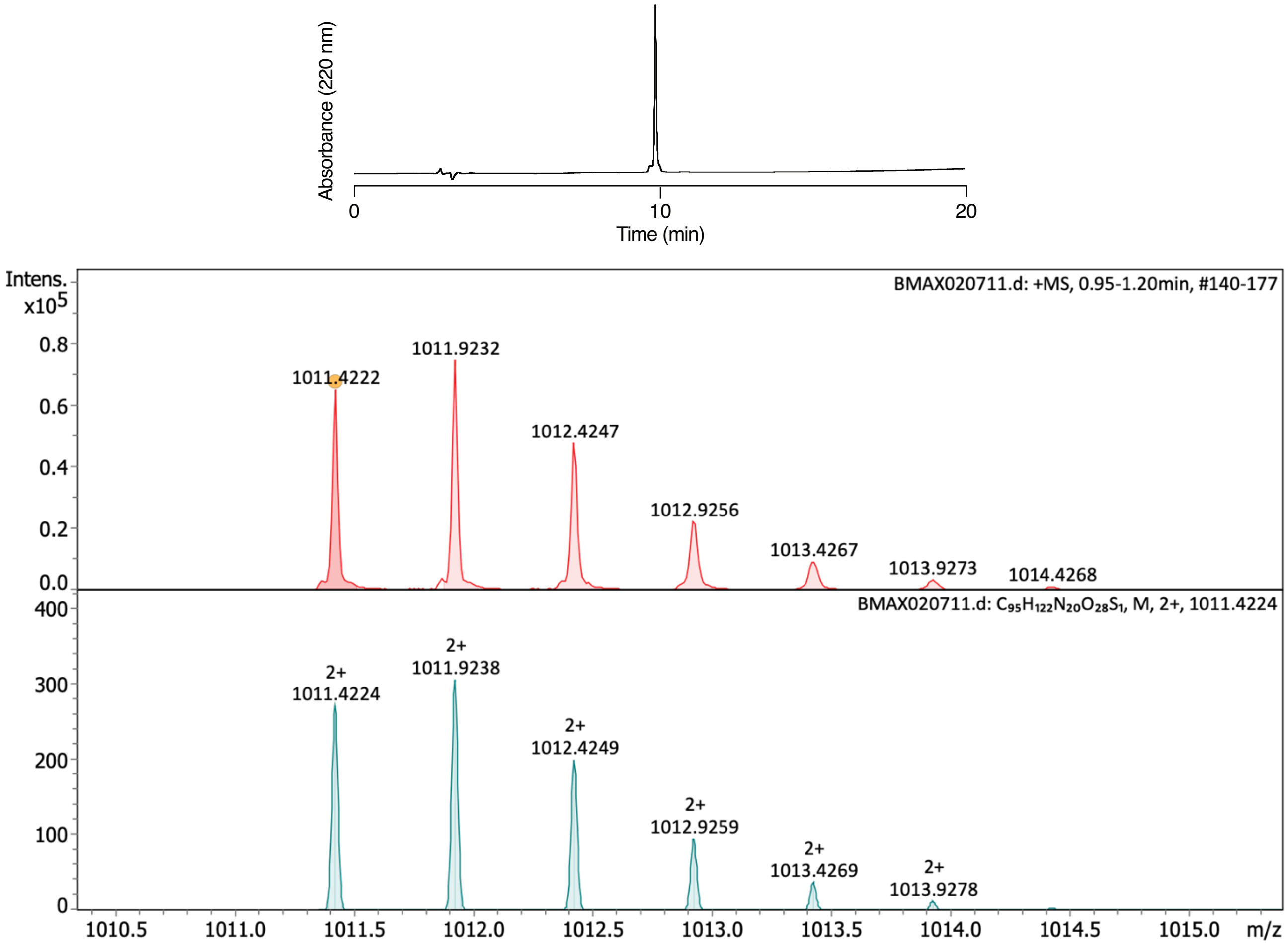

**Characterization of fluorescein-pep3–Q–COOH** Analytical RP-HPLC of purified peptide. HRMS (ESI) spectrum of purified peptide showing recorded mass spectrum (upper panel) and calculated spectrum (lower panel).

Synthesis of fluorescein-pep3–R–$CH_2OH$

**Fluorescein- KEEDEKGSRASDDFRDLR–$CH_2OH$**

The peptide was obtained as an orange solid.
HRMS (ESI): calculated for $[C_{108}H_{157}N_{30}O_{39}S_1]^{3+}$: m/z 843.3643, found: m/z 843.3649

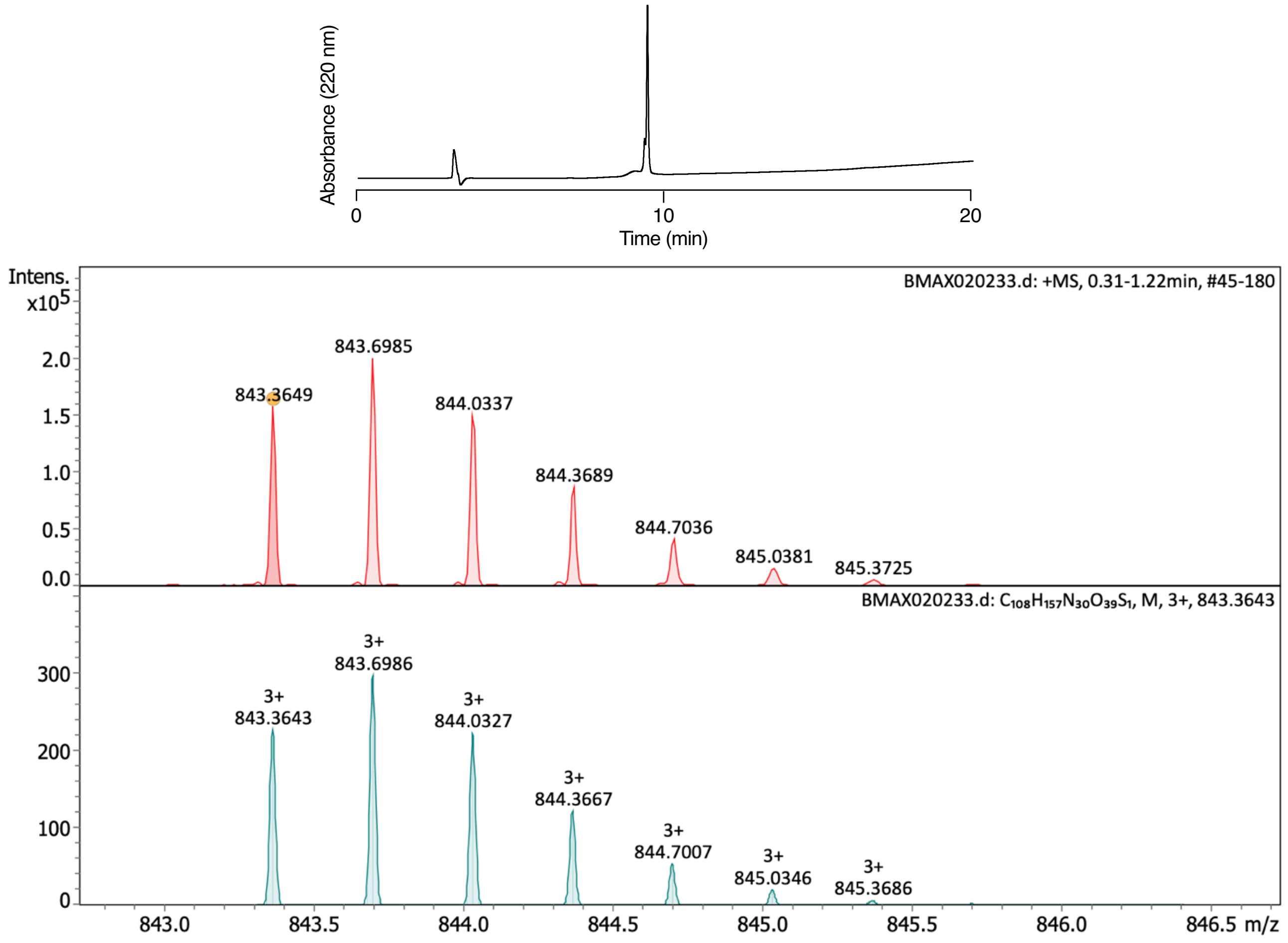

**Characterization of fluorescein-pep3–R–$CH_2OH$** Analytical RP-HPLC of purified peptide. HRMS (ESI) spectrum of purified peptide showing recorded mass spectrum (upper panel) and calculated spectrum (lower panel).

Synthesis of fluorescein-pep3–R–COOMe

**Fluorescein- KEEDEKGSRASDDFRDLR–COOMe**

The peptide was obtained as an orange solid.
HRMS (ESI): calculated for $[C_{109}H_{155}N_{30}O_{40}S_1]^{3+}$: m/z 852.0240, found: m/z 852.0253

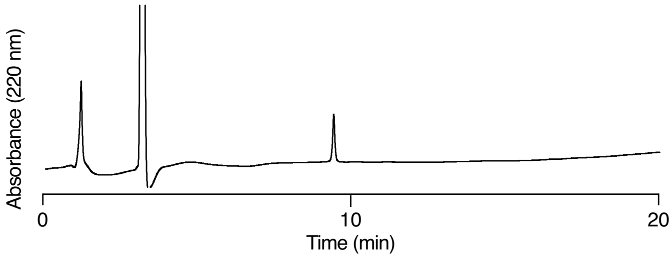

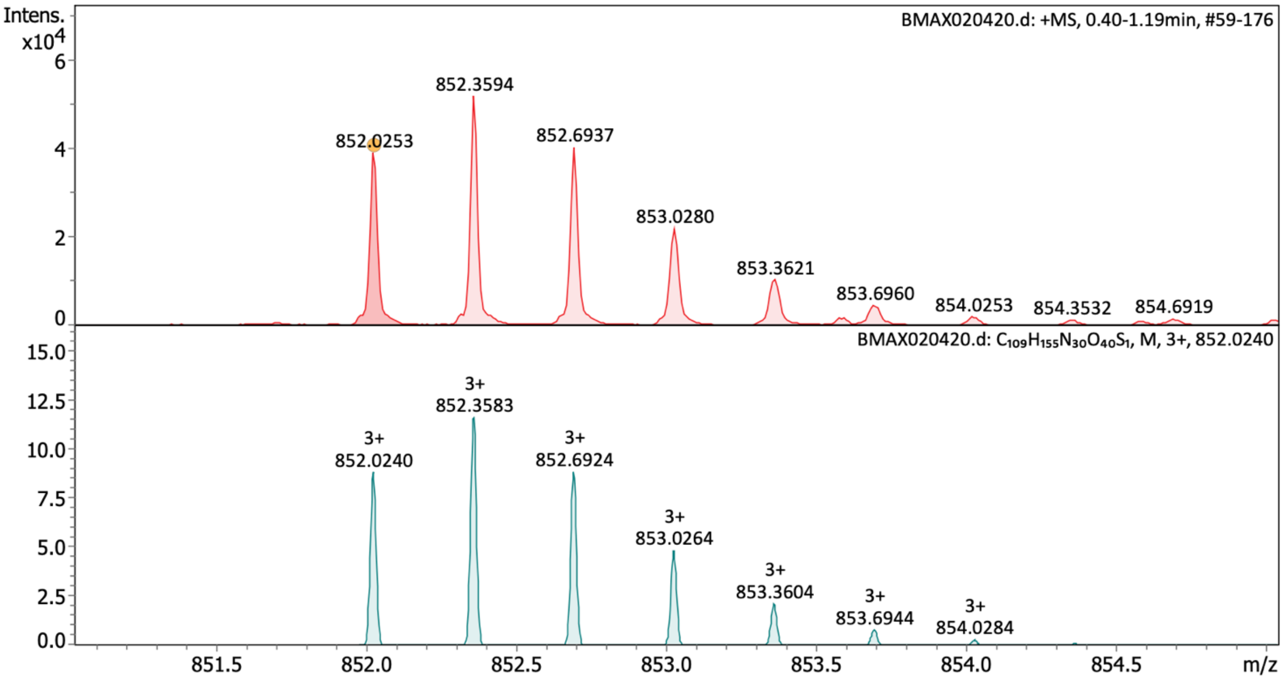

**Characterization of fluorescein-pep3–R–COOMe** Analytical RP-HPLC of purified peptide. HRMS (ESI) spectrum of purified peptide showing recorded mass spectrum (upper panel) and calculated spectrum (lower panel).

#### Characterization of SPPS products for sortase reaction

Synthesis of pep1 [homocitrulline (HCT)]–D–COOH

**GGGKDLEGKGGS[HCT]GSGS[HCT]GGSKYPYDVPDYAKD–COOH**

The peptide was obtained as a white solid.
HRMS (ESI): calculated for $[C_{142}H_{219}N_{41}O_{53}]^{+}$: 3346.5702, found: m/z 3346.5837

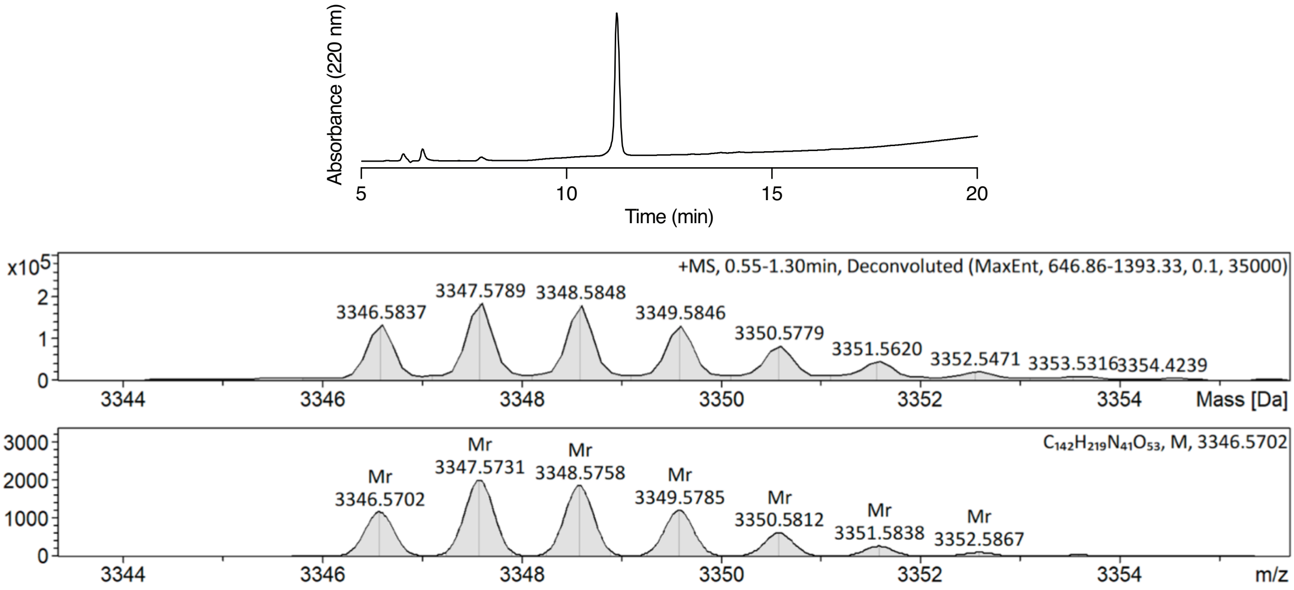

**Characterization of pep1 [HCT]–D–COOH** Analytical RP-HPLC of purified peptide. HRMS (ESI) spectrum of purified peptide showing recorded mass spectrum (upper panel) and calculated spectrum (lower panel).

Synthesis of pep1 [L-DOP (LDO)]–D–COOH

**GGGKDLEGKGGS[LDO]GSGS[LDO]GGSKYPYDVPDYAKD–COOH**

The peptide was obtained as a white solid.
HRMS (ESI): calculated for $[C_{146}H_{211}N_{37}O_{55}]^{+}$: 3362.4851, found: m/z 3362.4961

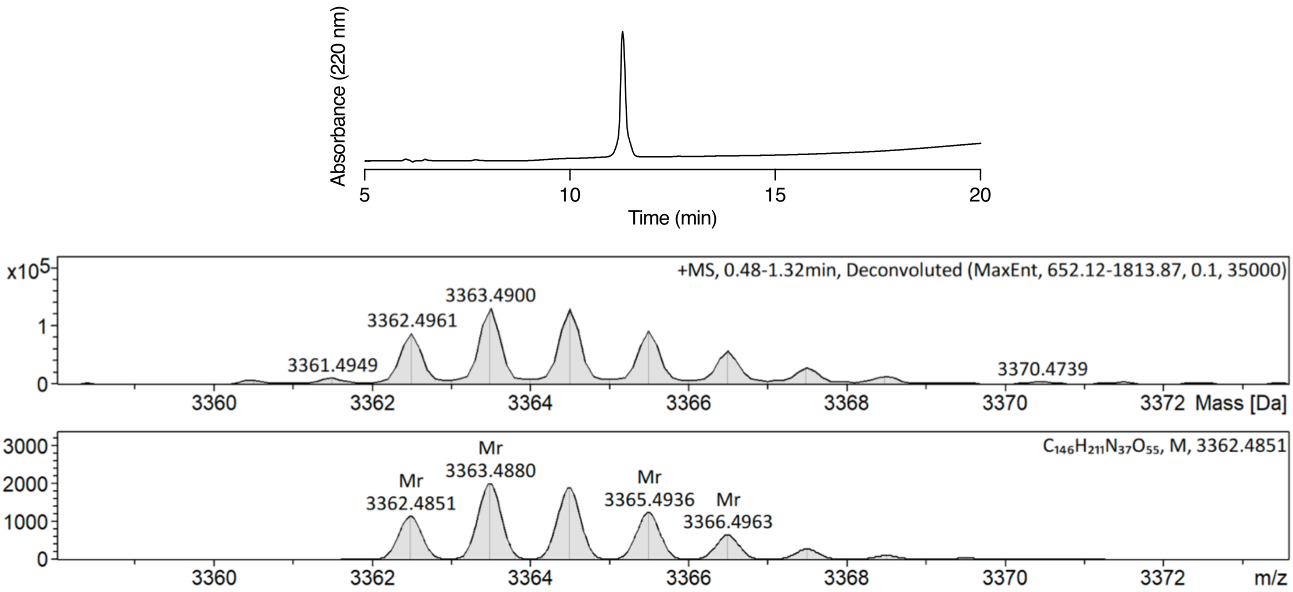

**Characterization of pep1 [LDO]–D–COOH** Analytical RP-HPLC of purified peptide. HRMS (ESI) spectrum of purified peptide showing recorded mass spectrum (upper panel) and calculated spectrum (lower panel).

Synthesis of pep1 [carboxymethyllysine (CML)]–D–COOH

**GGGKDLEGKGGS[CML]GSGS[CML]GGSKYPYDVPDYAKD–COOH**

The peptide was obtained as a white solid.
HRMS (ESI): calculated for $[C_{144}H_{221}N_{39}O_{55}]^{+}$: m/z 3376.5695, found: m/z 3376.5792

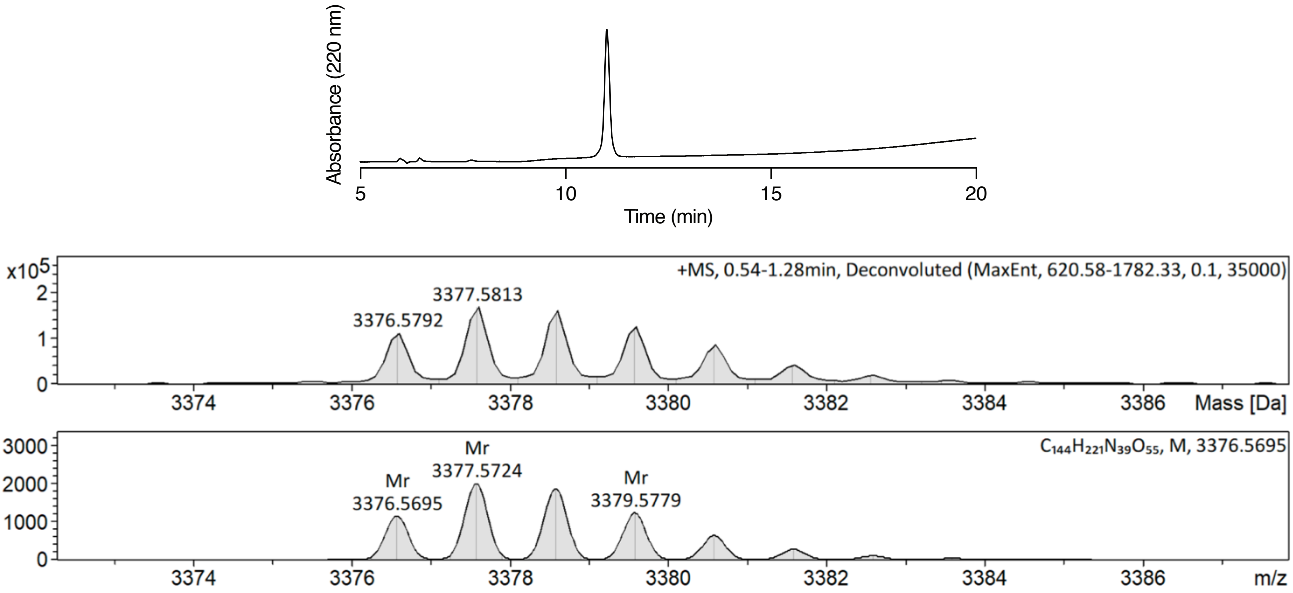

**Characterization of pep1 [CML]–D–COOH** Analytical RP-HPLC of purified peptide. HRMS (ESI) spectrum of purified peptide showing recorded mass spectrum (upper panel) and calculated spectrum (lower panel).

Synthesis of pep1 [A]–S–COOH

**GGGKDLEGKGGSAGSGSAGGSKYPYDVPDYAKS-COOH**

The peptide was obtained as a white solid.
HRMS (ESI): calculated for $[C_{133}H_{203}N_{37}O_{50}]^{+}$: m/z 3118.4480 found: m/z 3118.4579

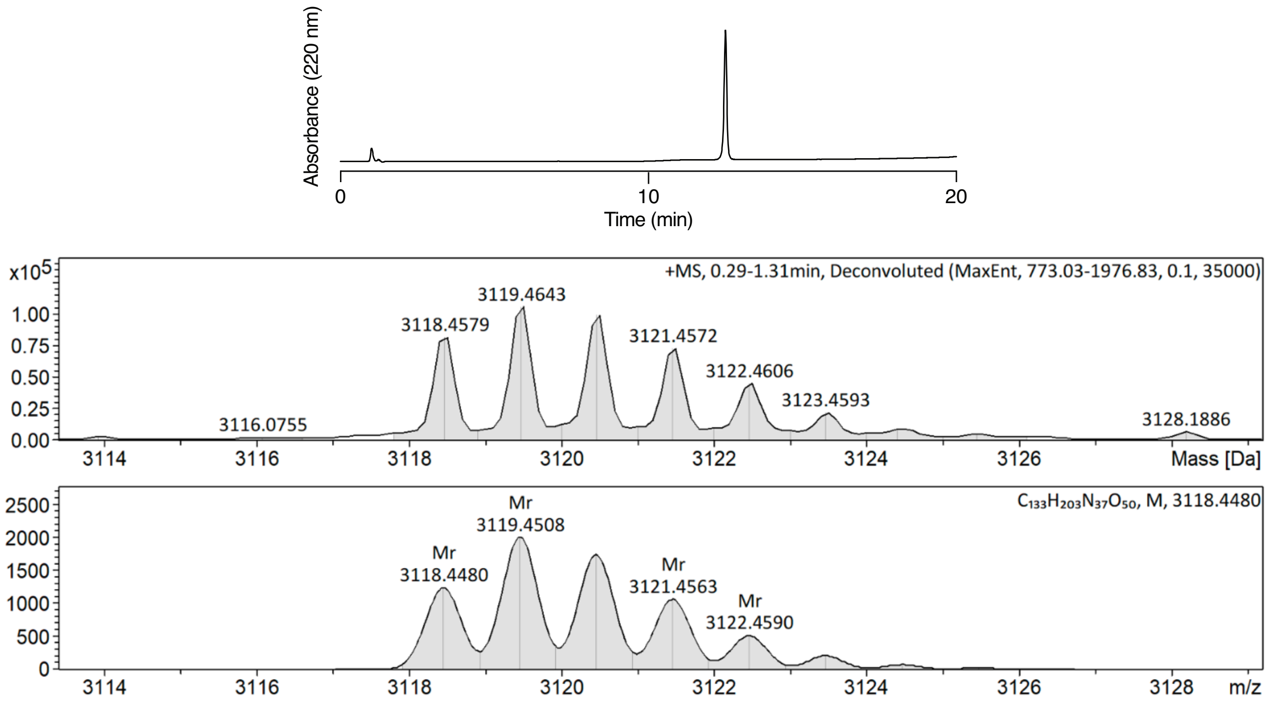

**Characterization of pep1–S–COOH** Analytical RP-HPLC of purified peptide. HRMS (ESI) spectrum of purified peptide showing recorded mass spectrum (upper panel) and calculated spectrum (lower panel).

Synthesis of pep1 [A]–S–$CONH_2$

**GGGKDLEGKGGSAGSGSAGGSKYPYDVPDYAKS-$CONH_2$**

The peptide was obtained as a white solid.
HRMS (ESI): calculated for $[C_{133}H_{204}N_{38}O_{49}]^+$: m/z 3117.4639 found: m/z 3117.4830

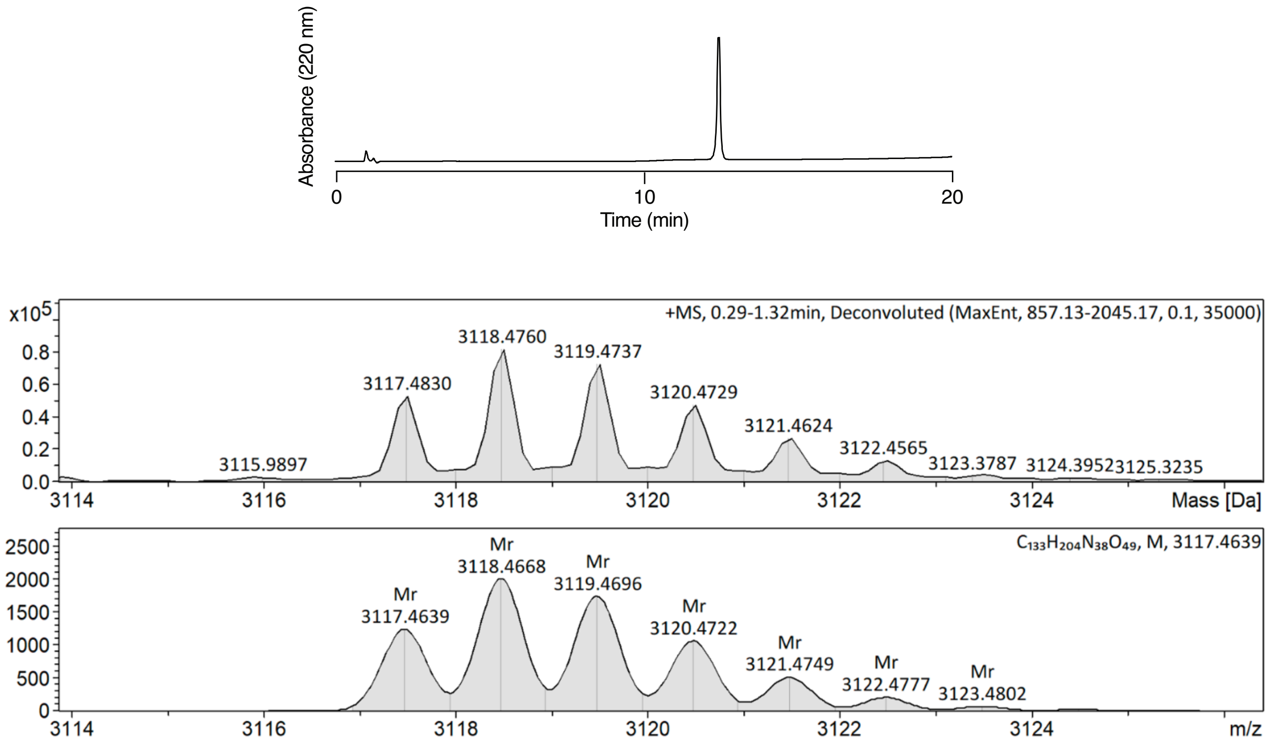

**Characterization of pep1–S–$CONH_2$** Analytical RP-HPLC of purified peptide. HRMS (ESI) spectrum of purified peptide showing recorded mass spectrum (upper panel) and calculated spectrum (lower panel).

Synthesis of pep1 [A]–D–COOH

**GGGKDLEGKGGSAGSGSAGGSKYPYDVPDYAKD–COOH**

The peptide was obtained as a white solid.
HRMS (ESI): calculated for $[C_{134}H_{203}N_{37}O_{51}]^{+}$: m/z 3146.4429, found: m/z 3146.4586

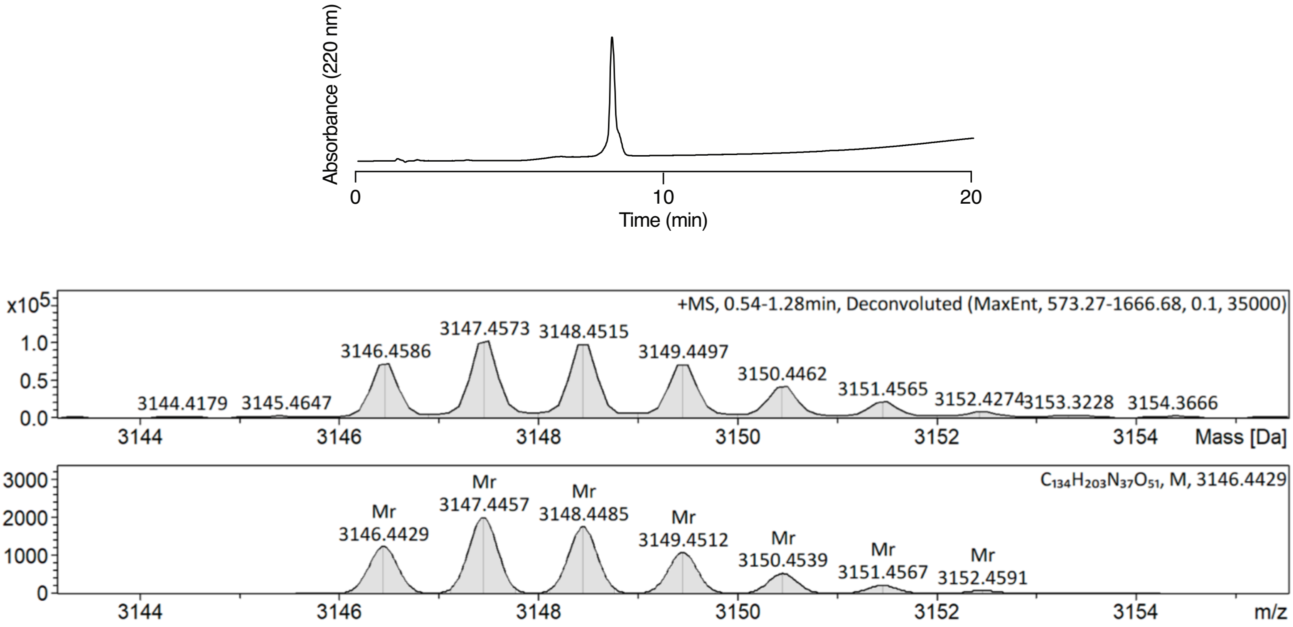

**Characterization of pep1 [A]–D–COOH** Analytical RP-HPLC of purified peptide. HRMS (ESI) spectrum of purified peptide showing recorded mass spectrum (upper panel) and calculated spectrum (lower panel).

<u>Synthesis of pep1 [hexanoyllysine (KHL)]–D–COOH</u>

**GGGKDLEGKGGS[KHL]GSGS[KHL]GGSKYPYDVPDYAKD–COOH**

The peptide was obtained as a white solid.
HRMS (ESI): calculated for $[C_{152}H_{237}N_{39}O_{53}]^{+}$: m/z 3456.7049, found: m/z 3456.7200

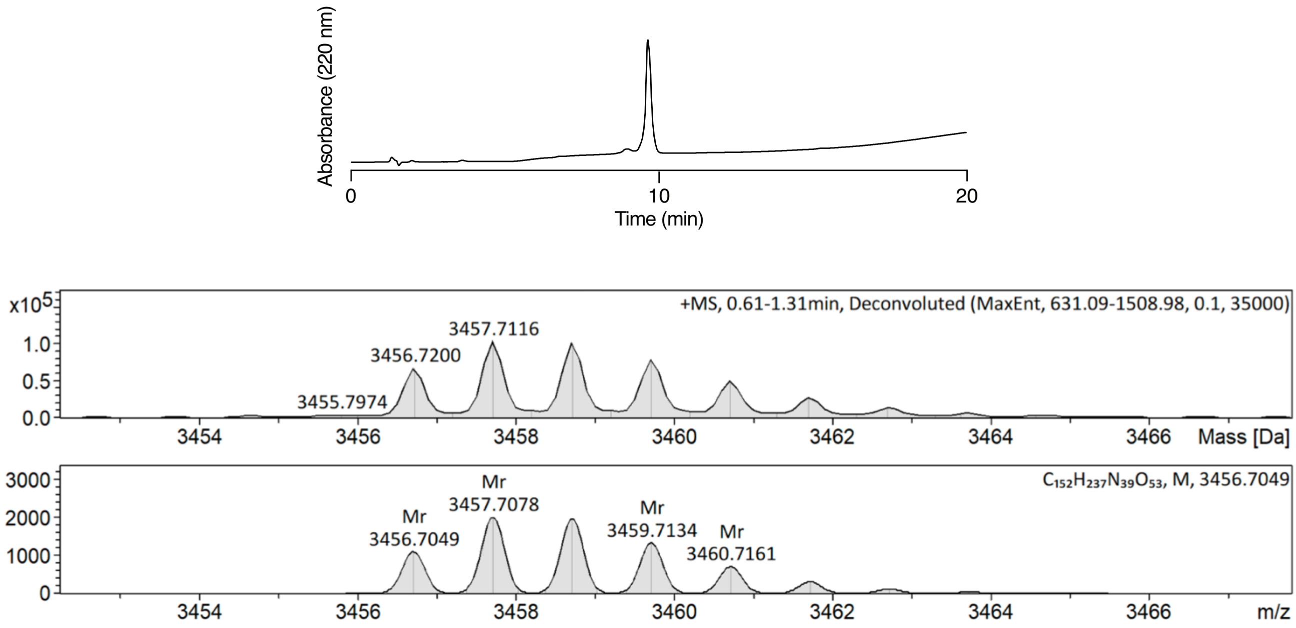

**Characterization of pep1 [KHL]–D–COOH** Analytical RP-HPLC of purified peptide. HRMS (ESI) spectrum of purified peptide showing recorded mass spectrum (upper panel) and calculated spectrum (lower panel).

Synthesis of pep2 [DxxD]–R–COOH

**GGGRRLEGKEEDEKGSRASDDFRDLR–COOH**

The peptide was obtained as a white solid.
HRMS (ESI): calculated for $[C_{118}H_{197}N_{43}O_{45}]^{2+}$: m/z 1468.2219, found: m/z 1468.2226

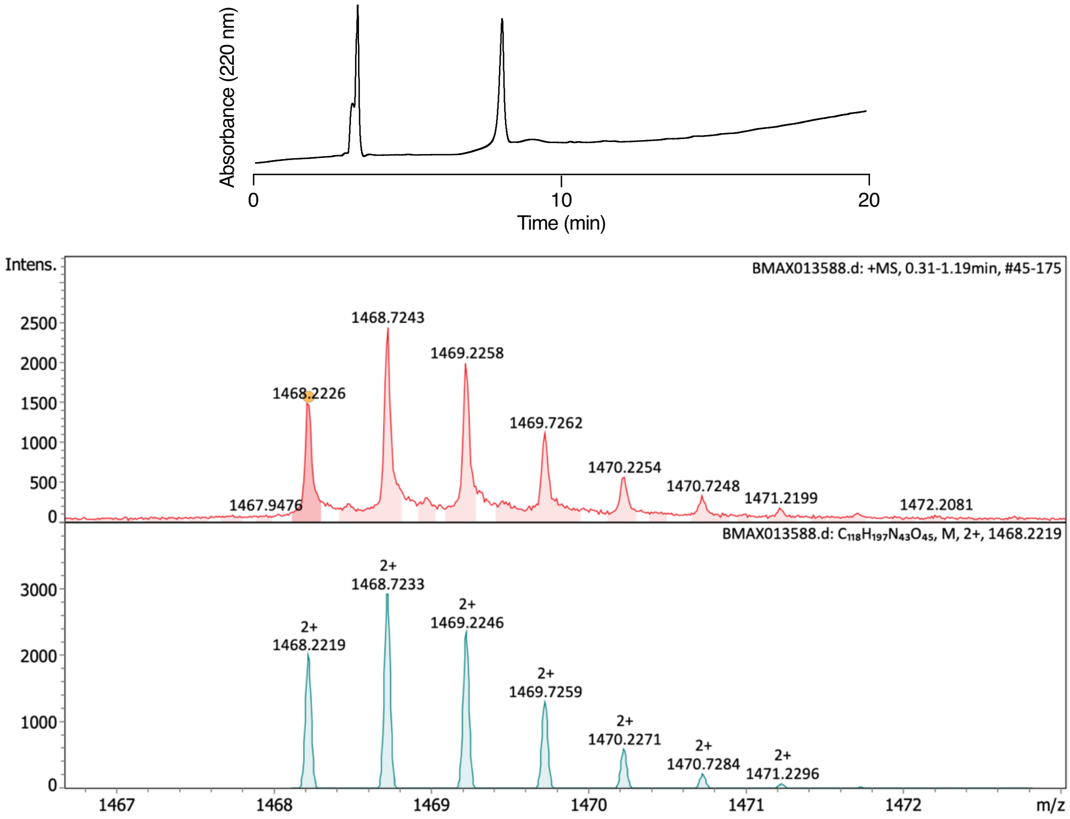

**Characterization of pep2 [DxxD]–R–COOH** Analytical RP-HPLC of purified peptide. HRMS (ESI) spectrum of purified peptide showing recorded mass spectrum (upper panel) and calculated spectrum (lower panel).

Synthesis of pep2 [DxxD]–R–$CONH_2$

**GGGRRLEGKEEDEKGSRASDDFRDLR–$CONH_2$**

The peptide was obtained as a white solid.
HRMS (ESI): calculated for $[C_{118}H_{199}N_{44}O_{44}]^{2+}$: m/z 1467.7299, found: m/z 1467.7301

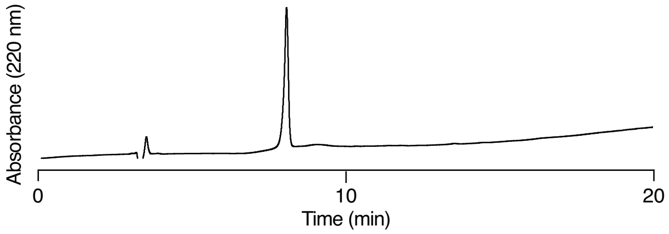

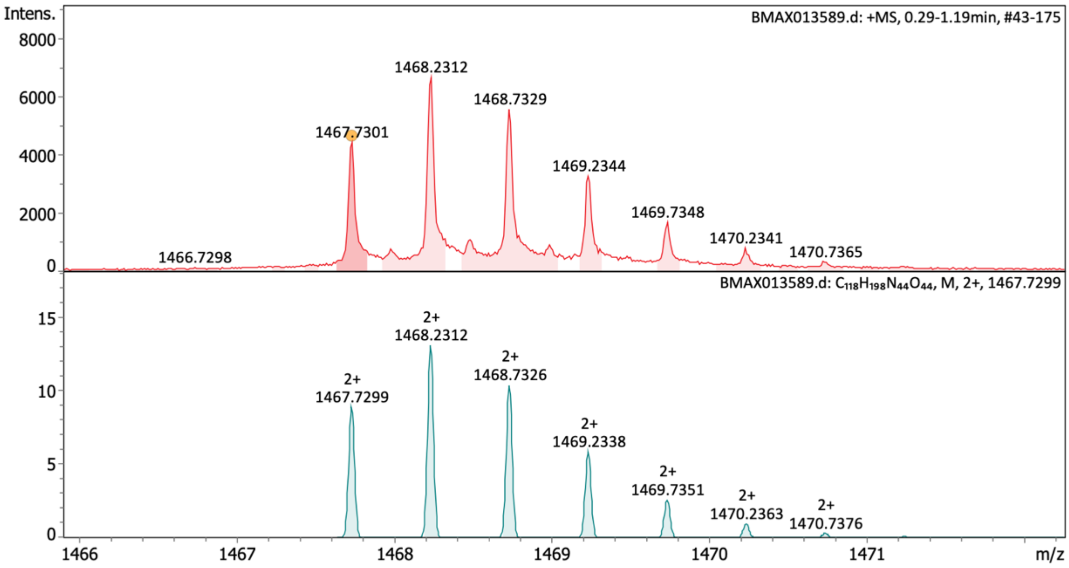

**Characterization of pep2 [DxxD]–R–CONH2** Analytical RP-HPLC of purified peptide. HRMS (ESI) spectrum of purified peptide showing recorded mass spectrum (upper panel) and calculated spectrum (lower panel).

Synthesis of pep2 [RxxG]–R–COOH

**GGGRRLEGKEEDEKGSRASDRFRGLR–COOH**

The peptide was obtained as a white solid.
HRMS (ESI): calculated for $[C_{118}H_{200}N_{46}O_{41}]^{+}$: m/z 2917.4979, found: m/z 2917.5090

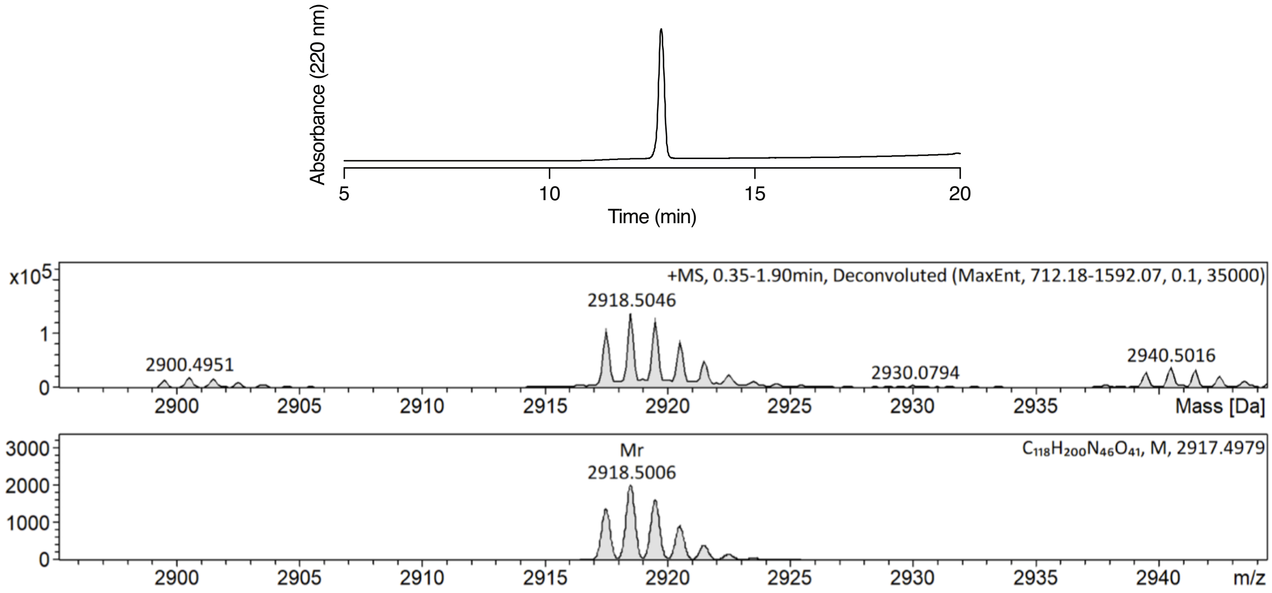

**Characterization of pep2 [RxxG]–R–COOH** Analytical RP-HPLC of purified peptide. HRMS (ESI) spectrum of purified peptide showing recorded mass spectrum (upper panel) and calculated spectrum (lower panel).

Synthesis of pep2 [RxxG]–R–CONH2

**GGGRRLEGKEEDEKGSRASDRFRGLR–CONH2**

The peptide was obtained as a white solid.
HRMS (ESI): calculated for $[C_{118}H_{201}N_{47}O_{40}]^{+}$: m/z 2916.5139, found: m/z 2916.5309

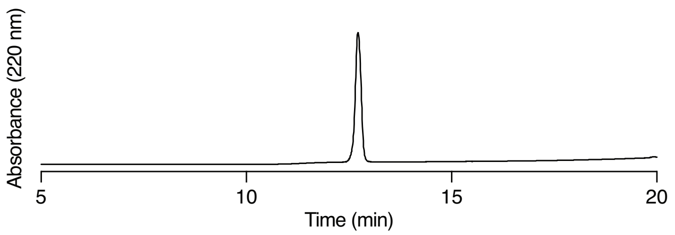

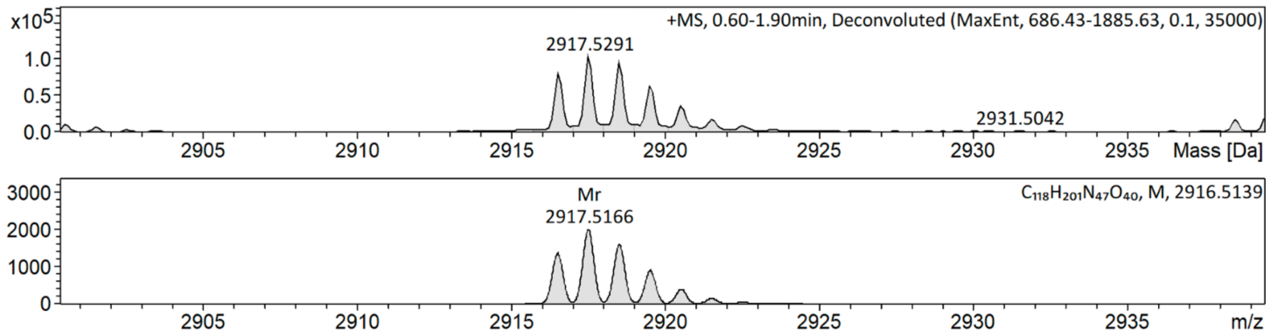

**Characterization of pep2 [RxxG]–R–CONH2** Analytical RP-HPLC of purified peptide. HRMS (ESI) spectrum of purified peptide showing recorded mass spectrum (upper panel) and calculated spectrum (lower panel).

#### Characterization of protein-peptide conjugates

Preparation of GFP–GGGKDLEGKGGS[HCT]GSGS[HCT]GGSKYPYDVPDYAKD–COOH

Protein was prepared as described in the general method.

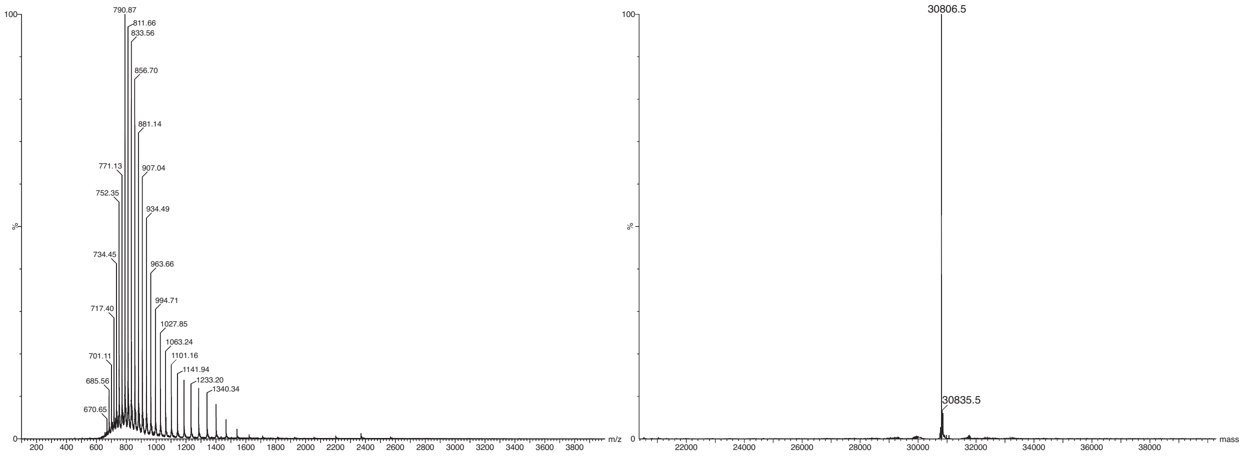

**ESI-MS characterization of protein conjugate sfGFP–pep1 [HCT]** Complex mass spectrum (left panel). Deconvoluted mass spectrum (right panel). Found: 30806.5, calculated for [$C_{1366}H_{2113}N_{371}O_{428}S_{7}$]: 30805.0.

Preparation of GFP– GGGKDLEGKGGS[LDO]GSGS[LDO]GGSKYPYDVPDYAKD–COOH

Protein was prepared as described in the general method.

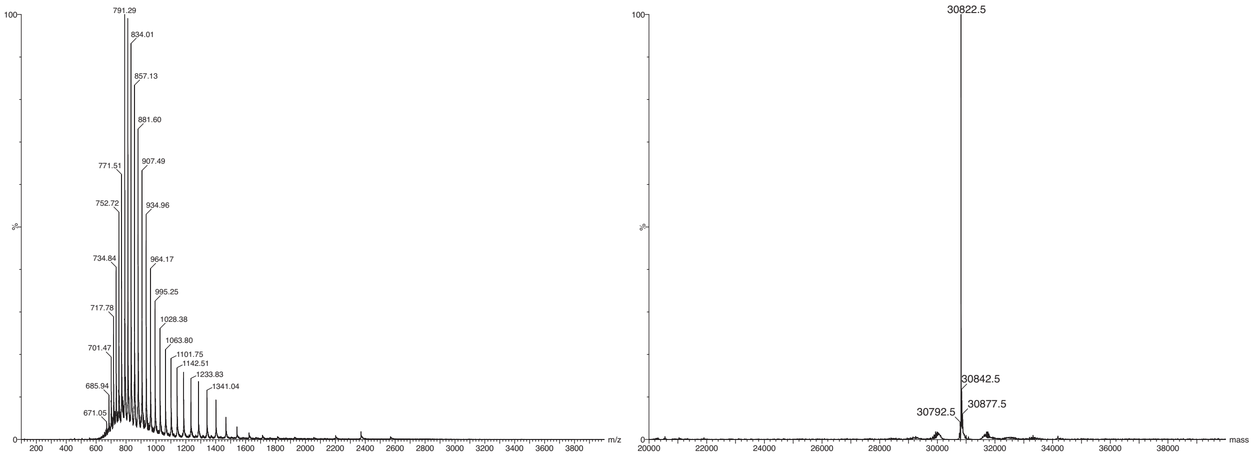

**ESI-MS characterization of protein conjugate sfGFP–pep1 [LDO]** Complex mass spectrum (left panel). Deconvoluted mass spectrum (right panel). Found: 30822.5, calculated for [$C_{1370}H_{2105}N_{369}O_{430}S_{7}$]: 30820.9.

Preparation of GFP– GGGKDLEGKGGS[CML]GSGS[CML]GGSKYPYDVPDYAKD–COOH

Protein was prepared as described in the general method.

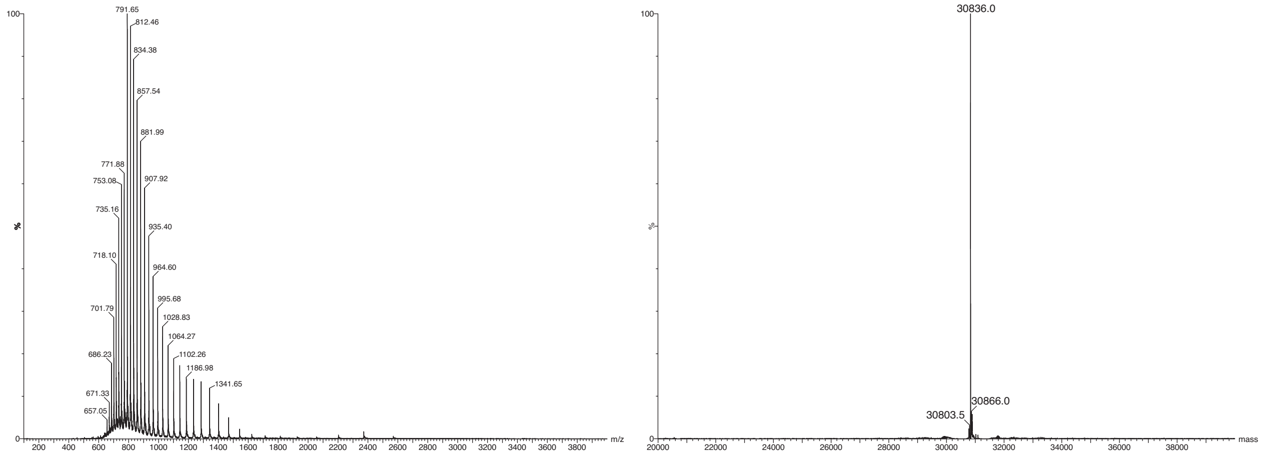

**ESI-MS characterization of protein conjugate sfGFP–pep1 [CML]** Complex mass spectrum (left panel). Deconvoluted mass spectrum (right panel). Found: 30836.0, calculated for [$C_{1368}H_{2115}N_{369}O_{430}S_7$]: 30835.0.

Preparation of GFP– GGGKDLEGKGGSAGSGSAGGSKYPYDVPDYAKD–COOH

Protein was prepared as described in the general method.

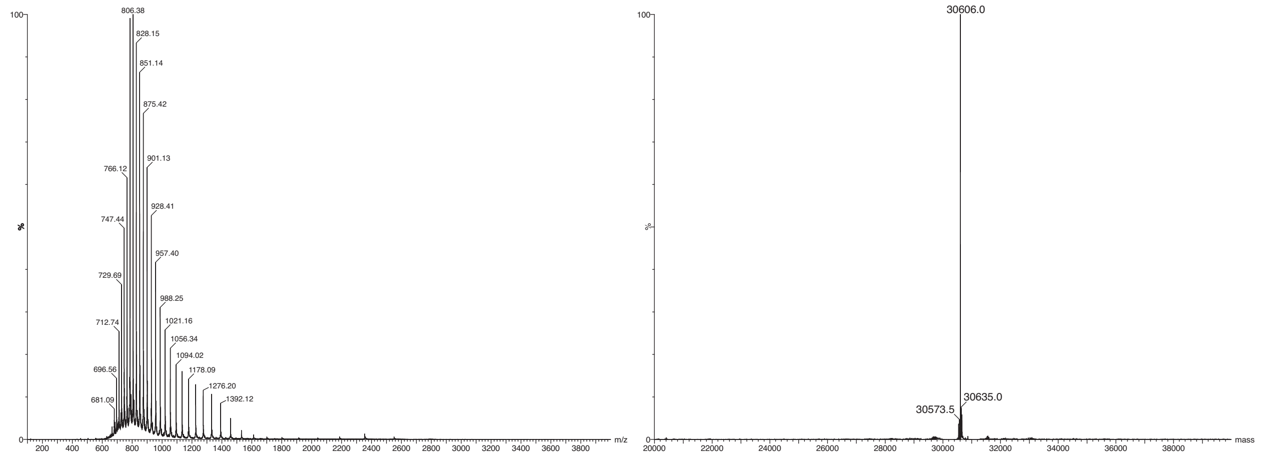

**ESI-MS characterization of protein conjugate sfGFP–pep1 [A]–D–COOH** Complex mass spectrum (left panel). Deconvoluted mass spectrum (right panel). Found: 30606.0, calculated for [$C_{1358}H_{2097}N_{367}O_{426}S_7$]: 30604.8.

Preparation of GFP– GGGKDLEGKGGS[KHL]GSGS[KHL]GGSKYPYDVPDYAKD–COOH

Protein was prepared as described in the general method.

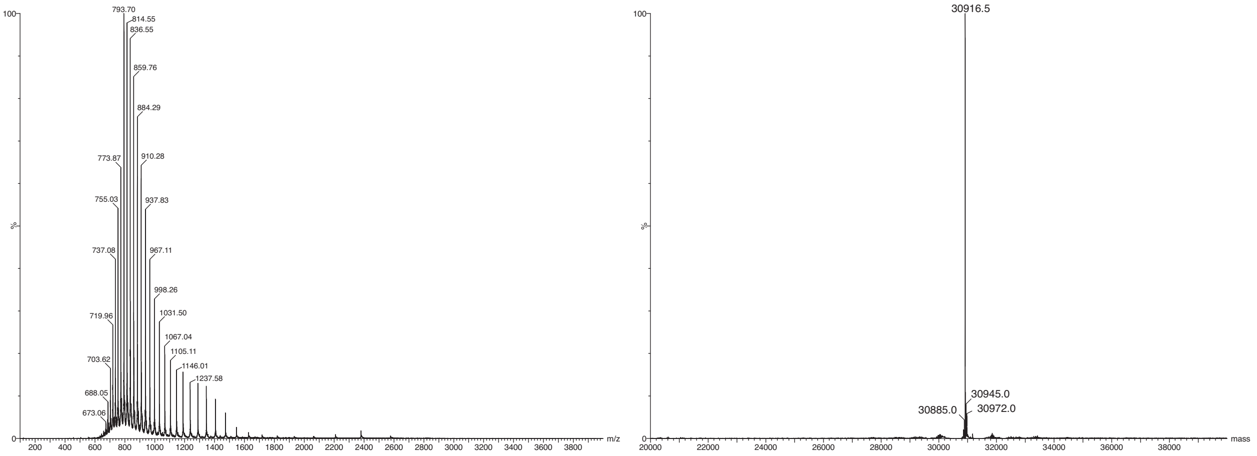

**ESI-MS characterization of protein conjugate sfGFP [KHL]** Complex mass spectrum (left panel). Deconvoluted mass spectrum (right panel). Found: 30916.5, calculated for [$C_{1376}H_{2131}N_{369}O_{428}S_7$]: 30915.2.

Preparation of GFP– GGGKDLEGKGGSAGSGSAGGSKYPYDVPDYAKS–COOH

Protein was prepared as described in the general method.

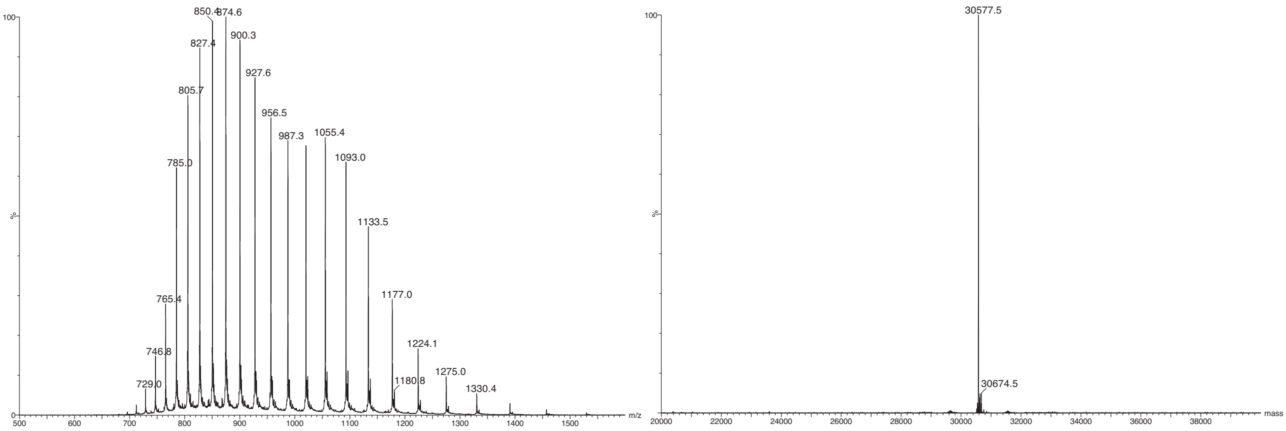

**ESI-MS characterization of protein conjugate sfGFP–pep1 [A]–S–COOH** Complex mass spectrum (left panel). Deconvoluted mass spectrum (right panel). Found: 30577.5, calculated for [$C_{1357}H_{2097}N_{367}O_{425}S_7$]: 30576.8.

Preparation of GFP– GGGKDLEGKGGSAGSGSAGGSKYPYDVPDYAKS–$CONH_2$

Protein was prepared as described in the general method.

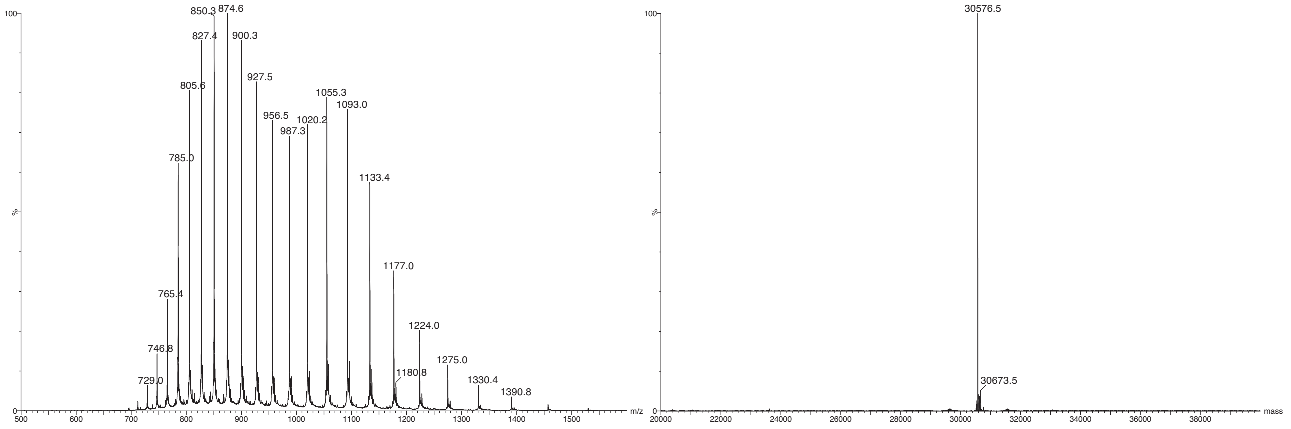

**ESI-MS characterization of protein conjugate sfGFP–pep1 [A]–S–$CONH_2$** Complex mass spectrum (left panel). Deconvoluted mass spectrum (right panel). Found: 30576.5, calculated for [$C_{1357}H_{2098}N_{368}O_{424}S_7$]: 30575.8.

Preparation of mCherry–GGGRRLEGKEEDEKGSRASDRFRGLR–COOH

Protein was prepared as described in the general method.

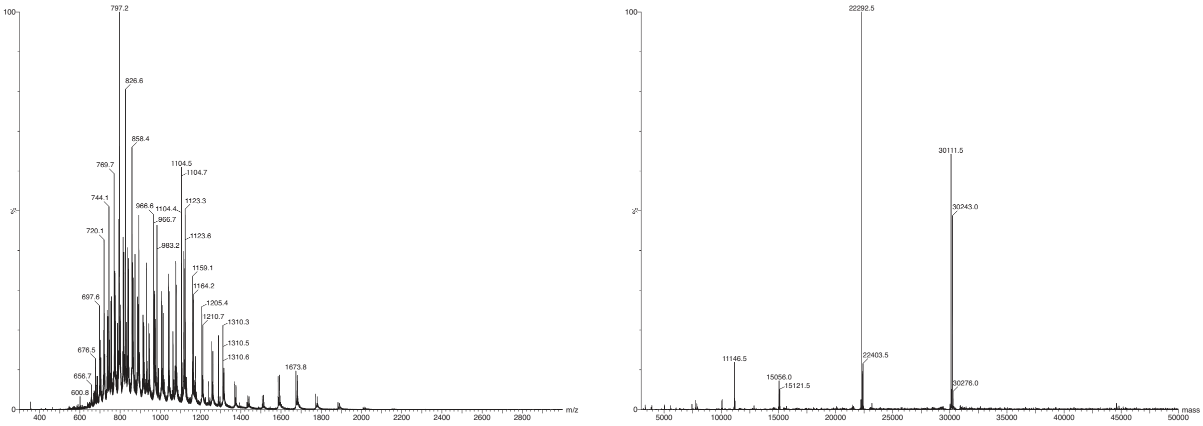

**ESI-MS characterization of protein conjugate mCherry–pep2 [RxxG]–R–COOH** Complex mass spectrum (left panel). Deconvoluted mass spectrum (right panel). A truncated species of mCherry with mass 22292.5 Da is observed. Initiator methionine processing is incomplete, providing two product mass peaks with 131.5 Da difference. Found: 30111.5, calculated for [$C_{1333}H_{2064}N_{366}O_{413}S_9$]: 30113.36.

Preparation of mCherry–GGGRRLEGKEEDEKGSRASDDFRDLR–COOH

Protein was prepared as described in the general method.

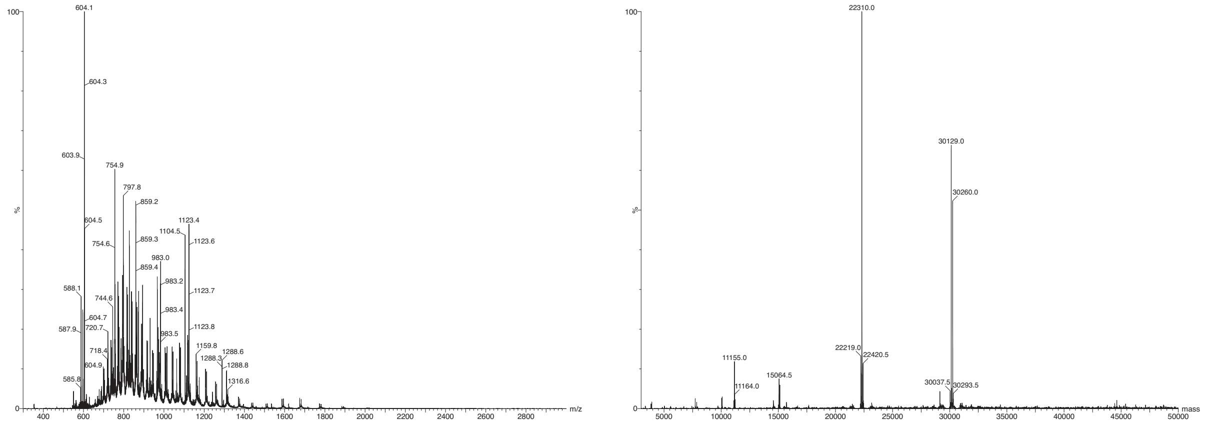

**ESI-MS characterization of protein conjugate mCherry–pep2 [RxxG]–R–COOH** Complex mass spectrum (left panel). Deconvoluted mass spectrum (right panel). A truncated species of mCherry with mass 22310.0 Da is observed. Initiator methionine processing is incomplete, providing two product mass peaks with 131 Da difference. Found: 30129.5, calculated for [$C_{1333}H_{2059}N_{363}O_{417}S_9$]: 30130.30.

Preparation of mCherry–GGGRRLEGKEEDEKGSRASDDFRDLR–$CONH_2$

Protein was prepared as described in the general method.

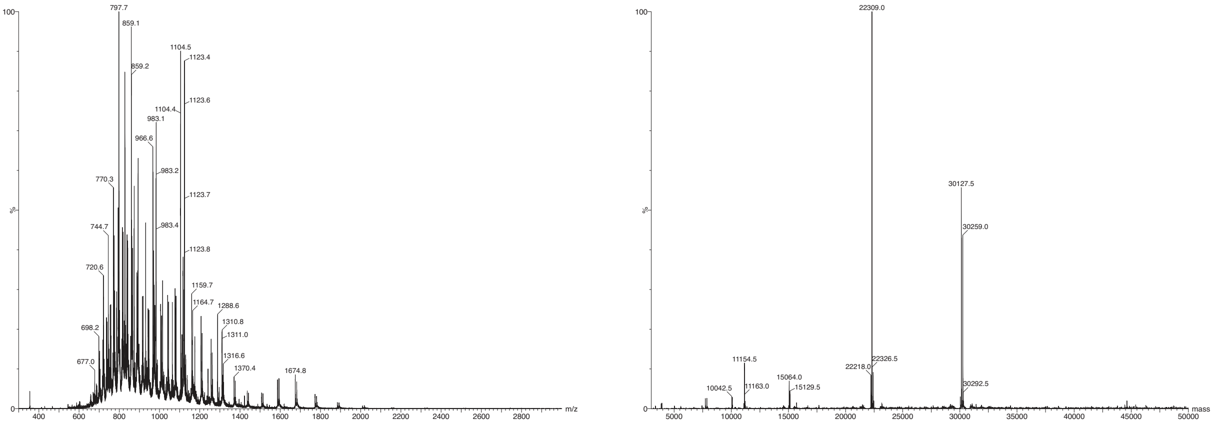

**ESI-MS characterization of protein conjugate mCherry–pep2 [DxxD]–R–$CONH_2$** Complex mass spectrum (left panel). Deconvoluted mass spectrum (right panel). A truncated species of mCherry with mass 22309.0 Da is observed. Initiator methionine processing is incomplete, providing two product mass peaks with 131.5 Da difference. Found: 30127.5, calculated for [$C_{1333}H_{2060}N_{364}O_{416}S_9$]: 30129.31.

Preparation of mTAGBFP2–GGGRRLEGKEEDEKGSRASDRFRGLR–COOH

Protein was prepared as described in the general method.

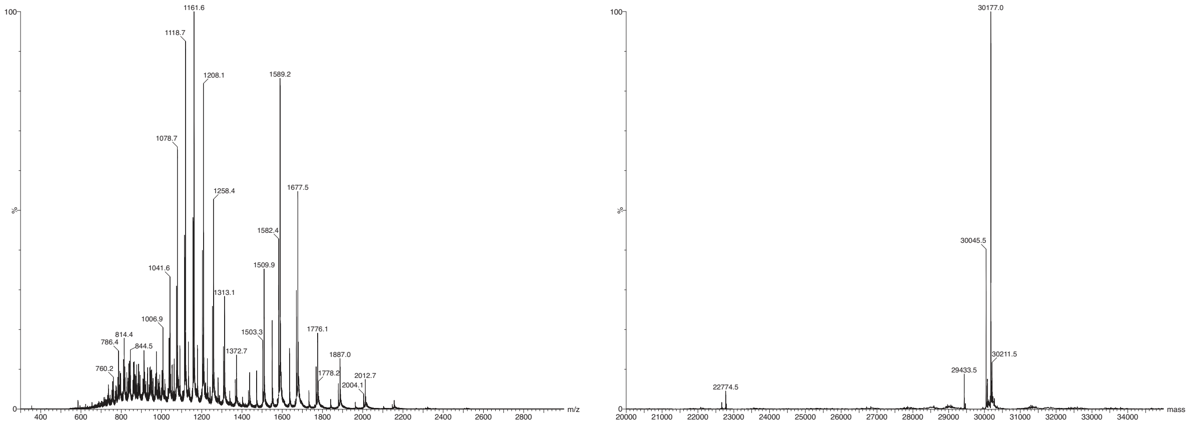

**ESI-MS characterization of protein conjugate mTAGBFP2–pep2 [RxxG]–R–COOH** Complex mass spectrum (left panel). Deconvoluted mass spectrum (right panel). Initiator methionine processing is incomplete, providing two product mass peaks with 131.5 Da difference. Found: 30177.0, calculated for [$C_{1336}H_{2081}N_{369}O_{407}S_{11}$]: 30176.68.

Preparation of mTAGBFP2–GGGRRLEGKEEDEKGSRASDDFRDLR–COOH

Protein was prepared as described in the general method.

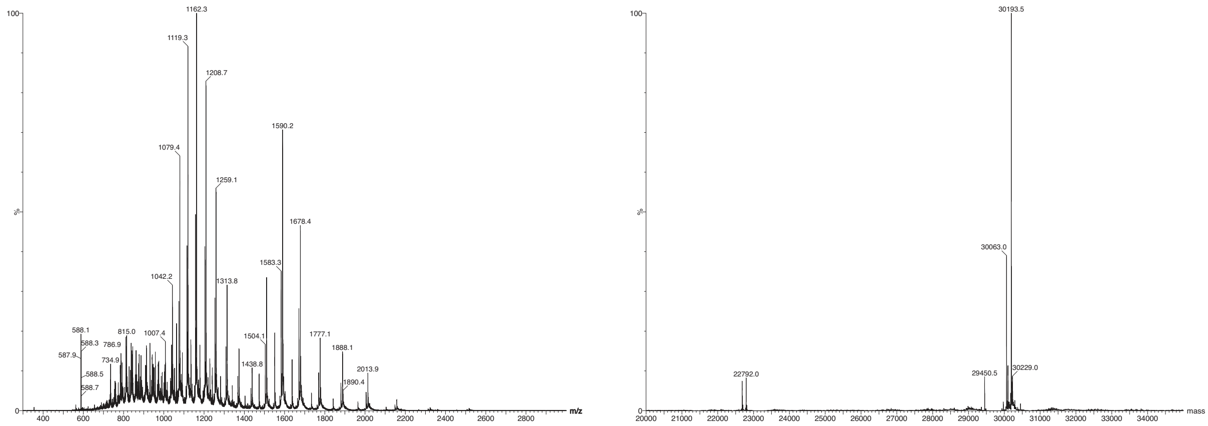

**ESI-MS characterization of protein conjugate mTAGBFP2–pep2 [RxxG]–R–CONH$_2$** Complex mass spectrum (left panel). Deconvoluted mass spectrum (right panel). Initiator methionine processing is incomplete, providing two product mass peaks with 130.5 Da difference. Found: 30193.5, calculated for [$C_{1336}H_{2076}N_{366}O_{411}S_{11}$]: 30193.62.

Preparation of mTAGBFP2– GGGRRLEGKEEDEKGSRASDDFRDLR–$CONH_2$

Protein was prepared as described in the general method.

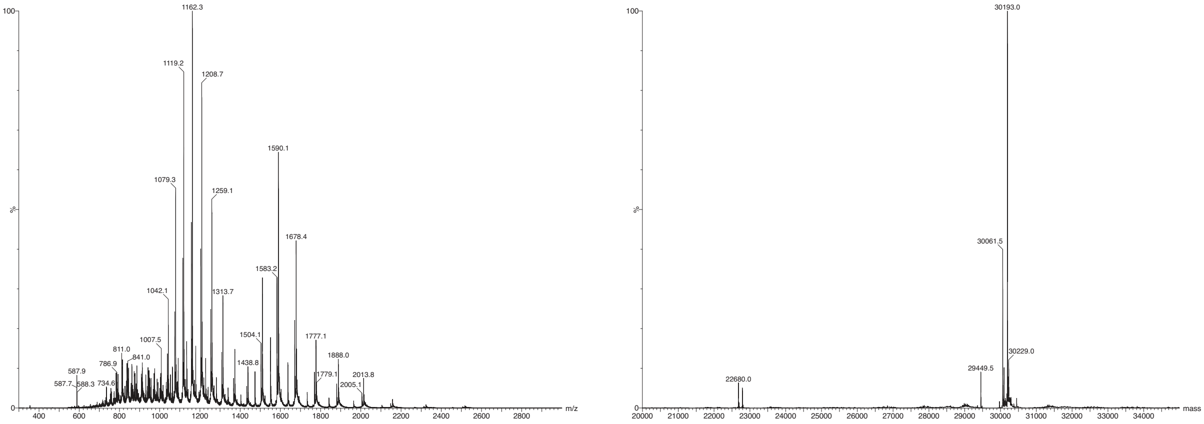

**ESI-MS characterization of protein conjugate mTAGBFP2–pep2 [DxxD]–R–$CONH_2$** Complex mass spectrum (left panel). Deconvoluted mass spectrum (right panel). Initiator methionine processing is incomplete, providing two product mass peaks with 130.5 Da difference. Found: 30193.0, calculated for [$C_{1336}H_{2077}N_{367}O_{410}S_{11}$]: 30192.62.

#### NMR spectra

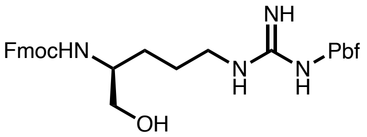

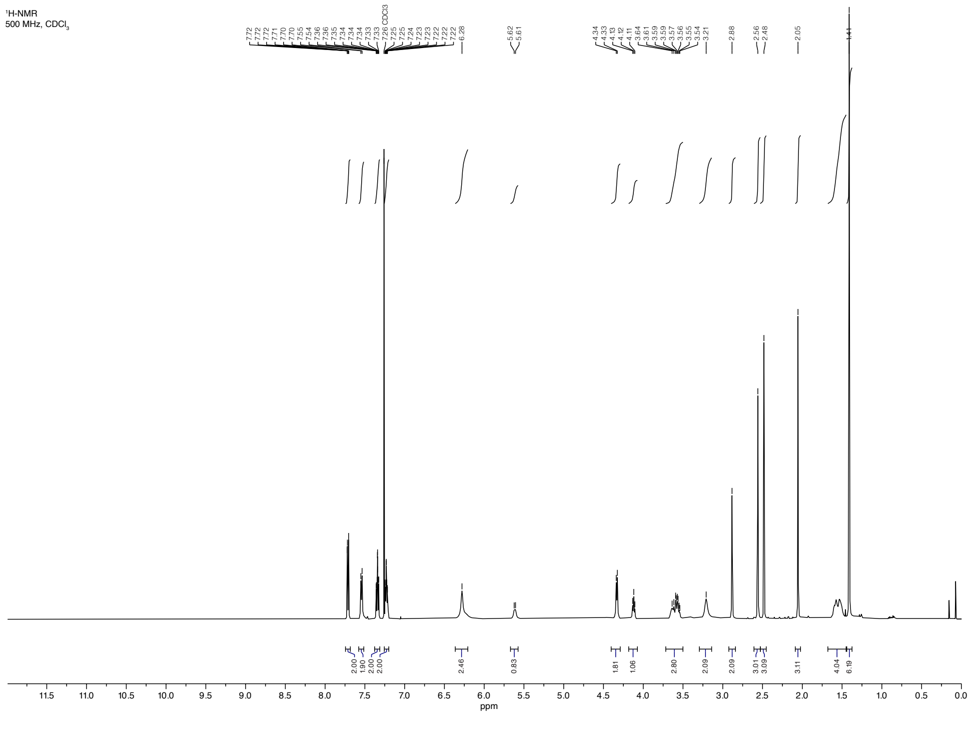

$^{13}C$-NMR
125 MHz, $CDCl_3$

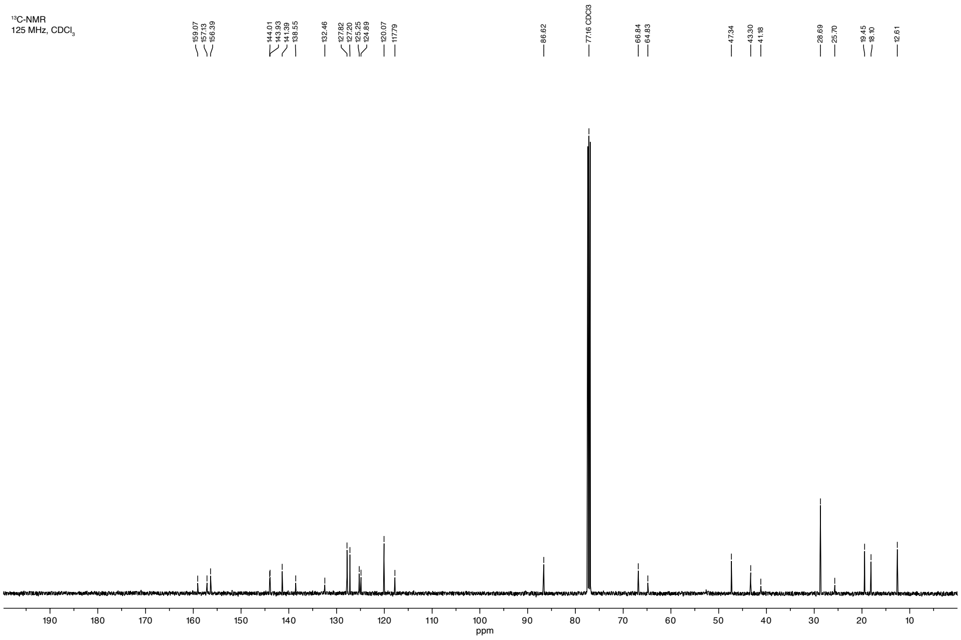
